# Poxviruses capture host genes by LINE-1 retrotransposition

**DOI:** 10.1101/2020.10.26.356402

**Authors:** Sarah M. Fixsen, Kelsey R. Cone, Stephen A. Goldstein, Thomas A. Sasani, Aaron R. Quinlan, Stefan Rothenburg, Nels C. Elde

**Affiliations:** Department of Human Genetics, University of Utah, Salt Lake City, USA; Department of Microbiology, University of California, Davis, USA

## Abstract

Horizontal gene transfer (HGT) provides a major source of genetic variation. Many viruses, including poxviruses, encode genes with crucial functions directly gained by gene transfer from hosts. The mechanism of transfer to poxvirus genomes is unknown. Using genome analysis and experimental screens of infected cells, we discovered a central role for Long Interspersed Nuclear Element-1 (LINE-1) retrotransposition in HGT to virus genomes. The process recapitulates processed pseudogene generation, but with host messenger RNA directed into virus genomes. Intriguingly, hallmark features of retrotransposition appear to favor virus adaption through rapid duplication of captured host genes on arrival. Our study reveals a previously unrecognized conduit of genetic traffic with fundamental implications for the evolution of many virus classes and their hosts.

**Summary:** Active selfish genetic elements in infected cells aid virus adaptation by catalyzing transfer of host genes to virus genomes.

## Introduction

Vertical inheritance of genetic material from parent to offspring can be augmented by horizontal gene transfer (HGT) (Keeling and Palmer, 2008). HGT can shortcut gradual mutational processes by facilitating adaptive leaps through single mutational events where genes can be transferred across kingdoms of life (Keeling and Palmer, 2008). HGT plays a central role in the diversification of many infectious microbes. This process is particularly well-described in bacteria, where HGT contributes to the emerging crisis of multi-antibiotic resistant pathogens (Wijayawardena et al., 2013). Many classes of viruses also acquire host genes by HGT (Caprari et al., 2015). A conspicuous example is Rous Sarcoma virus, in which recognition of the horizontal transfer of the host *c-Src* gene into the virus genome led to the discovery of oncogenes (Swanstrom et al., 1983). It has become increasingly clear that acquisition of host genes is a common mechanism by which viruses of many classes gain adaptive advantages to propagate and evolve. Despite the prevalence of HGT in diverse viruses and its importance in virus evolution, the mechanisms by which genes are transferred from host to virus genomes are not understood.

To investigate mechanisms of transfer we examined poxviruses, which encode a plethora of host genes gained by HGT. Based on phylogenetic analysis, more than 25% of poxvirus genes were acquired by HGT from hosts (Hughes and Friedman, 2005), although similarities not evident in sequence comparisons but revealed by shared protein structures suggest that a higher proportion of genes were acquired by horizontal transfer (Bahar et al., 2011). Many captured genes adapt to act as inhibitors of host immune defenses to favor virus replication (Elde and Malik, 2009). Some act as host range factors because their deletion leads to restriction of infectible cell types or species, implying that new acquisitions may aid in host jumps (Bratke et al., 2013; Haller et al., 2014). While the most notorious poxvirus, variola virus which caused smallpox, has been eradicated, other extant poxvirus strains are capable of infecting humans and pose the risk of new pandemics (Sklenovska and Van Ranst, 2018).

## Results

We hypothesized that examination of genes recently acquired by HGT might provide clues for revealing mechanisms of gene transfer. Virus homologs of the host Golgi Anti-Apoptotic Protein (GAAP; also called transmembrane Bax inhibitor motif protein family 4 or TMBIM4), found in several orthopoxviruses, are ~75% identical to human GAAP (Gubser et al., 2007; Saraiva et al., 2013), suggesting a recent acquisition and/or highly conserved function favoring virus replication. Strikingly, we observed nearly identical 21 base pair sequences flanking the open reading frame of virus GAAP (vGAAP), indicative of a target site duplication (TSD), consistent with gene capture resulting from LINE-1 mediated retrotransposition of host mRNA (Figs. 1A and S1). The vGAAP gene is situated near one of two identical inverted terminal repeat (ITR) regions of the virus genome, and on opposite ends of the genome in various viruses, suggesting that it was originally transferred into the ITR. A single copy of the putative 21bp TSD at an analogous position near the other ITR in the cowpox genome suggests a pre-integration sequence or pseudo empty site (Fig. 1A). Together with a published account of a non-autonomous Short Interspersed Nuclear Element (SINE) in the genome of taterapox virus (Piskurek and Okada, 2007), these observations suggest that host genes can be retrotransposed into poxvirus genomes by LINE-1 activity in a process akin to the generation of processed pseudogenes (Esnault et al., 2000).

**Fig. 1.**
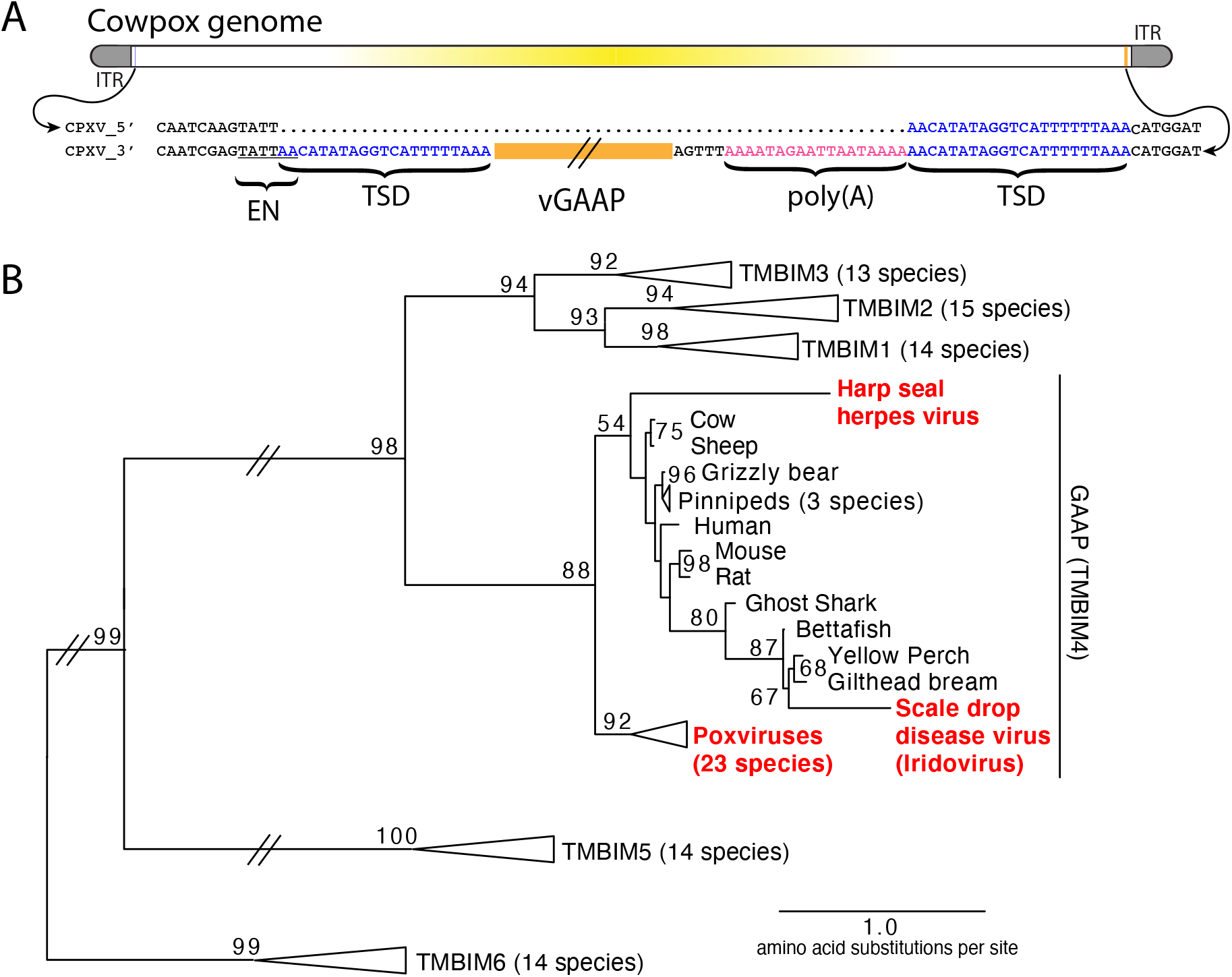
Retrotransposon-mediated transfer of GAAP into poxvirus genomes. (**A**) The poxvirus gene vGAAP (orange) is encoded near the inverted terminal repeat (ITR; gray) of cowpox genomes. Signatures of retrotransposition include TSDs (blue), a LINE-1-like endonuclease site (EN; underlined), and a partially degraded poly(A) tail (pink). Some cowpox genomes (CPXV_5’; top line) contain a pseudo-empty site, with a single copy of the TSD sequence. (**B**) Phylogeny of TMBIM proteins and virus-encoded TMBIM4s (red). Bootstrap values >50 are indicated.

Phylogenetic analysis revealed a single origin for the horizontal transfer of GAAP into poxvirus genomes. In addition, at least two other classes of DNA viruses, herpesviruses and iridoviruses, independently acquired GAAP by HGT (Fig. 1B) (Bellehumeur et al., 2016; de Groof et al., 2015). Another study revealed that the S1 gene of Squirrel Monkey cytomegalovirus encodes a protein more than 95% identical to squirrel monkey Signaling lymphocytic activation molecule family 6 (SLAMF6), followed by portions of the 3’ untranslated region (UTR) of the host mRNA (Perez-Carmona et al., 2015). These findings suggest that many DNA viruses acquire host genes by retrotransposition. However, the majority of host to virus gene transfers lack features of retrotransposition, which can quickly degrade through mutation, motivating our development of an experimental strategy for capturing HGT events in real time.

We developed a selection scheme using vaccinia virus (Copenhagen strain), which encodes nearly 200 genes in a 191,737 bp genome (See methods). We focused on K3L, a gene that was presumably acquired from a host by an ancient poxvirus based on sequence identity with human eukaryotic initiation factor 2α (eIF2α; 27.6% identical) (Beattie et al., 1991; Elde et al., 2009; Elde and Malik, 2009). K3L blocks the antiviral Protein Kinase R (PKR) pathway from arresting protein translation during viral infections by phosphorylating eIF2α (Fig. S2) (Beattie et al., 1991). Using a mutant virus deleted for K3L and another PKR inhibitor, E3L (Brennan et al., 2014), we infected a rabbit kidney (RK13) cell line expressing complementing chromosome-integrated copies of K3L fused to the gene encoding mCherry fluorescent protein (RK13-K3L) (Fig. 2A and see Methods for details). Viruses grown in RK13-K3L cells were transferred to a non-complementing baby hamster kidney (BHK) cell line to select for viruses that “reclaimed” the mCherry-K3L gene and/or somehow adapted in the absence of K3L to block host cell PKR and replicate.

**Fig. 2.**
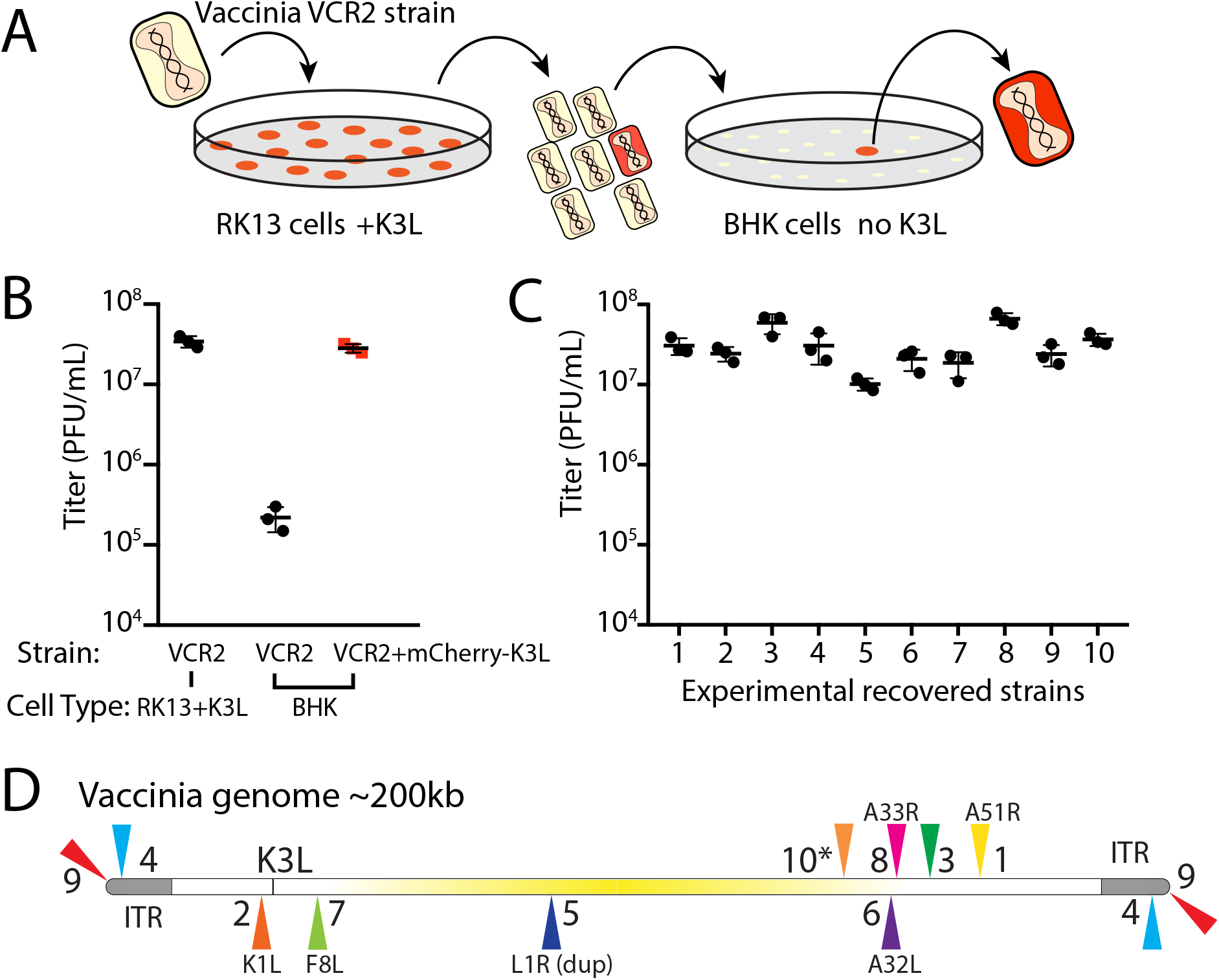
Experimental capture of K3L by horizontal gene transfer. (**A**) Viruses lacking K3L (VCR2) replicated in RK13 cells expressing mCherry-K3L and were transferred to BHK cells lacking K3L to select for capture of mCherry-K3L. (**B**) Replication of VCR2 in RK13-K3L cells and BHK cells compared to VCR2+mCherry-K3L (see Methods). (**C**) Replication of recovered isolates in BHK cells. Three biological replicates of each strain/isolate are shown in (B) and (C). (**D**) Vaccinia genome illustrating K3L integrations in recovered isolates indicated by colored triangles. Endogenous K3L location is shown. The central region of the genome is highlighted in yellow. Triangles above and below the genome denote positive and negative sense orientation respectively. Virus genes interrupted by K3L integrations are indicated. The asterisk denotes a genomic rearrangement in isolate 10 (Fig S3).

We screened ~500 million viruses over many experiments and recovered ten virus isolates from mCherry positive infected cell clones. In the restrictive BHK cells, viruses encoding K3L replicate 100-fold better than the parent virus deleted for K3L (Fig. 2B), providing strong selective pressure to screen for adapted viruses. After plaque purification of mCherry isolates, we infected BHK cells with each recovered virus isolate and observed titers similar to viruses encoding K3L (Fig. 2C). By sequencing the genome of each recovered virus isolate, we found that the mCherry-K3L fusion gene was integrated into the virus genome at a unique position in every isolate (Figs. 2D and S3), providing the opportunity to compare each independent integration event.

All integrations were consistent with a mechanism of LINE-1 retrotransposition-mediated HGT. The sequence of each transferred gene revealed hallmark features of mRNA gene capture, including 5’ and 3’ UTRs, precise intron splicing, and poly(A) tails (Fig. 3). For seven of ten isolates, TSDs flanked mCherry-K3L insertions and contained cut site sequences matching the consensus (TTTT/AA) of LINE-1 endonuclease (Fig. 3C). Although there were no TSDs flanking mCherry-K3L in isolates 5, 9, and 10, these integrations might result from a endonuclease-independent LINE-1 mechanism at DNA breaks (Morrish et al., 2002). Because our experiments lack any exogenous reverse transcriptase, we tested for endogenous activity in the RK13 cells and detected robust LINE-1 activity (Fig. S4). These results suggest that LINE-1 mediated retrotransposition may be a common mechanism of HGT into poxvirus genomes.

**Fig. 3.**
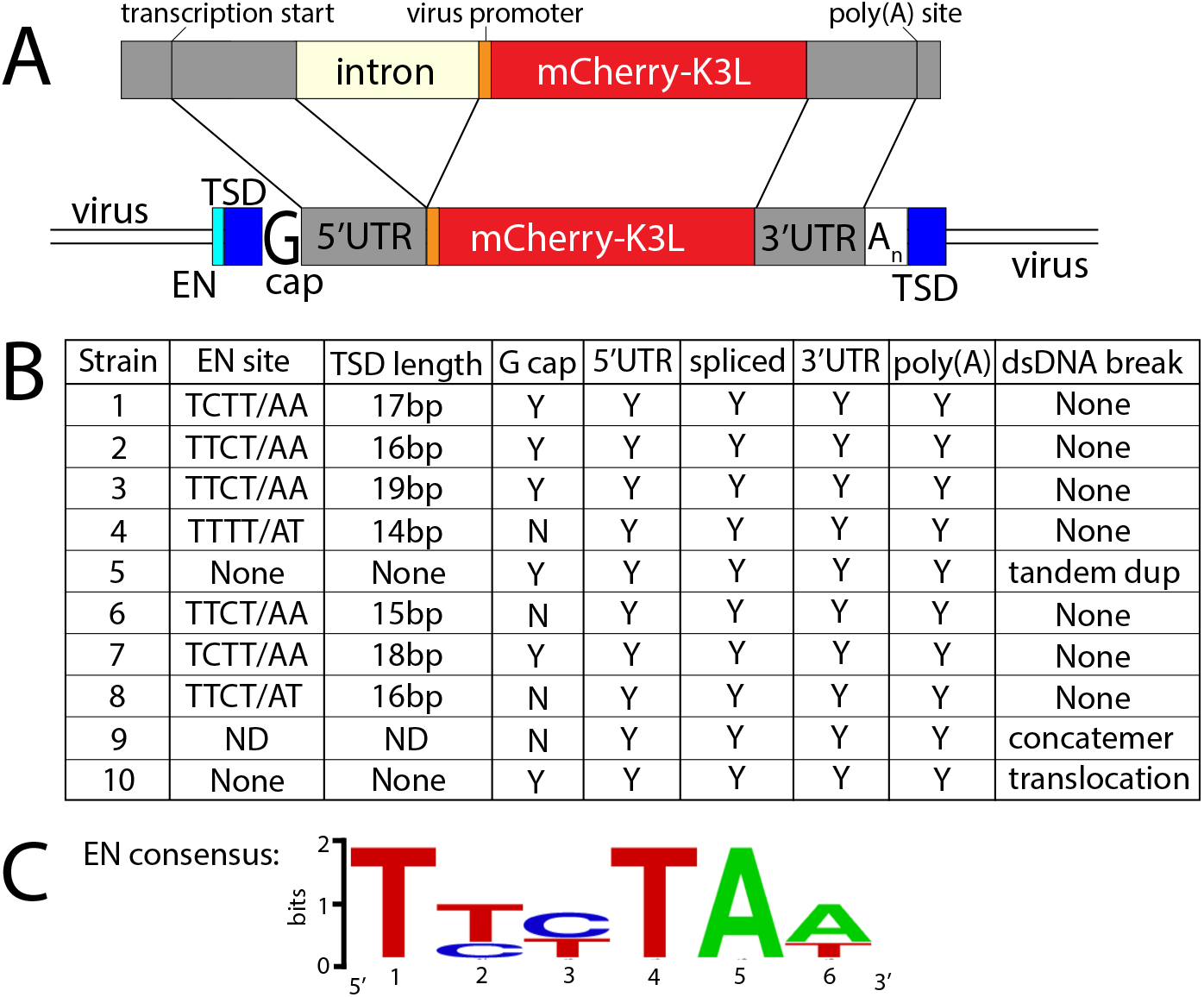
All virus isolates exhibit signatures of LINE-1 mediated retrotransposition. (**A**) Schematic of construct integrated in RK13-K3L cells (top) compared to K3L integrations in recovered viruses (bottom). Shared features include spliced introns, 5’ and 3’ untranslated regions (UTRs), and poly(A) tails (An). Seven isolates have target site duplications (TSDs, blue) with LINE-1 endonuclease cut sites (EN, cyan). Six isolates encode a guanine (G cap) adjacent to the 5’UTR, indicative of 7-methylguanylate mRNA capping. (**B**) Table highlights details of each isolate. (**C**) Logos plot of endonuclease cut site consensus sequence.

Poxvirus genomes are arranged with the essential genes, shared by all poxviruses, near the center of the genome. Clade- or species-specific genes, which more commonly exhibit evidence of HGT, are enriched near the ends of the genome (Lefkowitz et al., 2006; McLysaght et al., 2003). This pattern may reflect that gene transfer into the central region of genome is more likely to interrupt genes vital for virus replication (Lefkowitz et al., 2006). Consistent with this, all but one recovered isolate integrated K3L outside a roughly defined central region of the genome containing 90 core chordopoxvirus genes (Lefkowitz et al., 2006) (Fig. 2C). In the case of isolate 5, the K3L integration involved a duplication of two essential genes, G9R and L1R (Fig. S3). Because gene duplications can appear frequently in poxvirus genomes (Hughes and Friedman, 2005), this activity might enable the persistence of horizontally transferred genes by preserving existing ones, consistent with a process described in a contemporaneous study (Rahman *et. al.*, 2020).

Previous studies found that duplication of poxvirus genes, followed by rapid gene copy amplification, can boost virus replication (Brennan et al., 2014; Elde et al., 2012). In our analysis, we discovered two isolates where K3L had undergone polymorphic copy number increases (isolates 4 and 5; Fig. 4A). Homologous recombination, including unequal crossovers leading to duplications, requires as little as 16bp of homology in poxviruses (Yao and Evans, 2001), suggesting that LINE-1 induced TSDs might facilitate gene amplification. Genes transferred to within the ITR are duplicated on the other end by replication-based mechanisms (McFadden and Dales, 1979), as we observed in isolates 4 and 9 (Fig. 2C). Genes integrated into the near the termini of the genome by HGT, like isolate 4, may be especially prone to duplication due to stretches of short, high copy tandem repeats found in and near the inverted terminal repeat regions (Fig. 4A). HGT therefore likely occurs evenly across poxvirus genomes, but with an enrichment of transferred genes persisting near the ends, where insults to essential genes are less probable and duplication-based adaptation is favored (Fig. S5). Consistent with this idea, gene families found only in specific lineages of poxviruses, indicating more recent acquisition, are more likely to be present in multiple copies than more host genes acquired in (Hughes and Friedman, 2005).

**Fig. 4:**
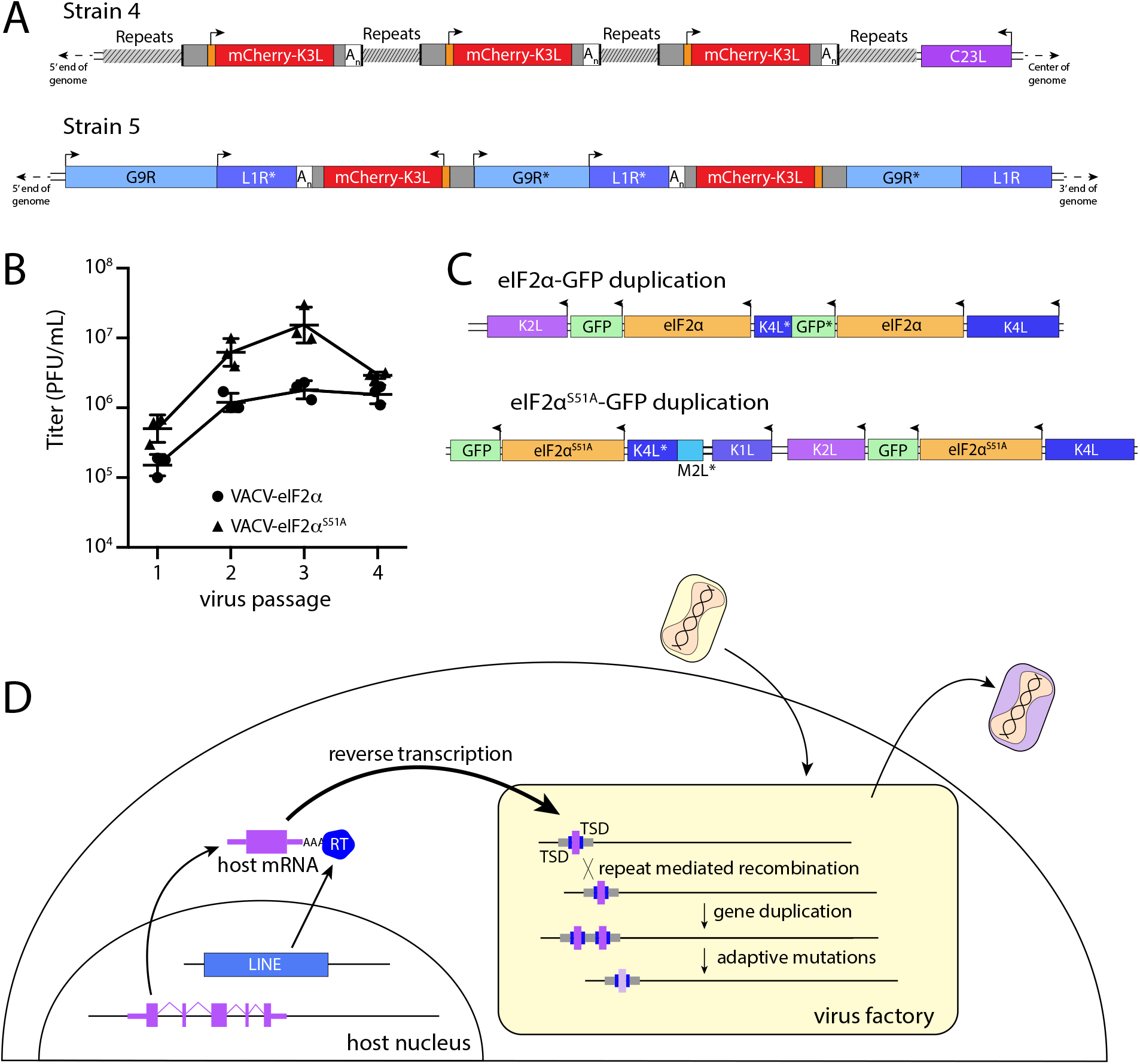
Homology driven gene duplication of K3L and eIF2α. (**A**) Schematics of mCherry-K3L duplications in isolates 4 and 5. (**B**) Virus titers after serial infections of eIF2α viruses in HeLa cells (see Methods). (**C**) Schematics of eIF2α duplications following virus passaging. Asterisks denote gene truncations (**D**) Model diagram highlighting how host genes (purple) are retrotransposed by LINE-1 reverse transcriptase (RT, blue), into virus genomes (yellow box). Following horizontal transfer, gene duplidation is facilitated by unequal crossover recombination between TSDs (blue) or flanking repeat sequence (gray).

While our experimental system revealed LINE-1 driven mechanisms of HGT, the scheme involved selection for viruses regaining an existing virus gene. To better model virus adaptation following acquisition of a host gene, we engineered viruses with the human eIF2α gene in place of K3L (Fig. S6; see Methods). We performed four serial infections of viruses expressing eIF2α or an eIF2α variant lacking the PKR phosphorylation site (S51A) in RK13 cells. While the recombinant viruses replicate weakly, we observed a nearly 10-fold increase in virus replication for both eIF2α-encoding viruses after passaging (Fig. 4B). By sequencing the eIF2α region in evolved virus strains, we found independent gene copy increases of eIF2α in both viruses (Fig. 4C). These results support a critical role for gene copy increases following HGT in poxvirus evolution, as also observed for K3L and other genes (Brennan et al., 2014; Elde et al., 2012).

## Discussion

Horizontal gene transfer is an important process facilitating virus evolution (Caprari et al., 2015; Deeg et al., 2018; Hughes and Friedman, 2005; Koonin and Yutin, 2019; McLysaght et al., 2003). Our discovery of LINE-1 retrotransposition-mediated HGT from host to poxvirus genomes has several notable implications. In addition to self-propagation in “host” genomes, retrotransposition to virus genomes could greatly extend the range of LINE-1 or related elements that subsequently mobilize to a newly infected virus host (Piskurek and Okada, 2007). The extensive host range of some poxviruses, for example between rodent and primate species, suggests the possibility of cross host lineage transfer of LINE-1s, non-autonomous Alus, and host transcripts between mammals. A process involving the frequent appearance of DNA transposons in some insect viruses suggests a parallel mechanism for the spread of transposable elements between species (Gilbert et al., 2016). In cases of retrotransposition of host transcripts, the resulting processed pseudogene products are enriched for highly expressed genes, including ribosomal genes (Zhang et al., 2004). The over representation of certain transcripts might provide clues for understanding and predicting how viruses might adapt with repurposed genes mimicking host functions (Elde and Malik, 2009). Currently recognized cases of HGT involving oncogenes and regulators of translation, like eIF2α, provide cases of transcripts exhibiting high gene expression and cellular functions easily repurposed to promote virus replication.

Repetitive target site duplications generated by LINE-1 activity may also enhance poxvirus adaptation by facilitating rapid gene copy number amplification of horizontally transferred genes on arrival. Increases in gene copy number augment the probability of beneficial mutations appearing, allowing for swift diversification of newly acquired genes (Elde et al., 2012). Thus, form follows function as the repetitive termini of poxvirus genomes further enable the emergence and persistence of acquired genes predominately at the ends of the genome (Fig. 4D). As beneficial virus gene variants endure, they end up closer to the center of the genome as subsequently captured genes appear near the ends.

Similar mechanisms of virus adaptation involving HGT may be shared by other diverse classes of nucleocytoplasmic DNA viruses (NCLDVs), in addition to poxviruses (Deeg et al., 2018; Koonin and Yutin, 2019). Many giant viruses, which can exceed a megabase in genome size and infect diverse host species, encode a variety of genes acquired by HGT, including ribosomal genes. Given a primary role for LINE-1 elements in facilitating gene transfer, we speculate that other classes of transposable elements may spur NCDLV evolution by mobilizing host genes into virus genomes across diverse ecosystems.

## Materials and methods

### Phylogenetic analysis and sequence comparison of vGAAP

TMBIM sequences for various species were downloaded from NCBI (Supplemental data) and aligned using the COBALT Multiple Alignment Tool. Amino acid Phylip alignments (see supplemental materials) were analyzed by PhyML (http://www.atgc-montpellier.fr/phyml/) for tree building with 100 bootstraps (Guindon et al., 2010).

### Cell culture

BHK and HeLa cells were cultured in Dulbecco minimum essential medium (DMEM; HyClone) supplemented with 10% FBS, 2 mM L-glutamine and 100 μg/mL of pen/strep. RK13 cells were cultured in MEM-alpha supplemented with 10% fetal bovine serum (FBS; Gibco), 2 mM L-glutamine (HyClone) and 100 μg/mL of penicillin-streptomycin (pen/strep; HyClone). RK13 cells expressing SLP-mCherry-K3L (RK13-K3L) were generated by transfecting 3×10^5^ RK13-E3K3 cells (Brennan et al., 2014) with 2.5ug piggybac vector (see supplemental sequences) and 0.5 ug transposase vector (gift from Ed Grow, University of Utah) with 9uL Fugene-HD. Cells with the piggybac construct integrated were selected by the presence of 1-2 ug/mL puromycin, and populations were monitored for mCherry expression. After two weeks of selection, remaining polyclonal cells were propagated in MEM-alpha media supplemented with 10% FBS, 2 mM L-glutamine and 100 μg/mL of pen/strep. All cultures were maintained and infections performed at 37°C in a humidified 5% CO2 incubator.

### Virus strains

VC-2 refers to the Vaccinia virus-Copenhagen strain (Goebel et al., 1990) and VCR2 is a genetically modified isolate of VC-2 deleted for E3L and K3L (Brennan et al., 2014). To generate VCR2+mCherry-K3L, we cloned pBlue_165_mCherry-K3L (see supplemental sequences) into the MCS of pBluescriptIIKS (-) (Addgene). This plasmid was transfected into BHK cells, which were then infected with VCR2 virus. Recombinant viruses were plaque purified in BHK cells and checked for purity by PCR amplifying across the mCherry-K3L insertion site. The amplicon was gel purified and sequenced.

To make eIF2α viruses, sequences flanking K3L from VACV were amplified from VC-2 viral DNA: 680bp of 5’ homologous sequence was amplified with the primers K2Lflank_F (5’-CTTCTTATC GATTTTTTATACCGAACATAAAAATAAGGTTAATTA) and K2Lflank_R (5’-CTTCTTCATATGG TGATTGTATTTCCTTGCAATTTAG), and 1024bp of 3’ homologous sequence (including the native K3L promoter) was amplified with the primers K4Lflank_F (5’-CGTCGTGCGGCCGCCTTGTTAACGGGCTCGTAAATT) and K4Lflank_R (5’-CGA GCGGAGCTCGTACGATACATAGATATTACAAATATCCTAG). A VACV synthetic early/late promoter (SLP)(Chakrabarti et al., 1997) was created by annealing primers SLP_F (5’ TCGACAATTGGATCAGCTTTTTTTTTTTTTTTTTTGGCATATAAATA AGAAGCTTCCCGGGTCTAGAC) and SLP_R (5’-AGCTCAGATCTGGGCCCTTCG AAGAATAAATATACGGTTTTTTTTTTTTTTTTTTCGACTAGGTTAAC). EGFP was amplified from pN1-EGFP (Addgene) using primers EGFP_F (5’-GGAGGACTCGAGATGGTGAGCAAGGGCGA) and EGFP_R (5’-GGAGGTATCGA TTTACTTGTACAGCTCGTCCATGC). Each PCR product was digested with restriction enzymes (New England Biolabs), gel purified (Zymo Research), and sequentially cloned into pB.2 as follows: 5’ flank with ClaI and NdeI, 3’ flank with NotI and SacI, SLP with SalI and XhoI, and EGFP with XhoI and ClaI. The resulting plasmid contained EGFP following SLP, between the two K3L flanking sequences (pB.2-EGFP). The K3L open reading frame was amplified from VC-2 viral DNA using primers K3L_F (5’-GTTGTAG GATCCATGCTTGCATTTTGTTATTCGTTGC) and K3L_R (5’-GTTCTTGTCGACTT ATTGATGTCTACACATCCTTTTG). The eIF2α gene was amplified from human cDNA using primers eIF2α_F (5’-GATGTAGGATCCATGCCGGGTCTAAGTTGTAGAT) and eIF2α_R (5’-CTACTTGTCGACTTAATCTTCAGCTTTGGCTTCCAT). The resulting PCR products were cloned into pB.2-EGFP with BamHI and SalI, placing K3L or eIF2α immediately following the native K3L promoter, and upstream of SLP-EGFP to create pB.2-K3L and pB.2-eIF2α, respectively. Site-directed mutagenesis was performed on pB.2-eIF2α or pB.2-eIF2α-ΔC using primers eIF2α_S51A_F (5’-GATAC GCCTTCTGGCTAATTCACTAAGAAGAATCATGCCTTC) and eIF2α_S51A_R (5’-GAAGGCATGATTCTTCTTAGTGAATTAGCCAGAAGGCGTATC) to generate pB.2-eIF2α-S51A.

Recombinant eIF2α-encoding viruses were constructed by replacing the K3L gene using homologous recombination. RK13-E3K3 cells were infected with VCR2 (MOI = 1.0) and transfected at 1-hour post-infection with pB.2-EGFP, pB.2-K3L, pB.2-eIF2α, or pB.2-eIF2α-S51A plasmids by use of FuGENE6 (Promega) according to the manufacturer’s protocol. Infected cells were collected at 48 hours post infection, and viruses were released by one freeze-thaw cycle followed by sonication. Resulting viruses were plaque purified in RK13-E3K3 cells four times, selecting for recombinants expressing EGFP. Final virus clones were verified by PCR and sequencing of viral DNA across the K2L-K4L region of the genome.

### Experimental screen for horizontal gene transfer events

Confluent 150mm dishes of RK13-K3L cells were infected with VCR2 at a MOI of 0.1. Cell-associated virus was collected after 48 hours of infection, and released from the cell by one freeze-thaw cycle followed by sonication as described (Isaacs, 2004). Confluent 150mm dishes of BHK cells were then infected with ~10^6^ PFUs of this virus stock for 7 days, during which they were monitored for mCherry expression. 500 plates were screened, which accounts for an estimated 500 million viruses. For mCherry positive clones observed by fluorescence microscopy, cell-associated virus was collected and released by one freeze-thaw cycle followed by sonication. mCherry-expressing virus was then plaque purified in BHK cells as described (Isaacs, 2004).

### Analysis of horizontal gene transfer events by inverse PCR (iPCR)

Virus DNA was extracted from mCherry-expressing clones as previously described from infected BHK cells (Esposito et al., 1981). Purified DNA was digested with XbaI and SpeI or BglII (New England Biolabs (NEB)), diluted, and ligated (Quick ligase; NEB) to circularize linear fragments. Primers pointing away from each other in both mCherry (F-CGTGGAACAGTACGAACGCG and R-CCATGTTATCCTCCTCGCCC) and K3L (F-GAGCATAATCCTTCTCGTATACTC and R-GAATATAGGGATAAACTGGTAGGG) were used to amplify regions flanking mCherry-K3L insertions. Resulting PCR bands were gel purified and Sanger sequenced with the iPCR primers. From this sequencing, the general location of the mCherry-K3L insert could be inferred. Primers flanking this region were then used to amplify the putative insertion, along with flanking sequences. These amplicons were then gel purified, Sanger sequenced, and compared to the parent (VCR2) genome to characterize gene integrations. Because isolates 7 and 9 incorporated mCherry-K3L within the repetitive ITR, each end was PCR amplified separately using unique flanking primers and internal primers (mCherry-R and K3L-F, above). However, only the 5’ end of the insertion of isolate 9 was amplified, despite multiple attempts. Thus, we cannot be sure whether this isolate includes a TSD or was cut at a LINE-1 endonuclease site.

### Long read genome sequencing of virus isolates

Virus particles from plaques expressing mCherry-K3L were isolated from BHK cells and virus cores were purified by ultracentrifugation through a 36% sucrose cushion at 60k rcf for 1 hr. Virus DNA was extracted as previously described (Esposito et al., 1981). The SQK-LSK108 library kit (Oxford Nanopore Technologies) was used to prep isolate 1 DNA, which was sequenced on a FLO-MIN106 cell. A SQK-RBK001 kit was used to prep DNA from isolates 2-5, which were multiplexed on a FLO-MIN107 cell. Isolates 6-10 were also multiplexed using the SQK-RBK004 library prep kit, and sequenced on FLO-MIN107. Reads were base-called with the Oxford Nanopore Albacore program and aligned to a reference genome that included the VCR2 genome on one contig and the cellularly-expressed mCherry-K3L construct on a separate contig using default NGMLR parameters (Sedlazeck et al., 2018). Integrative Genomics Viewer (IGV) software (Broad Institute) was used to visualize insertions (Robinson et al., 2011). Sequencing data is deposited in the NCBI SRA database, accession number: PRJNA614958.

### Measuring titers of virus strains and isolates

We plated 5×10^6^ cells in 100mm dishes and infected them 16 hours later at an MOI of 0.1. Three plates (biological replicates) were infected with each isolate or strain. After 48 hours, media was aspirated, and cell-associated virus stock was collected by one freeze-thaw and sonication cycle. Then, 6-well plates were seeded with 5×10^5^ BHK cells per well, and, 16 hours later, infected with 10-fold serial dilutions of virus stock (2 wells per dilution) in 200uL media. After 2 hours at 37°C, 2mL of media was added to each well. Cells were fixed and stained with 20% Methanol + 0.2% crystal violet 48 hours post infection. Wells with 10-100 plaques were counted and averaged to calculate virus titer. Each of the biological replicates is shown in Figure 2 and Figure 4 along with an average of the three and calculated standard deviations.

### Alu assay of RK13 cells

5×10^5^ HA-HeLa or RK13 cells were plated per 100mm dish (3x each cell type). 16 hours later, each plate was transfected with 5ug Alu-Neo (Dewannieux et al., 2003). After 2 days, all cells were treated with 2ug/mL G418. Media was replaced daily for 7 days, after which time it was removed, and cells were fixed and stained with 20% Methanol + 0.2% crystal violet.

### Detection of duplication events

Single Nanopore reads with multiple tandem copies of the mCherry-K3L fusion gene were analyzed using IGV (Robinson et al., 2011). Duplications were confirmed by PCR, using the inverse PCR primers in mCherry (F-CGTGGAACAGTACGAACGCG and R-CCATGTTATCCTCCTCGCCC). PCR amplicons were then gel purified and Sanger sequenced to determine break points. However, due to the repetitive nature of the sequence flanking the isolate 7 insertion, the exact breakpoint was not able to be determined.

### Serial passage of VACV-eIF2α virus strains

For each passage, 150-mm dishes were seeded with an aliquot from the same stock RK13-cells (5 x 10^6^ cells/dish). For P1, dishes were infected with eIF2α-GFP or eIF2α^S51A^-GFP virus (MOI = 1.0) for 2 hours in 5mL and then supplemented with 15mL medium. After 48 hours, cells were washed, pelleted, and resuspended in 1mL of medium. Virus was released by one freeze-thaw cycle followed by sonication. 900μl of virus was then used to infect a new dish of cells for P2, and the process was repeated for subsequent passages. Viral titers were determined using the remaining 100μl of reserved virus stocks from each passage by 48-hour plaque assay in RK13-E3K3 cells.

### Analysis of viral-encoded eIF2α genes

RK13-E3K3 cells were infected with P4 eIF2α-GFP or eIF2α^S51A^-GFP viruses (MOI = 0.1) for 24 hours. Virus-infected cells were collected, and total viral DNA extracted as previously described(Esposito et al., 1981). The region between K2L and K4L containing the different viral-encoded eIF2α genes was amplified by PCR with primers K2L_seq_F and K4L_seq_R (5’-GGCATTGGTAAATCCTTGCAGA and 5’-CACCTTTTAGTAGGACTAGTATCGTACAA, respectively). SNV detection was performed by sequencing across eIF2α using primers eIF2α_F and eIF2α_R for eIF2α-GFP and eIF2α^S51A^-GFP PCR products, CNV analysis was performed by PCR using primers eIF2α_rep_F (5’-CCTCCTATGGAAGCCAAAGCTGAAGATGAA) and eIF2α_rep_R (5’-CCTCCTATCTACAACTTAGACC CGGCAT) for eIF2α-GFP and eIF2α^S51A^-GFP viral DNA. Any CNV PCR products formed were sequenced using the same primers for breakpoint detection.

## Acknowledgments

We thank D. Hancks, E. Choung, and E. Grow for reagents and advice.

## Author contributions

N.C.E. and S.R. conceptualized the project. N.C.E., S.M.F. and S.R. contributed to experimental design. S.M.F. and K.R.C. performed experiments. S.M.F, K.R.C., and T.A.S. analyzed and validated data. S.M.F., S.A.G. and N.C.E. visualized data. N.C.E. supervised. S.M.F wrote the original draft and N.C.E., S.M.F., and S.A.G. reviewed and edited the manuscript.

## Data and materials availability

Bioproject PRJNA614958

## Supplemental Materials

Figures S1-S6

Table S1-S2

Supplementary sequences

---

**Fig. S1.**
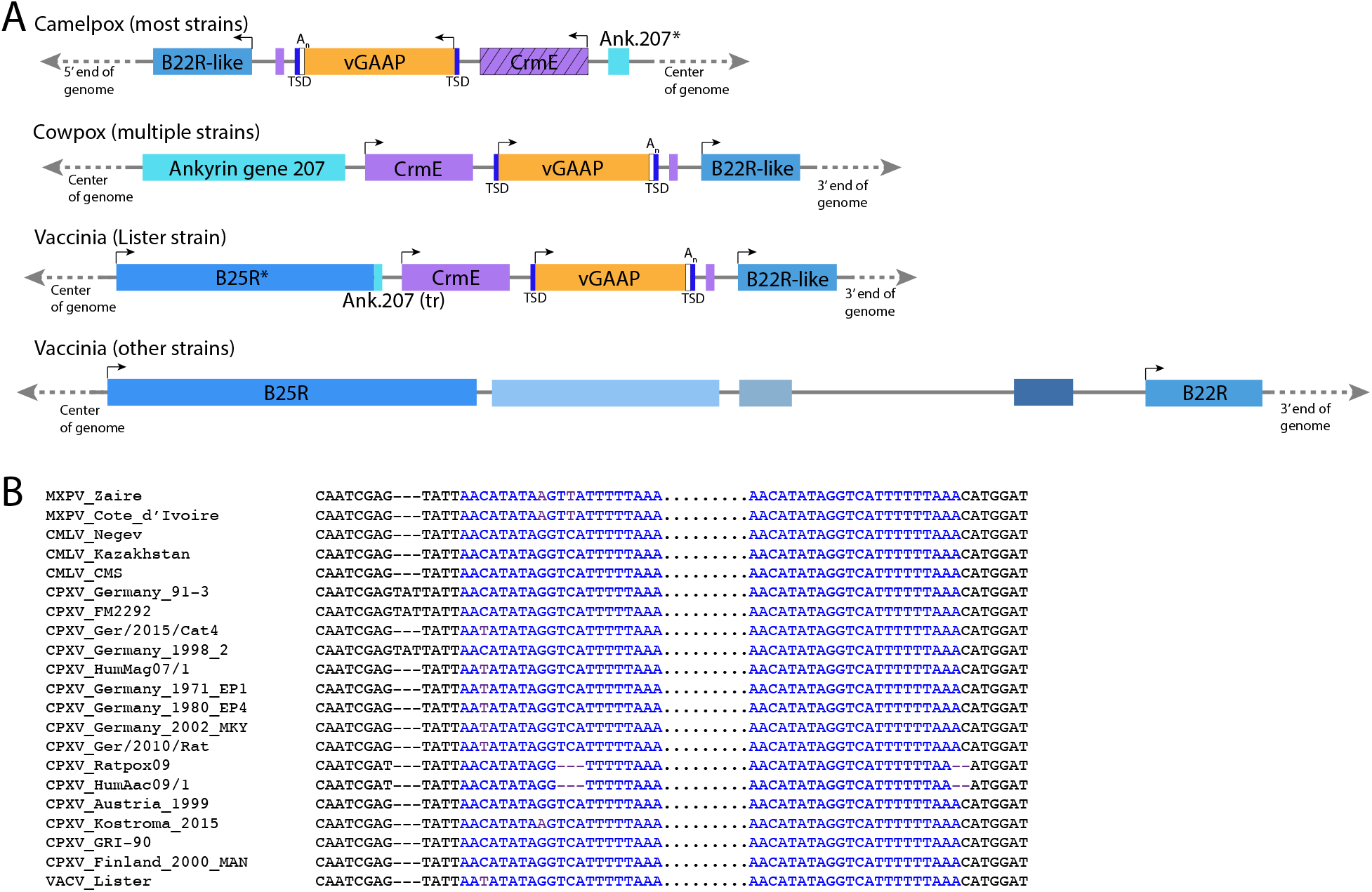
(A) v-GAAP (orange bar) is encoded by many orthopox species, including most camelpox strains, many cowpox strains, and at least one vaccinia strain. In camelpox viruses, v-GAAP is encoded on the left end of the genome; in all other viruses, v-GAAP is on the right end. In all viruses, v-GAAP is flanked by CrmE and vaccinia-B22R. In vaccinia strains that do not encode v-GAAP, CrmE is also missing, suggesting recombination of the region between B22R and B25R, which is truncated in the Lister strain. (B) Alignment of target site duplications (blue) and flanking sequence in 21 v-GAAP-encoding poxvirus strains. MPXV, monkeypox; CMLV, camelpox; CPXV, cowpox; VACV, vaccinia. Asterisks denote gene truncations.

**Fig. S2.**
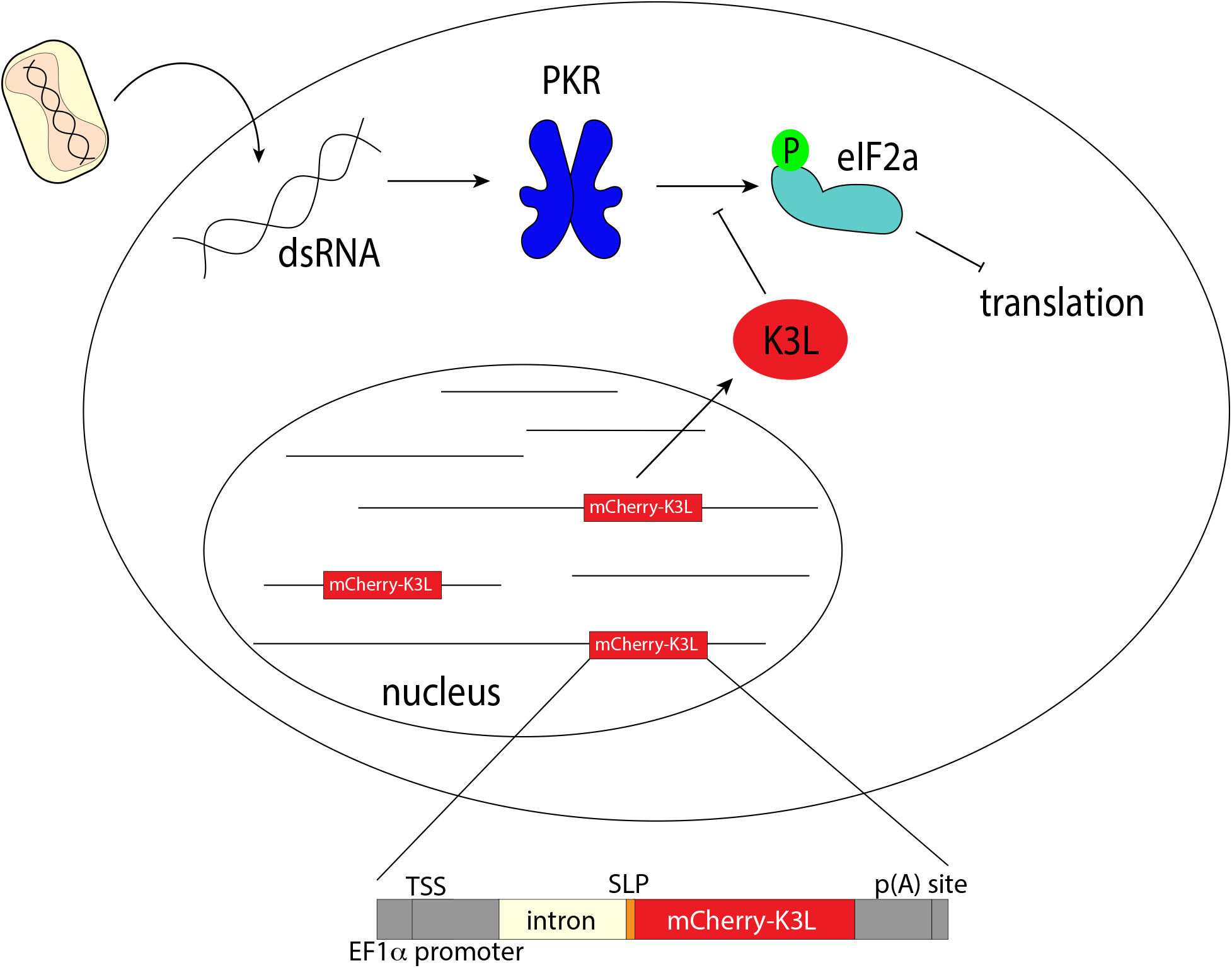
Protein Kinase R (PKR) is activated by binding dsRNA which is made in the cytoplasm during the course of a viral infection. When activated, PKR binds and phosphorylates the eukaryotic initiation factor, eIF2α, leading to a block in translation and preventing viral replication. K3L acts as a pseudo-substrate of PKR, preventing activated PKR from binding eIF2α, which is then free to initiate translation. In the complementing RK13-K3L cells, an mCherry-K3L fusion gene was integrated into cellular chromosomes. The cellular mCherry-K3L expression construct is shown below. The fusion gene is driven by a strong EF1a promoter, but the construct also includes a poxvirus-specific promoter, the SLP, so that mCherry-K3L could be immediately expressed if transferred.

**Fig. S3.**
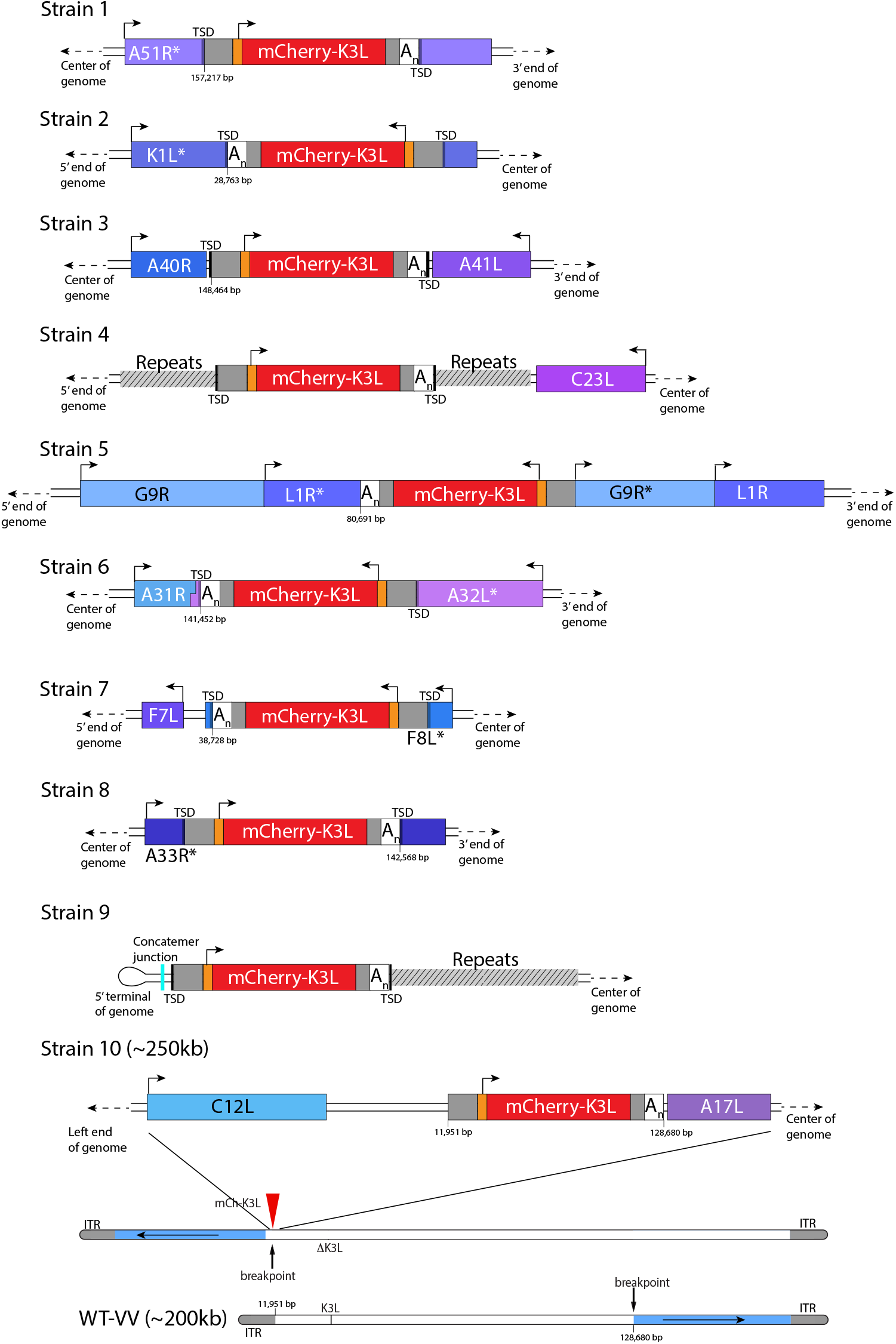
A detailed view of the region K3L inserted in each experimentally captured virus isolate. Arrows above cartoons indicate reading frame orientation. Flanking and/or interrupted (*) viral genes (blue/purple boxes), 3’/5’ untranslated regions (UTRs; gray), and poly(A) tails (An; white) are shown. Genomic rearrangements included in isolate 10 are shown below details. Some isolate 4 viruses include a second insertion that precisely matches that of isolate 2. We believe that this is due to recombination between the two strains during purification.

**Fig. S4.**
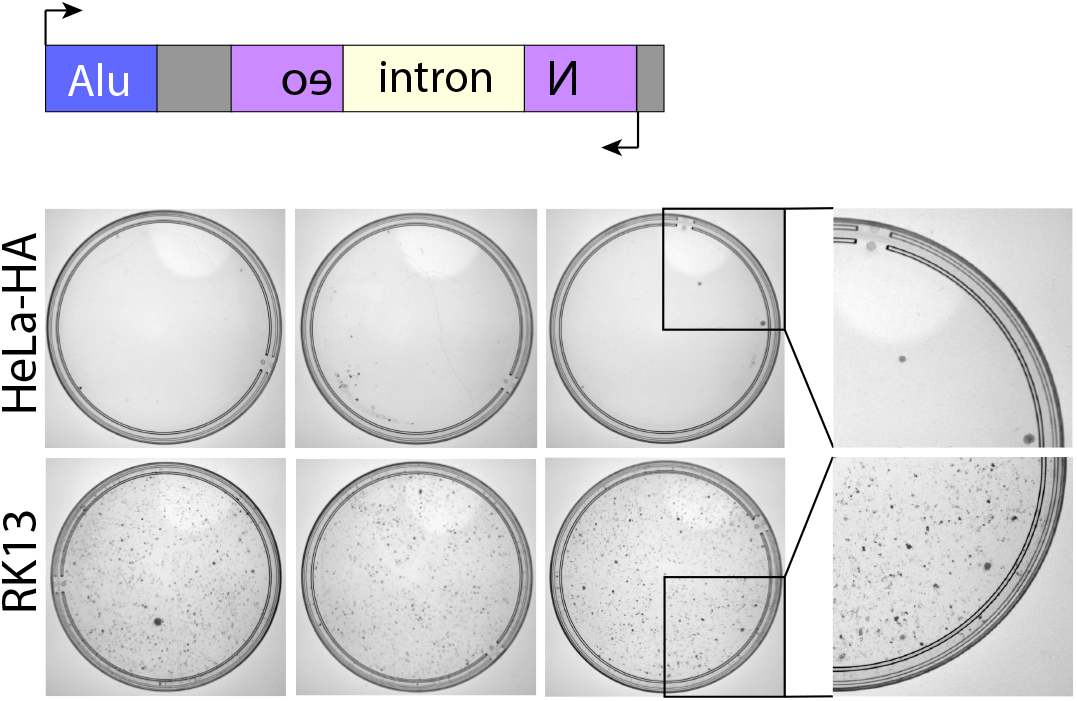
An Alu-Neo reporter (cartoon above) was used to interrogate the endogenous LINE-1 activity of the RK13-K3L cells compared to HeLa-HA cells. Below, representative plates from several experiments showing Neo-resistant colonies, with zoomed-in views to the right.

**Fig. S5.**
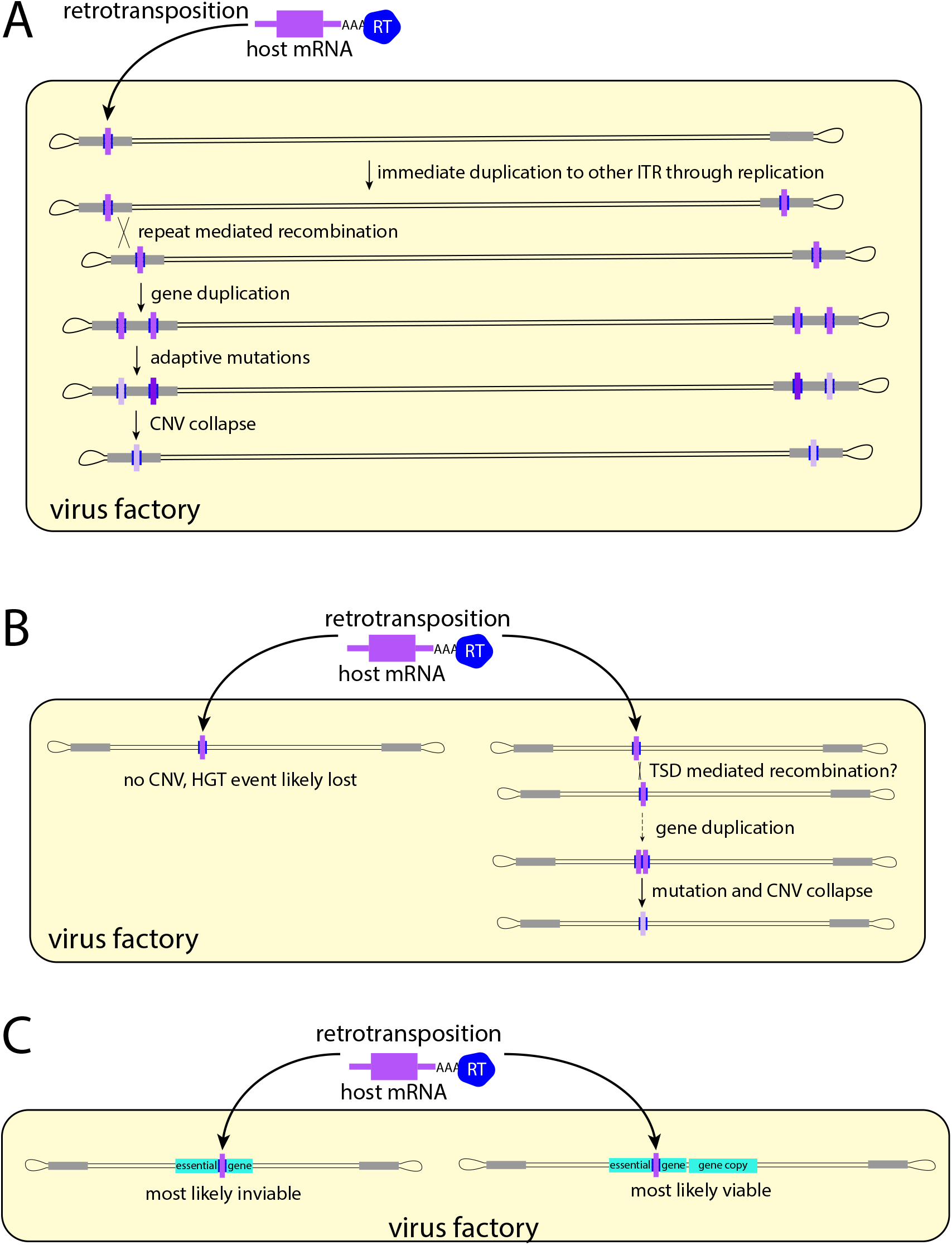
Models of evolution of newly acquired virus genes. (A) Genes that land in the repetitive region of the ITR are likely to be duplicated by both replication mechanisms (to the other ITR) and recombination mechanisms (tandem duplications). Duplications provide more chances for advantageous mutations to arise, increasing the likelihood of fixation. (B) Genes acquired outside the ITR may still be duplicated via TSD-mediated recombination. (C) Acquired genes that interrupt essential genes are unlikely to be maintained.

**Fig. S6.**
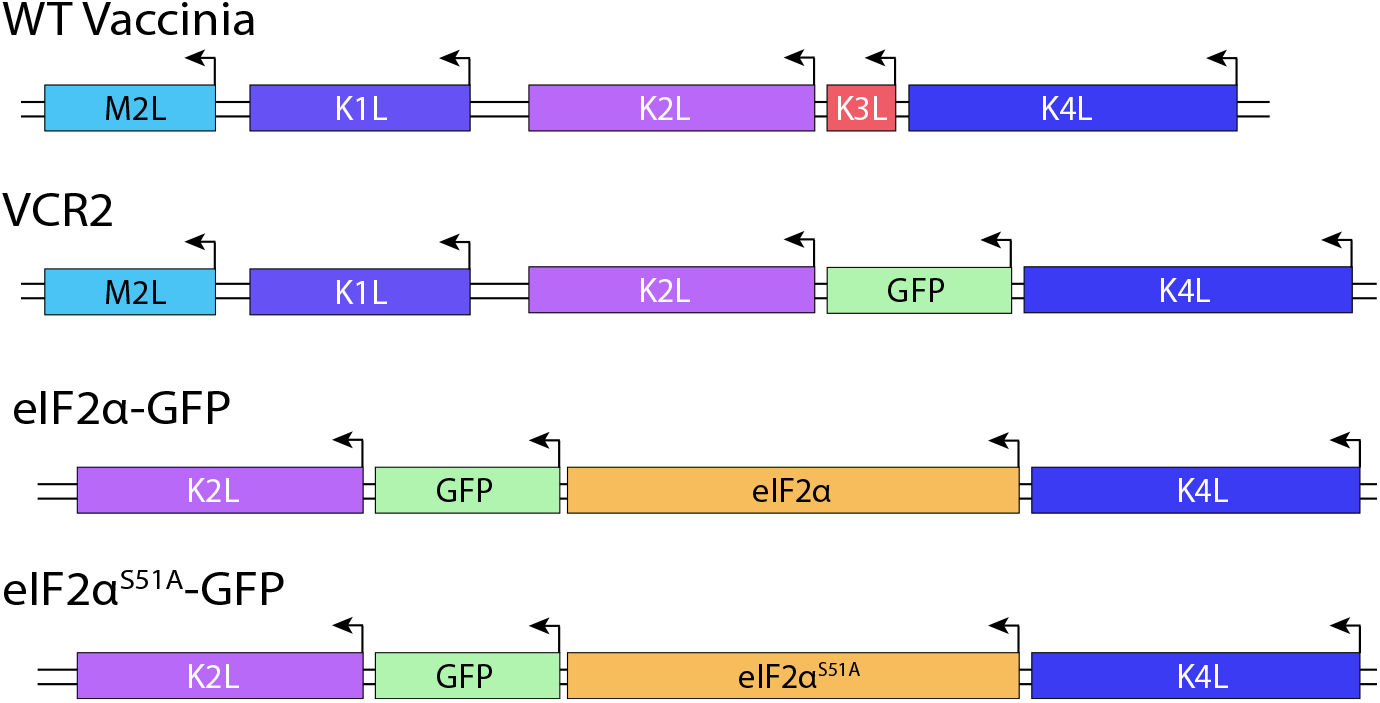
Structure of K3L region in eIF2α vaccinia strains engineered for experimental evolution compared to wild type (WT) vaccinia and VCR2.

**Table S1.**
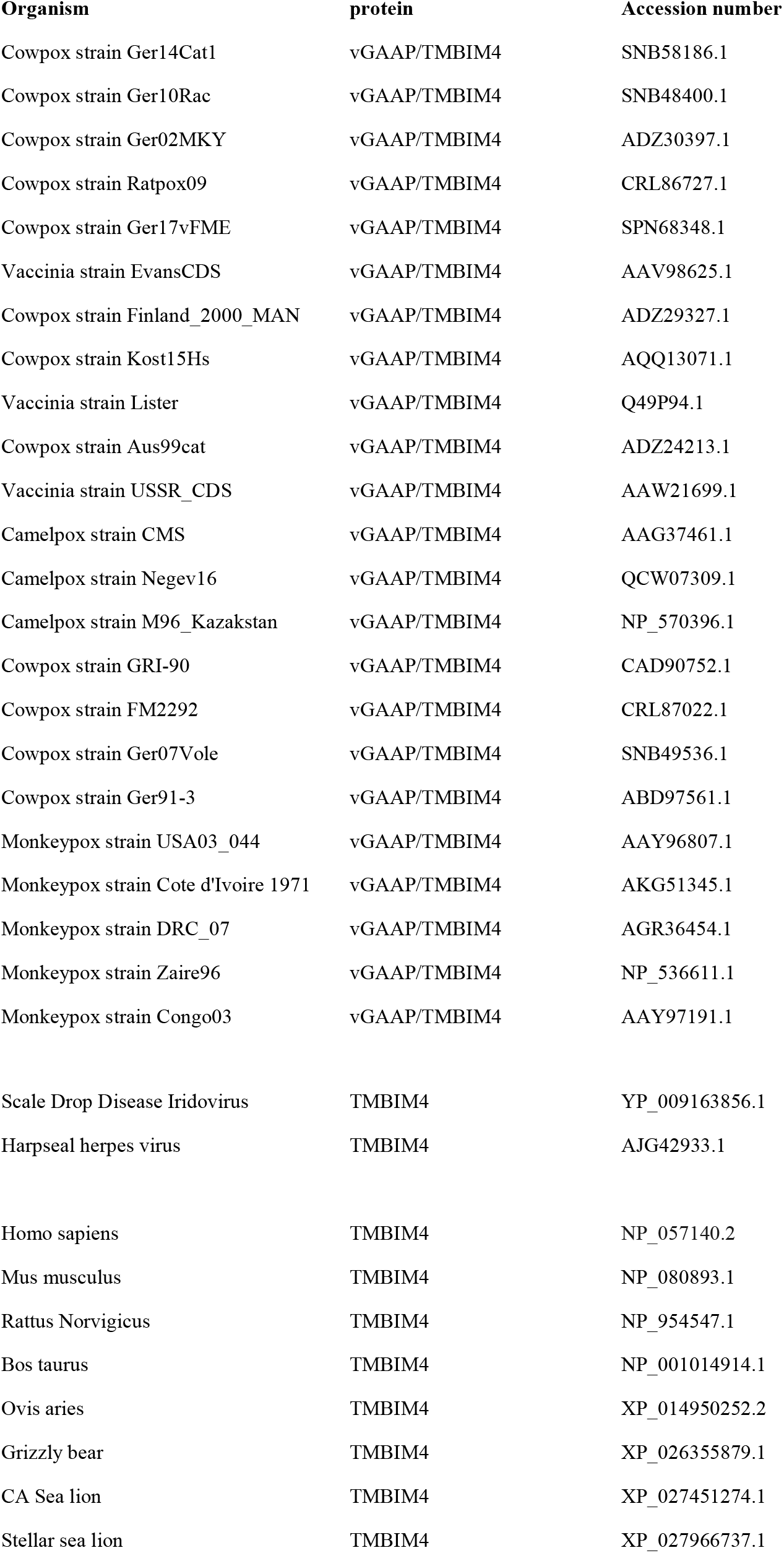
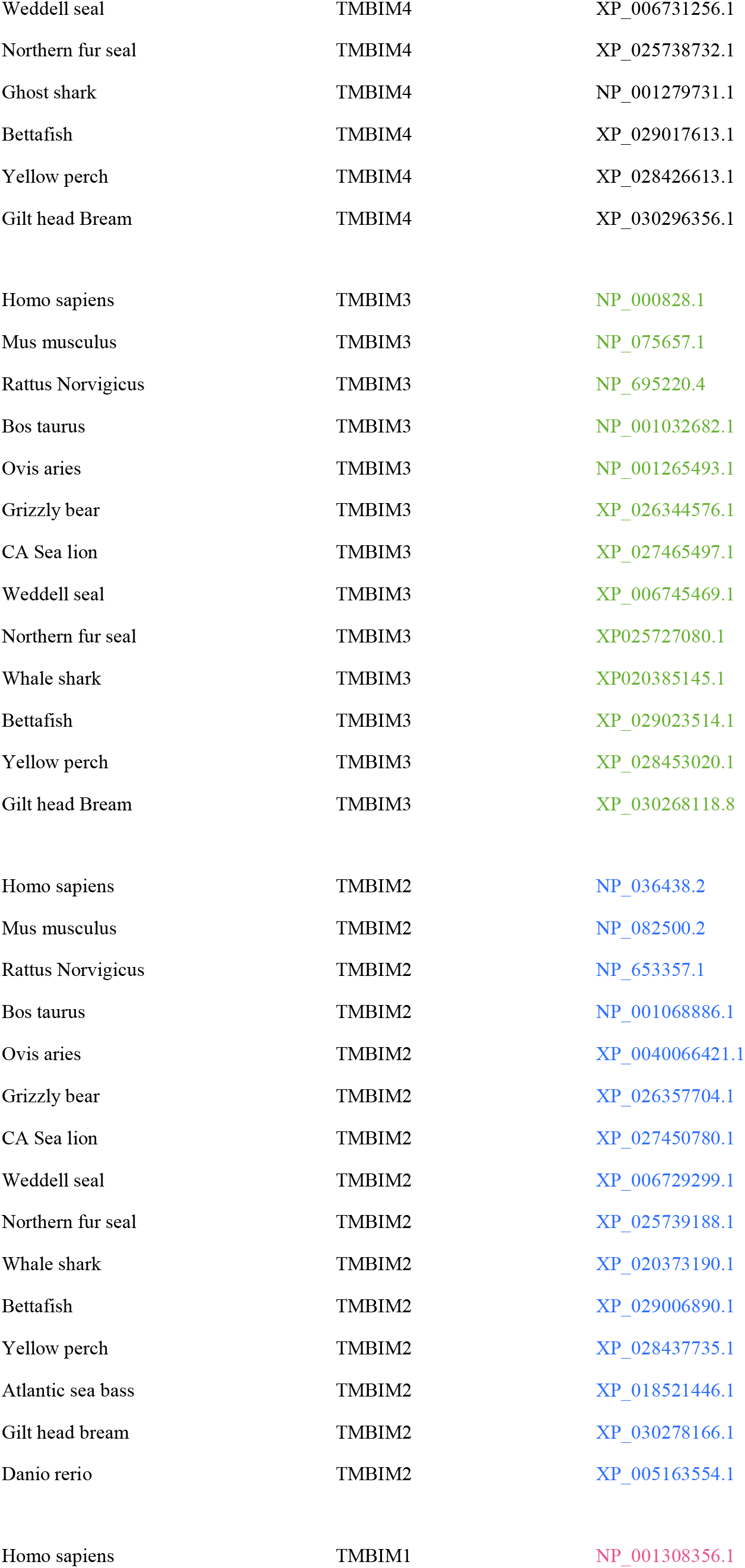
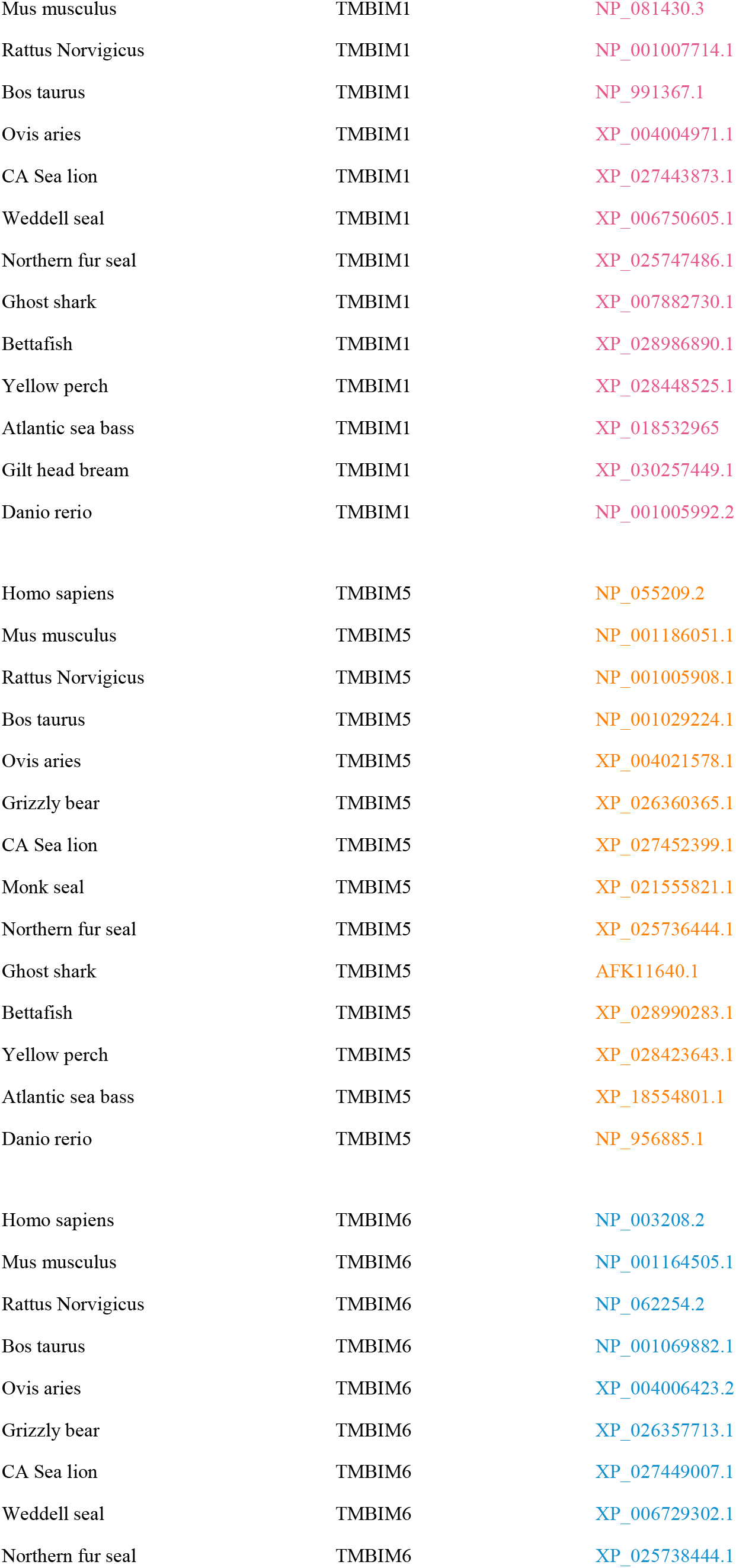
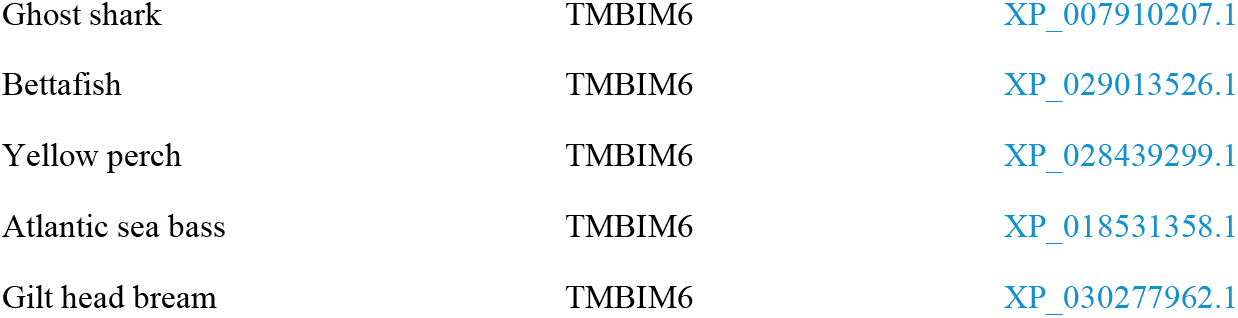
Accession numbers for sequences used in phylogenetic analysis (Fig. 1B)

**Table S2.**
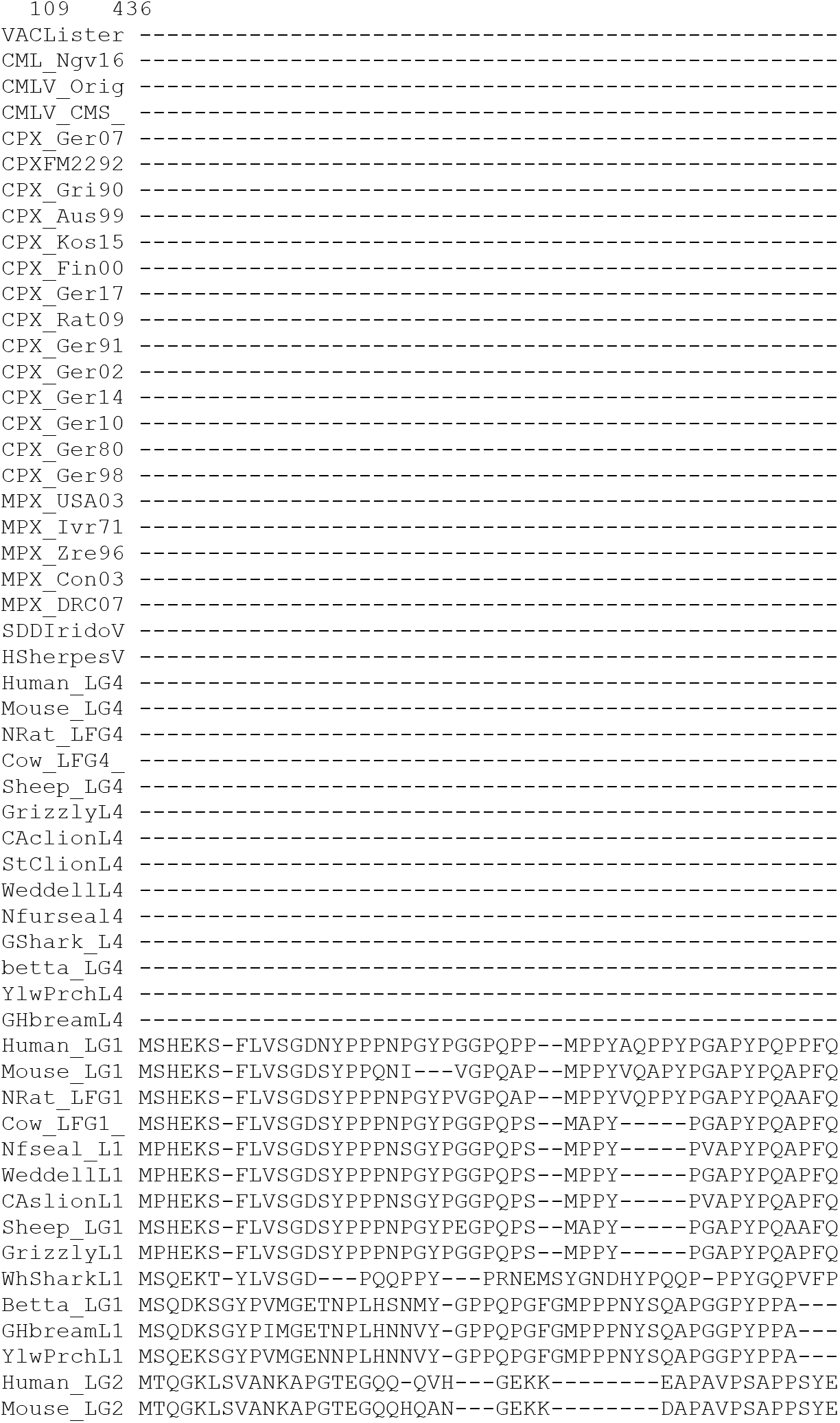
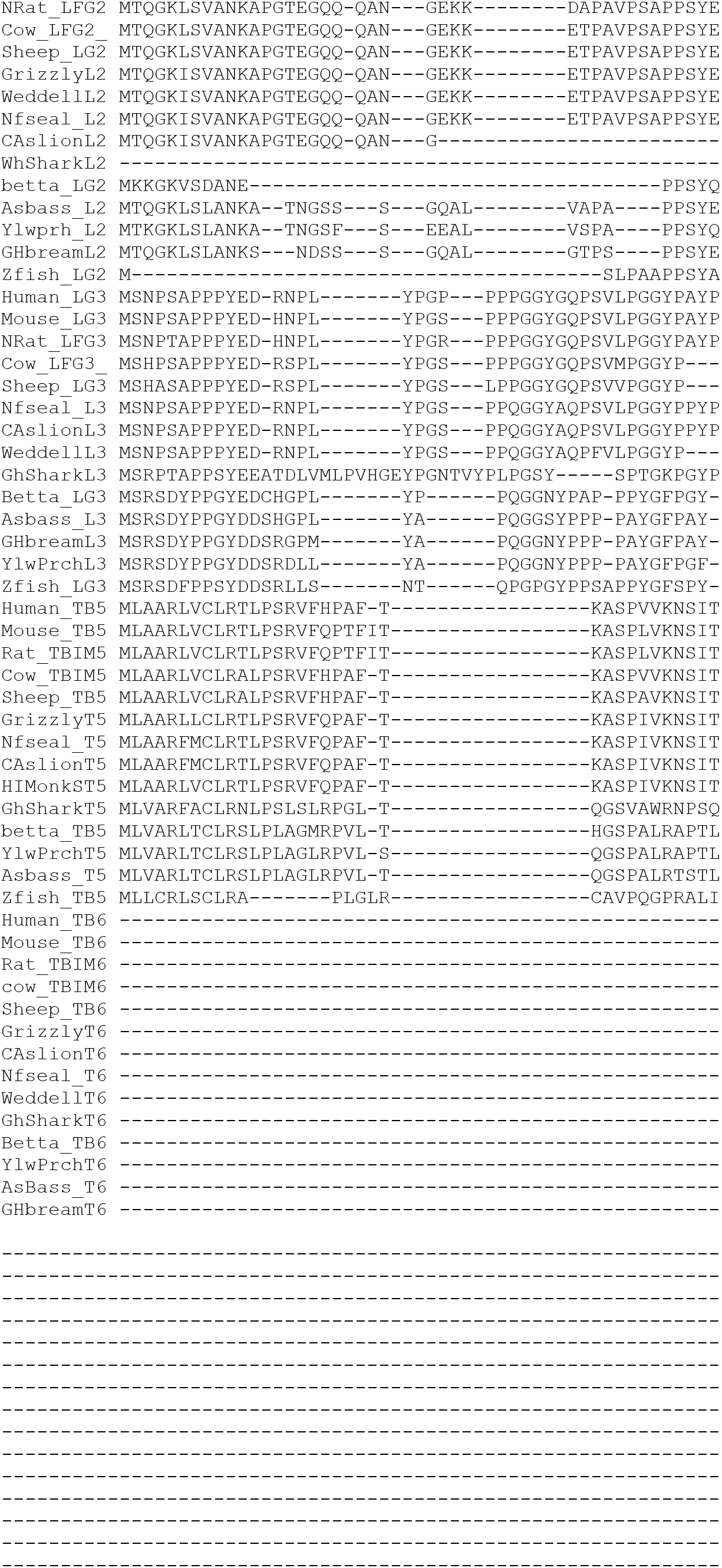
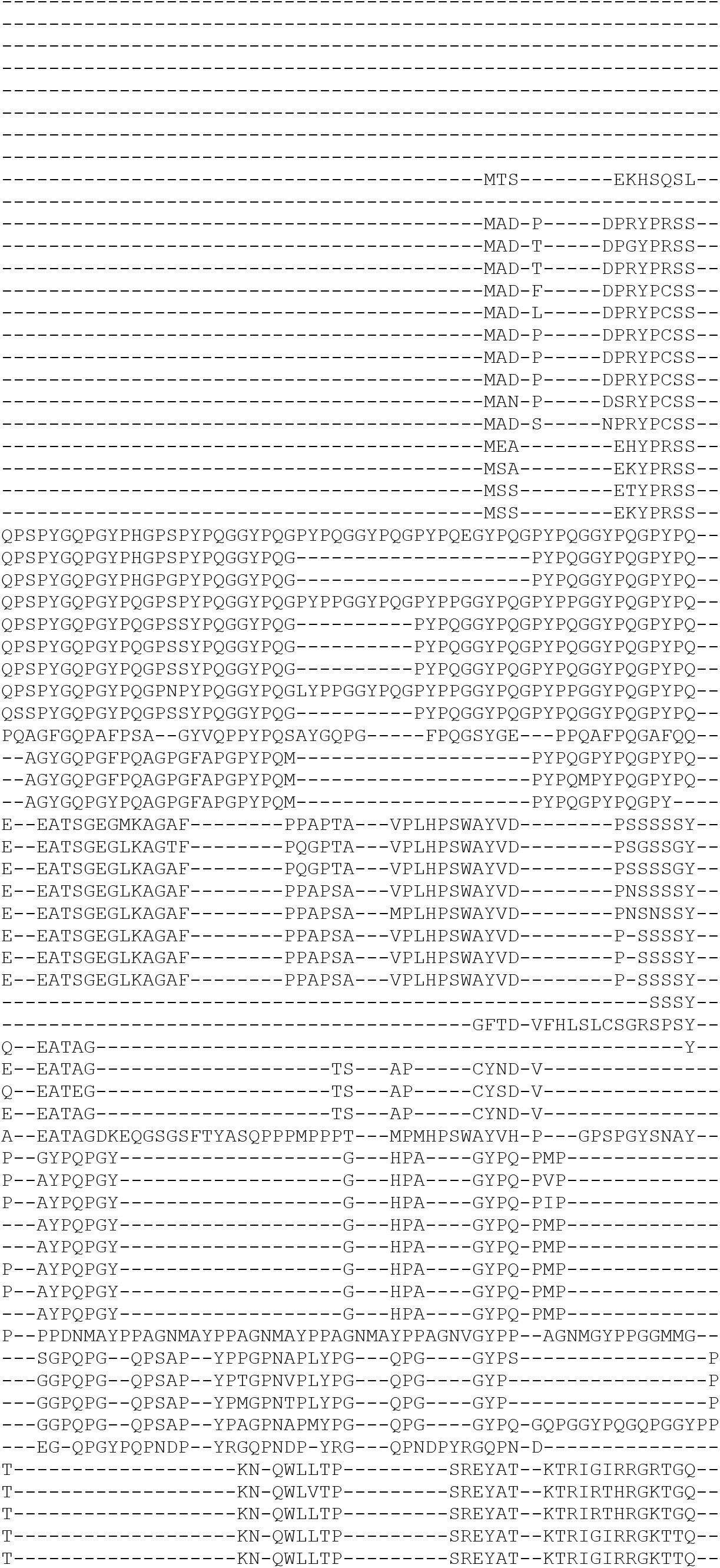
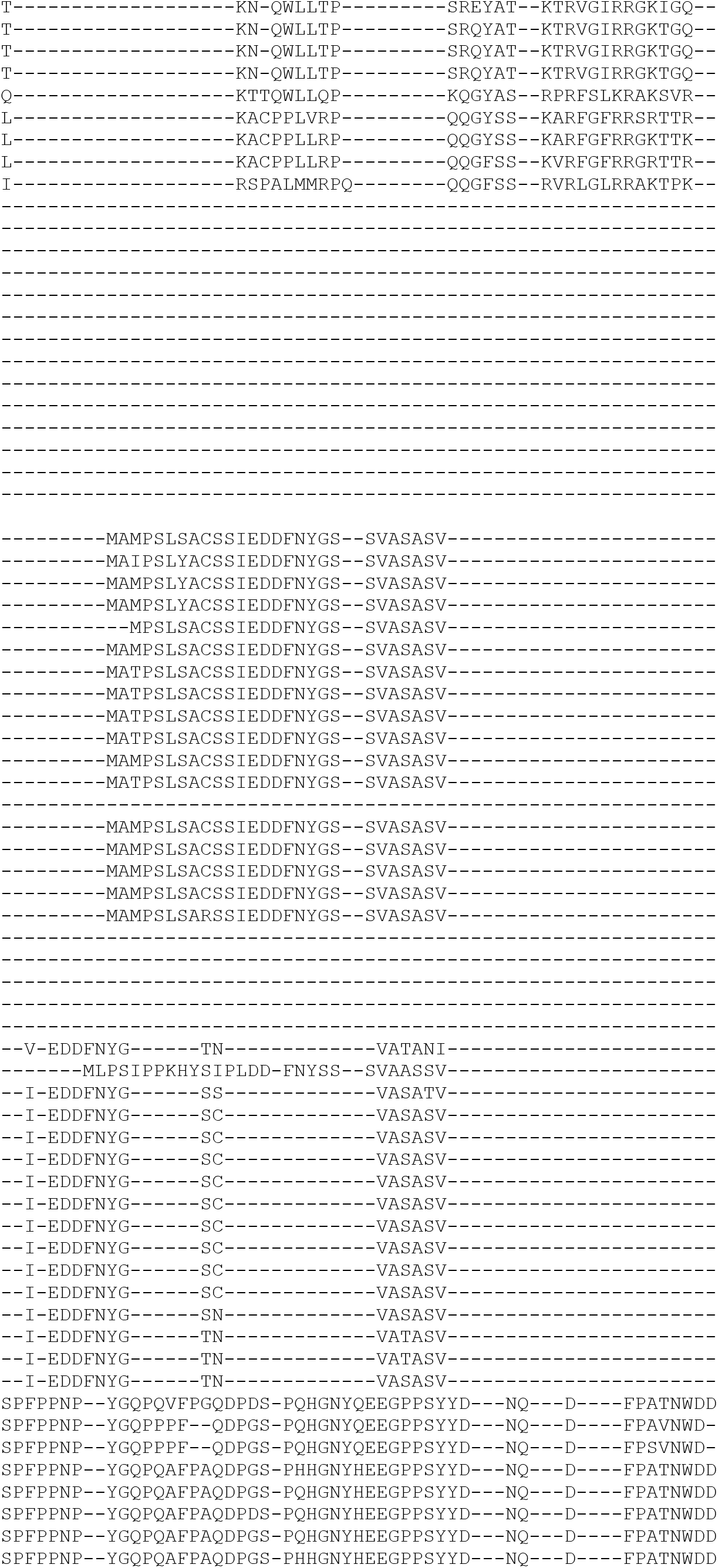
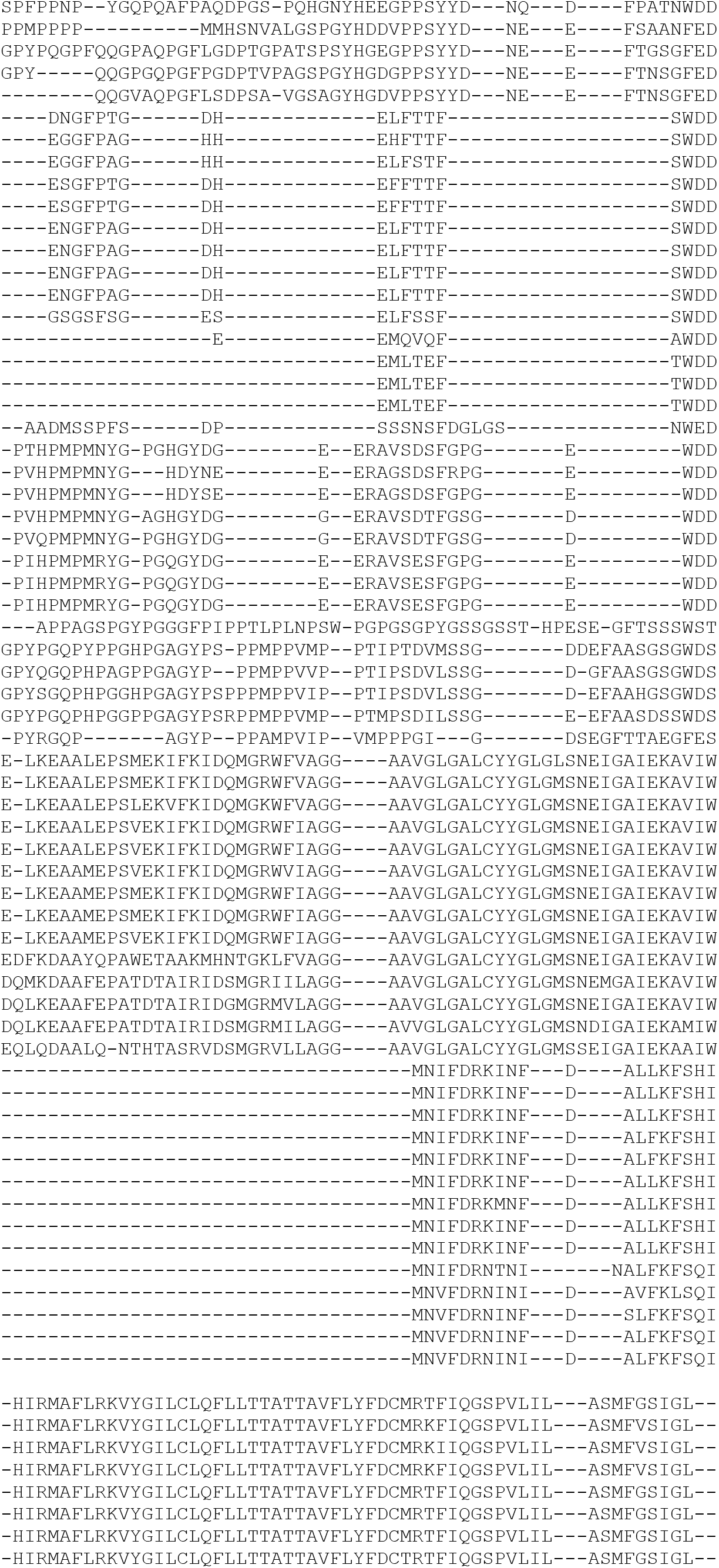
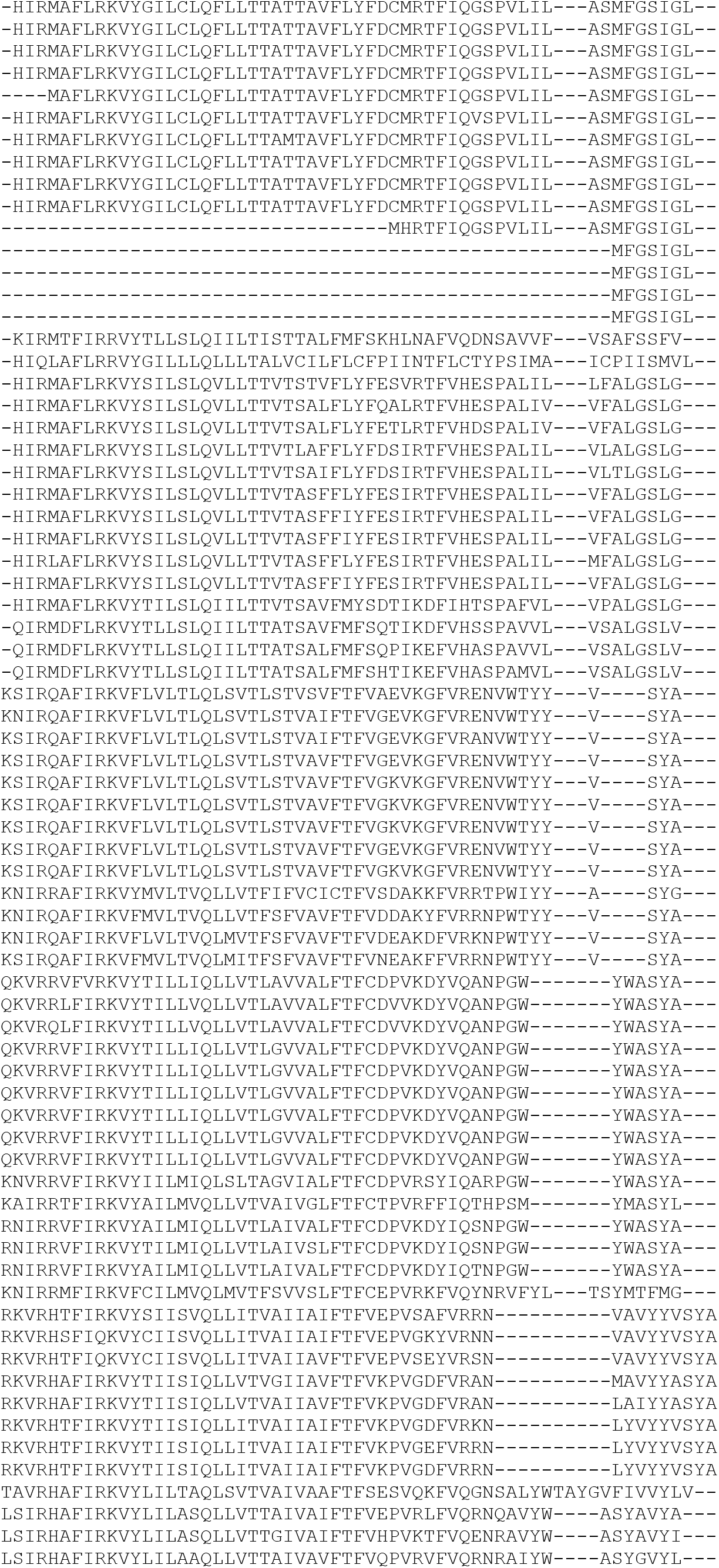
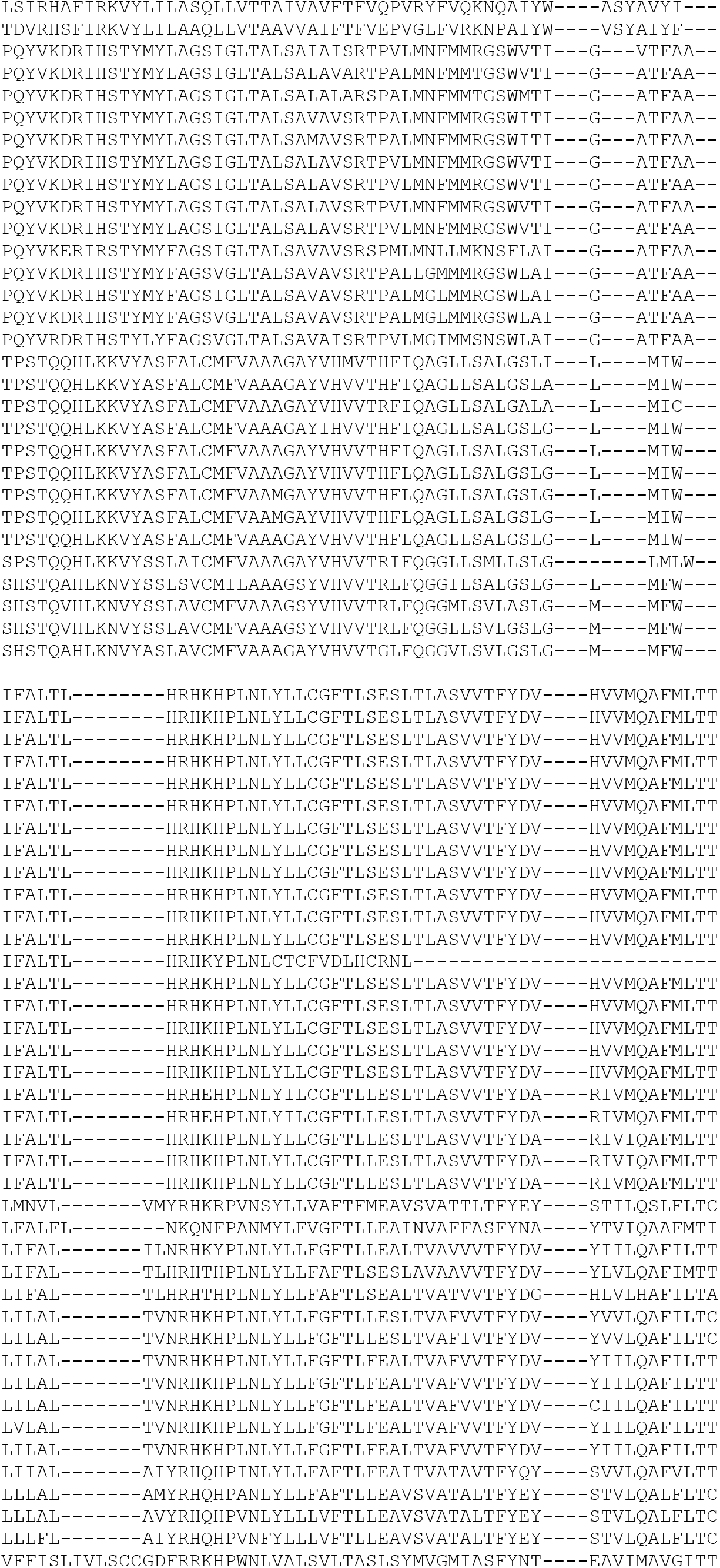
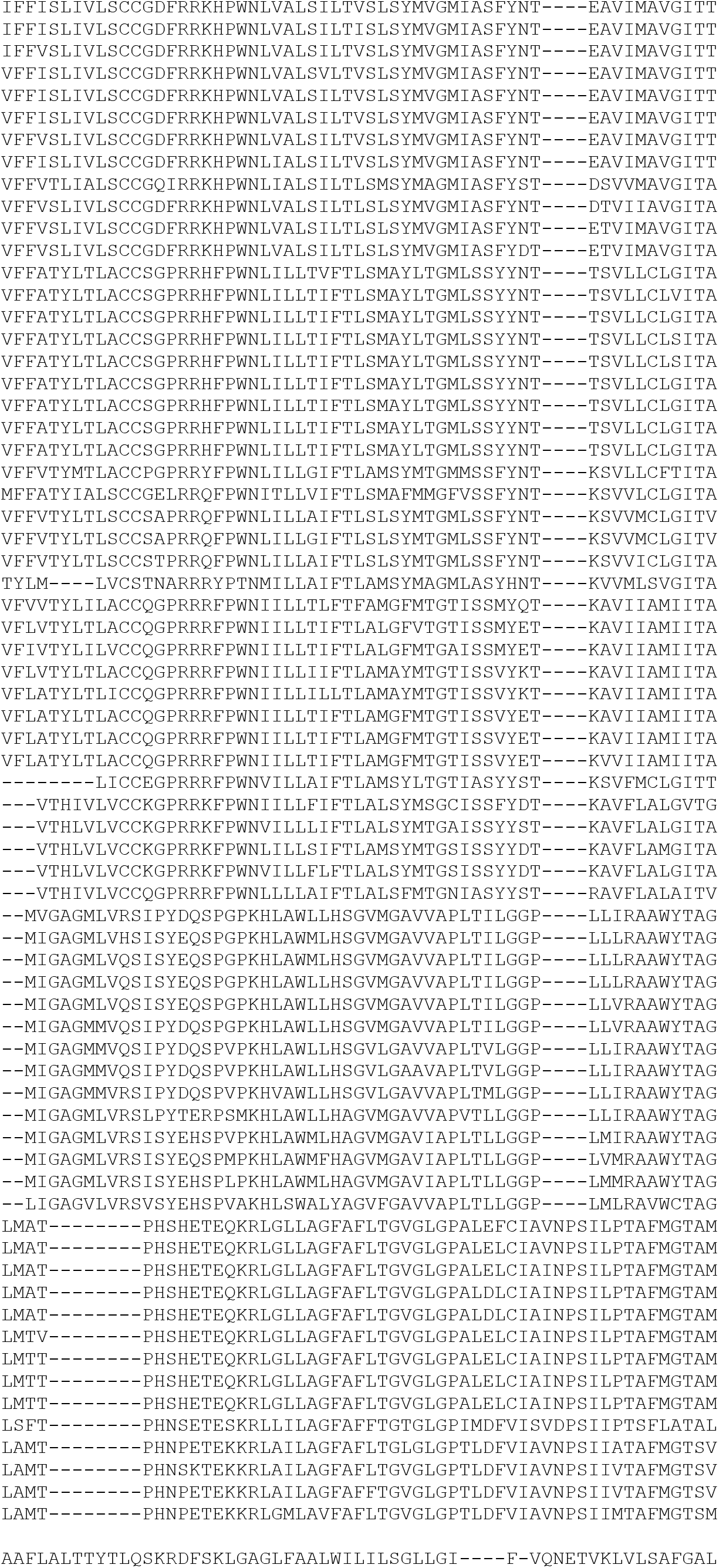
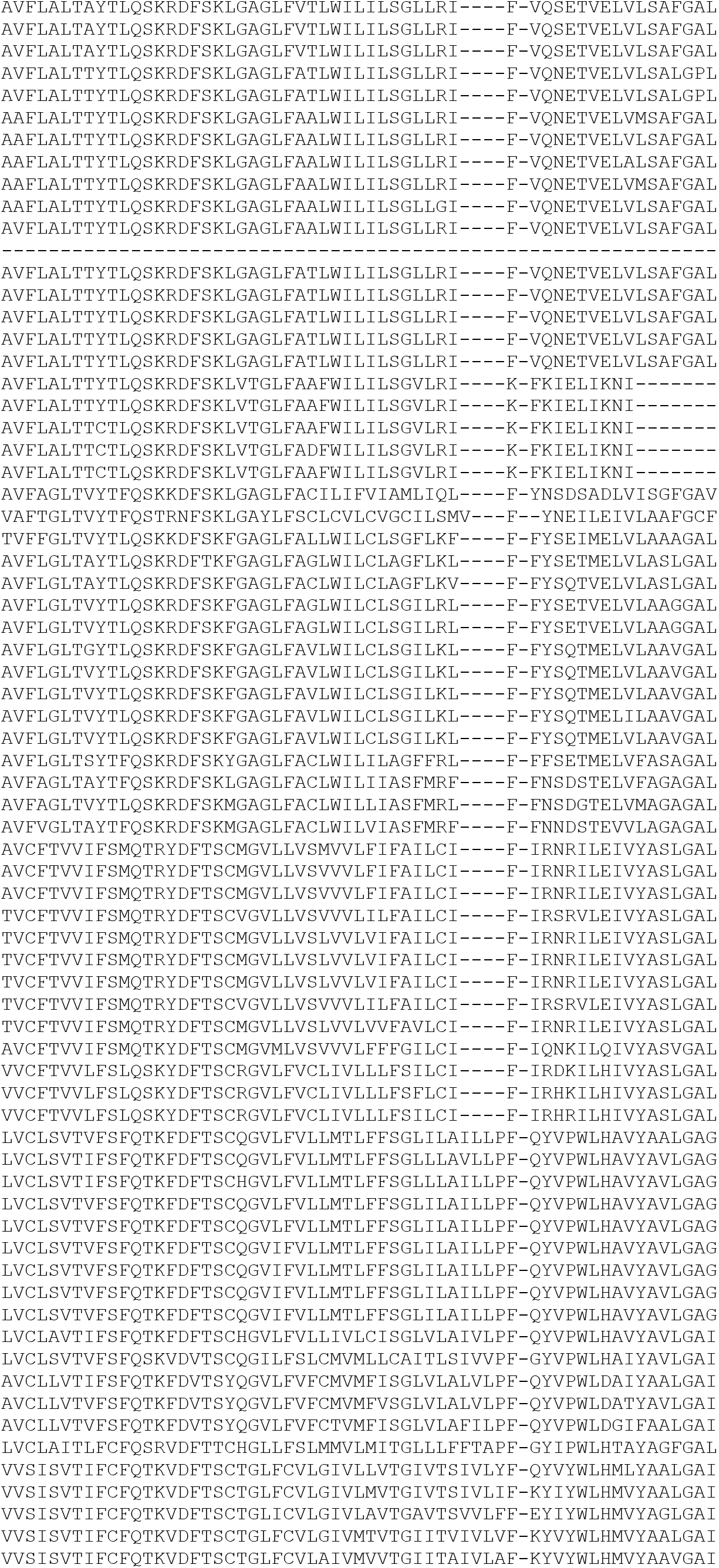
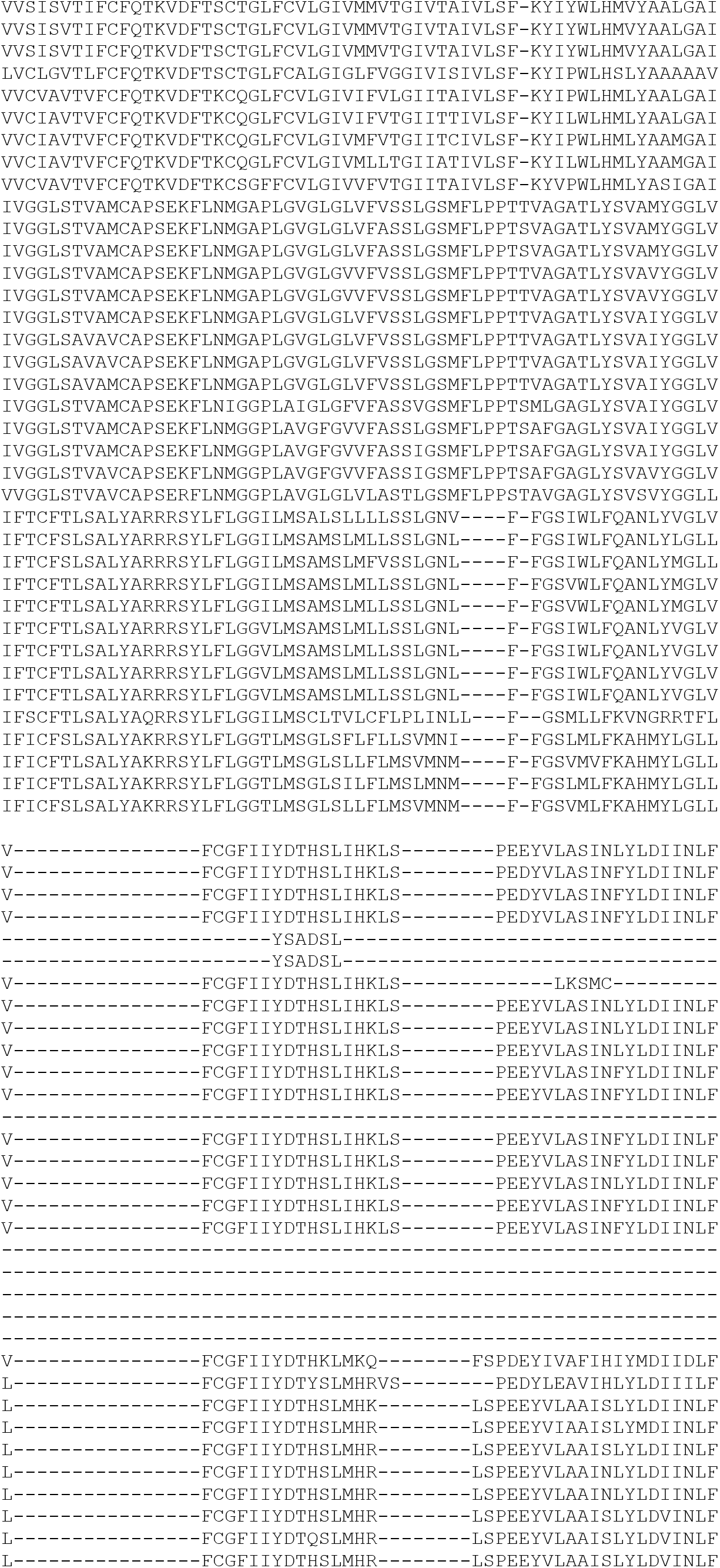
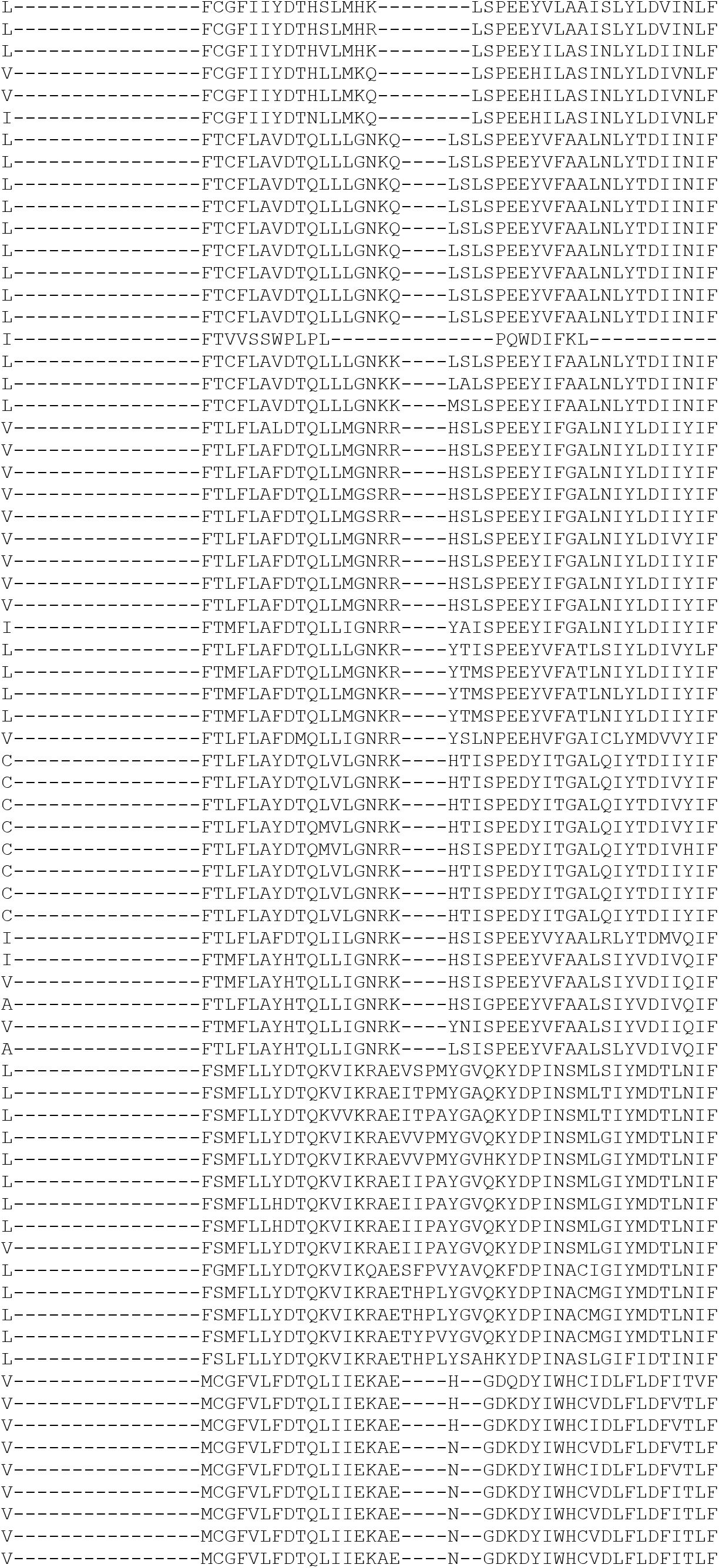
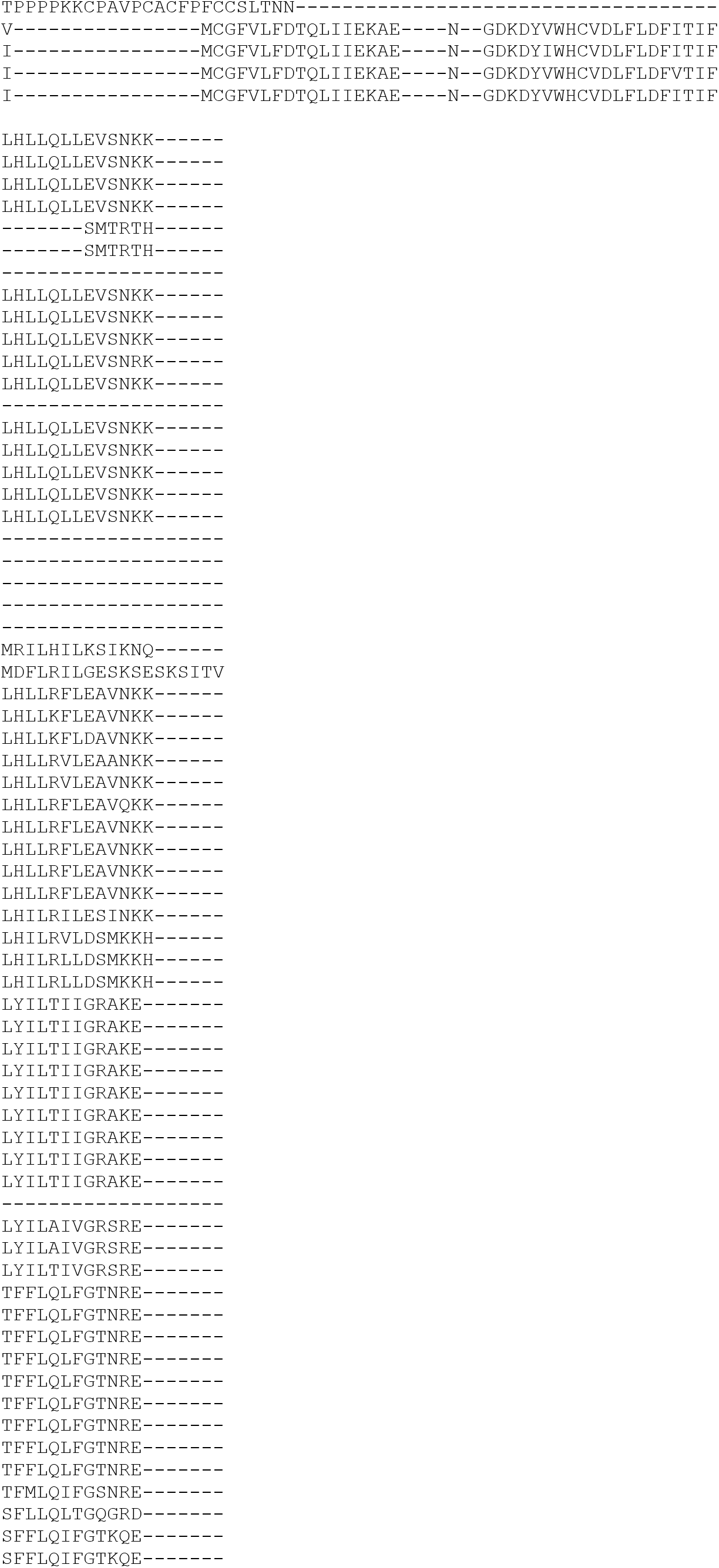
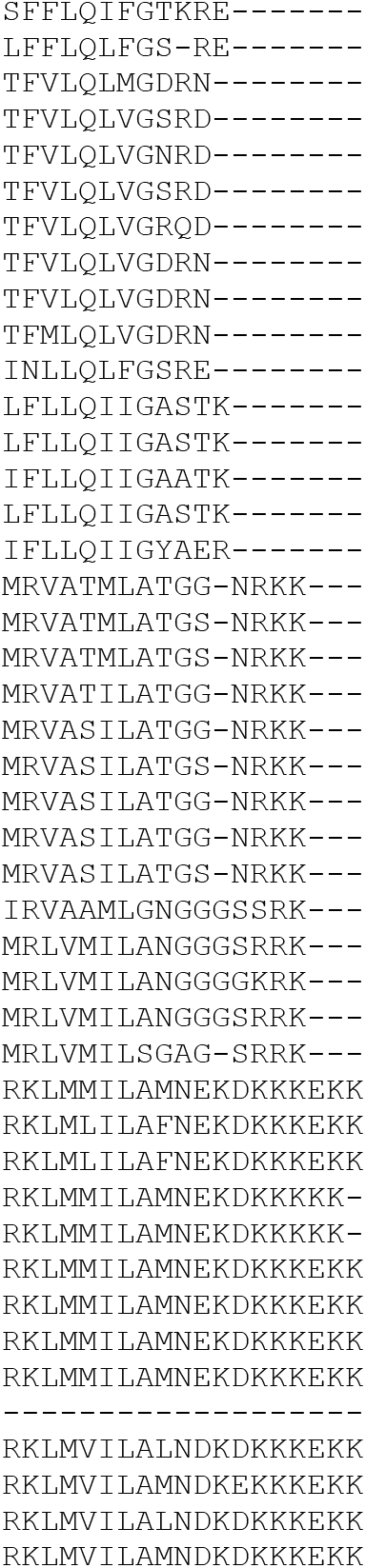
Phylip alignment for sequences used in phylogenetic analysis (Fig. 1B)

## Supplementary sequences

### piggybac_SLP-mCherry-K3L

actcttcctttttcaatattattgaagcatttatcagggttattgtctcatgagcggatacatatttgaatgtatttagaaaaataaacaaataggggttccgcgcacatttccccgaaaagtgccacctaaattgtaagcgttaatattttgttaaaattcgcgttaaatttttgttaaatcagctcattttttaaccaataggccgaaatcggcaaaatcccttataaatcaaaagaatagaccgagatagggttgagtgttgttccagtttggaacaagagtccactattaaagaacgtggactccaacgtcaaagggcgaaaaaccgtctatcagggcgatggcccactacgtgaaccatcaccctaatcaagttttttggggtcgaggtgccgtaaagcactaaatcggaaccctaaagggagcccccgatttagagcttgacggggaaagccggcgaacgtggcgagaaaggaagggaagaaagcgaaaggagcgggcgctagggcgctggcaagtgtagcggtcacgctgcgcgtaaccaccacacccgccgcgcttaatgcgccgctacagggcgcgtcccattcgccattcaggctgcgcaactgttgggaagggcgatcggtgcgggcctcttcgctattacgccagctggcgaaagggggatgtgctgcaaggcgattaagttgggtaacgccagggttttcccagtcacgacgttgtaaaacgacggccagtgagcgcgcctcgttcattcacgtttttgaacccgtggaggacgggcagactcgcggtgcaaatgtgttttacagcgtgatggagcagatgaagatgctcgacacgctgcagaacacgcagctagattaaccctagaaagataatcatattgtgacgtacgttaaagataatcatgcgtaaaattgacgcatgtgttttatcggtctgtatatcgaggtttatttattaatttgaatagatattaagttttattatatttacacttacatactaataataaattcaacaaacaatttatttatgtttatttatttattaaaaaaaaacaaaaactcaaaatttcttctataaagtaacaaaacttttatgagggacagcccccccccaaagcccccagggatgtaattacgtccctcccccgctagggggcagcagcgagccgcccggggctccgctccggtccggcgctccccccgcatccccgagccggcagcgtgcggggacagcccgggcacggggaaggtggcacgggatcgctttcctctgaacgcttctcgctgctctttgagcctgcagacacctggggggatacggggaaaaGGCCTCCACGGCCAGACTAGAACTTTTAAAAGAAAAGGGGGGATTGGGGGGTACAGTGCAGGGGAAAGAATAGTAGACATAATAGCAACAGACATACAAACTAAAGAATTACAAAAACAAATTACAAAAATTCAAAATTTTATCGATCACGAGACTAGCCTCGAGGGATCatccggtgcccgtcagtgggcagagcgcacatcgcccacagtccccgagaagttggggggaggggtcggcaattgaacgggtgcctagagaaggtggcgcggggtaaactgggaaagtgatgtcgtgtactggctccgcctttttcccgagggtgggggagaaccgtatataagtgcagtagtcgccgtgaacgttctttttcgcaacgggtttgccgccagaacacagctgaagcttcgaggggctcgcatctctccttcacgcgcccgccgccctacctgaggccgccatccacgccggttgagtcgcgttctgccgcctcccgcctgtggtgcctcctgaactgcgtccgccgtctaggtaagtttaaagctcaggtcgagaccgggcctttgtccggcgctcccttggagcctacctagactcagccggctctccacgctttgcctgaccctgcttgctcaactctacgtctttgtttcgttttctgttctgcgccgttacagatccaagctgtgaccggcgccGAATTCCTTCAGGGTGAGTTTGGGGACCCTTGATTGTTCTTTCTTTTTCGCTATTGTAAAATTCATGTTATATGGAGGGGGCAAAGTTTTCAGGGTGTTGTTTAGAATGGGAAGATGTCCCTTGTATCACCATGGACCCTCATGATAATTTTGTTTCTTTCACTTTCTACTCTGTTGACAACCATTGTCTCCTCTTATTTTCTTTTCATTTTCTGTAACTTTTTCGTTAAACTTTAGCTTGCATTTGTAACGAATTTTTAAATTCACTTTTGTTTATTTGTCAGATTGTAAGTACTTTCTCTAATCACTTTTTTTTCAAGGCAATCAGGGTATATTATATTGTACTTCAGCACAGTTTTAGAGAACAATTGTTATAATTAAATGATAAGGTAGAATATTTCTGCATATAAATTCTGGCTGGCGTGGAAATATTCTTATTGGTAGAAACAACTACATCCTGGTCATCATCCTGCCTTTCTCTTTATGGTTACAATGATATACACTGTTTGAGATGAGGATAAAATACTCTGAGTCCAAACCGGGCCCCTCTGCTAACCATGTTCATGCCTTCTTCTTTTTCCTACAGCTCCTGGGCAACGGTACCGGATCCCTGCAGAAGCTTCTAAAAATTGAAATTTTATTTTTTTTTTTTGGAATATAAATGGTGAGCAAGGGCGAGGAGGATAACATGGCCATCATCAAGGAGTTCATGCGCTTCAAGGTGCACATGGAGGGCTCCGTGAACGGCCACGAGTTCGAGATCGAGGGCGAGGGCGAGGGCCGCCCCTACGAGGGCACCCAGACCGCCAAGCTGAAGGTGACCAAGGGTGGCCCCCTGCCCTTCGCCTGGGACATCCTGTCCCCTCAGTTCATGTACGGCTCCAAGGCCTACGTGAAGCACCCCGCCGACATCCCCGACTACTTGAAGCTGTCCTTCCCCGAGGGCTTCAAGTGGGAGCGCGTGATGAACTTCGAGGACGGCGGCGTGGTGACCGTGACCCAGGACTCCTCCCTGCAGGACGGCGAGTTCATCTACAAGGTGAAGCTGCGCGGCACCAACTTCCCCTCCGACGGCCCCGTAATGCAGAAGAAGACCATGGGCTGGGAGGCCTCCTCCGAGCGGATGTACCCCGAGGACGGCGCCCTGAAGGGCGAGATCAAGCAGAGGCTGAAGCTGAAGGACGGCGGCCACTACGACGCTGAGGTCAAGACCACCTACAAGGCCAAGAAGCCCGTGCAGCTGCCCGGCGCCTACAACGTCAACATCAAGTTGGACATCACCTCCCACAACGAGGACTACACCATCGTGGAACAGTACGAACGCGCCGAGGGCCGCCACTCCACCGGCGGCATGGACGAGCTGTACAAGTCTAGATCTGCCACCATGCTTGCATTTTGTTATTCGTTGCCCAATCCTCACTTTGAAGCTATCTTGGCAGAGAGTGTTAAGATGCATATGGATAGATATGTTGAATATAGGGATAAACTGGTAGGGAAAACTGTAAAAGTTAAAGTGATTAGAGTTGATTATACAAAAGGATATATAGATGTCAATTACAAAAGGATGTGTAGACATCAATAATAgCGGCCGCCTAaacttgtttattgcagcttataatggttacaaataaagcaatagcatcacaaatttcacaaataaagcatttttttcactgcattctagttgtggtttgtccaaactcatcaatgtatcttatcatgtctGGATCCggcctccgcgccgggttttggcgcctcccgcgggcgcccccctcctcacggcgagcgctgccacgtcagacgaagggcgcagcgagcgtcctgatccttccgcccggacgctcaggacagcggcccgctgctcataagactcggccttagaaccccagtatcagcagaaggacattttaggacgggacttgggtgactctagggcactggttttctttccagagagcggaacaggcgaggaaaagtagtcccttctcggcgattctgcggagggatctccgtggggcggtgaacgccgatgattatataaggacgcgccgggtgtggcacagctagttccgtcgcagccgggatttgggtcgcggttcttgtttgtggatcgctgtgatcgtcacttggtgagtagcgggctgctgggctggccggggctttcgtggccgccgggccgctcggtgggacggaagcgtgtggagagaccgccaagggctgtagtctgggtccgcgagcaaggttgccctgaactgggggttggggggagcgcagcaaaatggcggctgttcccgagtcttgaatggaagacgcttgtgaggcgggctgtgaggtcgttgaaacaaggtggggggcatggtgggcggcaagaacccaaggtcttgaggccttcgctaatgcgggaaagctcttattcgggtgagatgggctggggcaccatctggggaccctgacgtgaagtttgtcactgactggagaactcggtttgtcgtctgttgcgggggcggcagttatggcggtgccgttgggcagtgcacccgtacctttgggagcgcgcgccctcgtcgtgtcgtgacgtcacccgttctgttggcttataatgcagggtggggccacctgccggtaggtgtgcggtaggcttttctccgtcgcaggacgcagggttcgggcctagggtaggctctcctgaatcgacaggcgccggacctctggtgaggggagggataagtgaggcgtcagtttctttggtcggttttatgtacctatcttcttaagtagctgaagctccggttttgaactatgcgctcggggttggcgagtgtgttttgtgaagttttttaggcaccttttgaaatgtaatcatttgggtcaatatgtaattttcagtgttagactagtaaattgtccgctaaattctggccgtttttggcttttttgttagacgctagcggatAGCGCTTCTCATTGGGCATTCCAGCCTACCCAGCTCGGAGTTAGTTACTCCGTAAGTGTGGCCGGAACAGAGTTCGTCCATCTAAAAAGGGAGGGGACCGGCGAACACATCTAGAGCCACCATGGCCACCGAGTACAAGCCCACGGTGCGCCTCGCCACCCGCGACGACGTCCCCCGGGCCGTACGCACCCTCGCCGCCGCGTTCGCCGACTACCCCGCCACGCGCCACACCGTCGAYCCGGACCGCCACATCGAGCGGGTCACCGAGCTGCAAGAACTCTTCCTCACGCGCGTCGGGCTCGACATCGGCAAGGTGTGGGTCGCGGACGACGGCGCCGCGGTGGCGGTCTGGACCACGCCGGAGAGCGTCGAAGCGGGGGCGGTGTTCGCCGAGATCGGCYCGCGCTGGCCGAGTTGAGCGGTTCCCGGCTGGCCGCGCAGCAACAGATGGAAGGCCTCCTGGCGCCGCACCGGCCCAAGGAGCCCGCGTGGTTCCTGGCCACCGTCGGCGTCTCGCCCGACCACCAGGGCAAGGGTCTGGGCAGCGCCGTCGTGCTCCCCGGAGTGGAGGCGGCCGAGCGCGCYGGGGTGCCCGCCTTCCTGGAGACCTCCGCGCCCCGCAACCTCCCCTTCTACGAGCGGCTCGGCTTCACCGTCACCGCCGACGTCGAGGTGCCCGAAGGACCGCGCACCTGGTGCATGACCCGCAAGCCCGGTGCCGTCGACAATCAACCTCTGGATTACAAAATTTGTGAAAGATTGACTGGTATTCTTAACTATGTTGCTCCTTTTACGCTATGTGGATACGCTGCTTTAATGCCTTTGTATCATGCGTTAACTAACTAaacttgtttattgcagcttataatggttacaaataaagcaatagcatcacaaatttcacaaataaagcatttttttcactgcattctagttgtggtttgtccaaactcatcaatgtatcttatcatgtctggaattgactcaaatgatgtcaattagtctatcagaagctatctggtctcccttccgggggacaagacatccctgtttaatatttaaacagcagtgttcccaaactgggttcttatatcccttgctctggtcaaccaggttgcagggtttcctgtcctcacaggaacgaagtccctaaagaaacagtggcagccaggtttagccccggaattgactggattccttttttagggcccattggtatggctttttccccgtatccccccaggtgtctgcaggctcaaagagcagcgagaagcgttcagaggaaagcgatcccgtgccaccttccccgtgcccgggctgtccccgcacgctgccggctcggggatgcggggggagcgccggaccggagcggagccccgggcggctcgctgctgccccctagcgggggagggacgtaattacatccctgggggctttgggggggggctgtccctgatatctataacaagaaaatatatatataataagttatcacgtaagtagaacatgaaataacaatataattatcgtatgagttaaatcttaaaagtcacgtaaaagataatcatgcgtcattttgactcacgcggtcgttatagttcaaaatcagtgacacttaccgcattgacaagcacgcctcacgggagctccaagcggcgactgagatgtcctaaatgcacagcgacggattcgcgctatttagaaagagagagcaatatttcaagaatgcatgcgtcaattttacgcagactatctttctagggttaatctagctgcatcaggatcatatcgtcgggtcttttttccggctcagtcatcgcccaagctggcgctatctgggcatcggggaggaagaagcccgtgccttttcccgcgaggttgaagcggcatggaaagagtttgccgaggatgactgctgctgcattgacgttgagcgaaaacgcacgtttaccatgatgattcgggaaggtgtggccatgcacgcctttaacggtgaactgttcgttcaggccacctgggataccagttcgtcgcggcttttccggacacagttccggatggtcagcccgaagcgcatcagcaacccgaacaataccggcgacagccggaactgccgtgccggtgtgcagattaatgacagcggtgcggcgctgggatattacgtcagcgaggacgggtatcctggctggatgccgcagaaatggacatggataccccgtgagttacccggcgggcgcgcttggcgtaatcatggtcatagctgtttcctgtgtgaaattgttatccgctcacaattccacacaacatacgagccggaagcataaagtgtaaagcctggggtgcctaatgagtgagctaactcacattaattgcgttgcgctcactgcccgctttccagtcgggaaacctgtcgtgccagctgcattaatgaatcggccaacgcgcggggagaggcggtttgcgtattgggcgctcttccgcttcctcgctcactgactcgctgcgctcggtcgttcggctgcggcgagcggtatcagctcactcaaaggcggtaatacggttatccacagaatcaggggataacgcaggaaagaacatgtgagcaaaaggccagcaaaaggccaggaaccgtaaaaaggccgcgttgctggcgtttttccataggctccgcccccctgacgagcatcacaaaaatcgacgctcaagtcagaggtggcgaaacccgacaggactataaagataccaggcgtttccccctggaagctccctcgtgcgctctcctgttccgaccctgccgcttaccggatacctgtccgcctttctcccttcgggaagcgtggcgctttctcatagctcacgctgtaggtatctcagttcggtgtaggtcgttcgctccaagctgggctgtgtgcacgaaccccccgttcagcccgaccgctgcgccttatccggtaactatcgtcttgagtccaacccggtaagacacgacttatcgccactggcagcagccactggtaacaggattagcagagcgaggtatgtaggcggtgctacagagttcttgaagtggtggcctaactacggctacactagaaggacagtatttggtatctgcgctctgctgaagccagttaccttcggaaaaagagttggtagctcttgatccggcaaacaaaccaccgctggtagcggtggtttttttgtttgcaagcagcagattacgcgcagaaaaaaaggatctcaagaagatcctttgatcttttctacggggtctgacgctcagtggaacgaaaactcacgttaagggattttggtcatgagattatcaaaaaggatcttcacctagatccttttaaattaaaaatgaagttttaaatcaatctaaagtatatatgagtaaacttggtctgacagttaccaatgcttaatcagtgaggcacctatctcagcgatctgtctatttcgttcatccatagttgcctgactccccgtcgtgtagataactacgatacgggagggcttaccatctggccccagtgctgcaatgataccgcgagacccacgctcaccggctccagatttatcagcaataaaccagccagccggaagggccgagcgcagaagtggtcctgcaactttatccgcctccatccagtctattaattgttgccgggaagctagagtaagtagttcgccagttaatagtttgcgcaacgttgttgccattgctacaggcatcgtggtgtcacgctcgtcgtttggtatggcttcattcagctccggttcccaacgatcaaggcgagttacatgatcccccatgttgtgcaaaaaagcggttagctccttcggtcctccgatcgttgtcagaagtaagttggccgcagtgttatcactcatggttatggcagcactgcataattctcttactgtcatgccatccgtaagatgcttttctgtgactggtgagtactcaaccaagtcattctgagaatagtgtatgcggcgaccgagttgctcttgcccggcgtcaatacgggataataccgcgccacatagcagaactttaaaagtgctcatcattggaaaacgttcttcggggcgaaaactctcaaggatcttaccgctgttgagatccagttcgatgtaacccactcgtgcacccaactgatcttcagcatcttttactttcaccagcgtttctgggtgagcaaaaacaggaaggcaaaatgccgcaaaaaagggaataagggcgacacggaaatgttgaatactcat

### pBlue_165_mCherry-K3L

GAGCTCtggcaacctagataaaaaatcaactattgttttatccactttctcgtatgttcgaatgagattataatcctgtattatatgggatgtggaagaattggaacacacgcgtgctatataatgaagagataaatatacactccagtcaagtatttcctttttaaaaaaatccatatataatttatattctgtaacatgttatcccttttcaattaacaatgttagtttataaaaaattaaagaagcgaatcaatgattaatagatgttaagaactataattacgatgtattaataggtatatagttagttggttagttaaaaaagagataacagttactaattaattgttagttattgtctatatgatattacaacctattatttgttctctatagttacattaattaaaattttatatgtgacacccattcatctggagaatacttcttgataccattagtatccatataGGAATATAAATGGTGAGCAAGGGCGAGGAGGATAACATGGCCATCATCAAGGAGTTCATGCGCTTCAAGGTGCACATGGAGGGCTCCGTGAACGGCCACGAGTTCGAGATCGAGGGCGAGGGCGAGGGCCGCCCCTACGAGGGCACCCAGACCGCCAAGCTGAAGGTGACCAAGGGTGGCCCCCTGCCCTTCGCCTGGGACATCCTGTCCCCTCAGTTCATGTACGGCTCCAAGGCCTACGTGAAGCACCCCGCCGACATCCCCGACTACTTGAAGCTGTCCTTCCCCGAGGGCTTCAAGTGGGAGCGCGTGATGAACTTCGAGGACGGCGGCGTGGTGACCGTGACCCAGGACTCCTCCCTGCAGGACGGCGAGTTCATCTACAAGGTGAAGCTGCGCGGCACCAACTTCCCCTCCGACGGCCCCGTAATGCAGAAGAAGACCATGGGCTGGGAGGCCTCCTCCGAGCGGATGTACCCCGAGGACGGCGCCCTGAAGGGCGAGATCAAGCAGAGGCTGAAGCTGAAGGACGGCGGCCACTACGACGCTGAGGTCAAGACCACCTACAAGGCCAAGAAGCCCGTGCAGCTGCCCGGCGCCTACAACGTCAACATCAAGTTGGACATCACCTCCCACAACGAGGACTACACCATCGTGGAACAGTACGAACGCGCCGAGGGCCGCCACTCCACCGGCGGCATGGACGAGCTGTACAAGTCTAGATCTGCCACCATGCTTGCATTTTGTTATTCGTTGCCCAATGCGGGcGATGTAATAAAGGGCAGAGTATACGAGAAGGATTATGCTCTATAcATTTATCTTTTTGACTATCCTCACTcTGAAGCTATCTTGGCAGAGAGTGTTAAGATGCATATGGATAGATATGTTGAATATAGGGATAAACTGGTAGGGAAAACTGTAAAAGTTAAAGTGATTAGAGTTGATTATACAAAAGGATATATAGATGTCAATTACAAAAGGATGTGTAGACATCAATAAtaggcgtttaaacaattggaATCAGCaggctattcggctatgactgggcacaacagacaatcggctgctctgatgccgccgtgttccggctgtcagcgcaggggcgcccggttctttttgtcaagaccgacctgtccggtgccctgaatgaactgcaggacgaggcagcgcggctatcgtggctggccacgacgggcgttccttgcgcagctgtgctcgacgttgtcactgaagcgggaagggactggctgctattgggcgaagtgccggggcaggatctcctgtcatctcaccttgctcctgccgagaaagtatccatcatggctgatgcaatgcggcggctgcatacgcttgatccggctacctgcccattcgaccaccaagcgaaacatcgcatcgagcgagcacgtactcggatggaagccggtcttgtcgatcaggatgatctggacgaagagcatcaggggctcgcgccagccgaactgttcgccaggctcaaggcgcgcatgcccgacggcgaggatctcgtcgtgacccatggcgatgcctgcttgccgaatatcatggtggaaaatggccgcttttctggattcatcgactgtggccggctgggtgtggcggaccgctatcaggacatagcgttggctacccgtgatattgctgaagagcttggcggcgaatgggctgaccgcttcctcgtgctttacggtatcgccgctcccgattcgcagcgcatcgccttctatcgccttcttgacgagttcttctgaGCGGGACTCTGGGGTTCGCGAAATGATCGACCAAGCGACGCCCAACCTGCCATCACGAGATTTCGATTCCACCGCCGCCTTCTATGAAAGGTTGGGCTTCGGAATCGTTTTCCGGGACGCCGGCTGGATGATCCTCCAGCGCGGGGATCTCATGCTGGAGTTCTTCGCCCACCCCAACTTGTTTATTGCAGCTTATAATGGTTACAAATAAAGCAATAGatagcgcacacgtatgtcgaggtagcctcatccccaggtttatataccttgatgaatcgacacgcgtacttgatgtcctctttcttctcacagaaatgcacaacacaaagtcttttgatatgcttttctatatcattctctaccagtctgaggtagtcgtaacccaccgtgaatccaaatactttgtgtgtattattatcaactccaatgaacaaatatccaccctttgtgttggtaaaagatgagagtataaaggaagctgctgccgtatacgtgttcttagttgcttagctgaaacagatgtatgtttaacaattaatagaattaccaaatttgactaatttaccagcctgaagttctgatctattgaagaactcctctaccaatctctcaattgattcagtgtcttccactccatctggatattcaaattcctgcatttctggtctgggactccatccacctgattccttcagttcgctGCTAGCCATATGGGTACC

## Isolate sequences

### Isolate 1

caagtcccataccgcaaatggaatatccattagatttcctagatttactaggataagagaggataaaacgtggaaagaatctactcatctaaacgatttagtaaacttgactaaatcttaatagttacatacaaactgaaaattaaaataacaccatttagttggtggtcgccatggatggtgttattgtatactgtctaaacgcgttagtaaaacatggcgaggaaataaatcatataaaaaatgatttcatgattaaaccatgttgtgaaagagtttgtgaaaaagttaagaacgttcacattggcggacaatctaaaaacaatacagtgattgcagatttgccatatatggataatgctgtatcggatgtatgcaattcactgtataaaaagaatgtatcaagaatatccagatttgctaatttgataaagatagatgacgatgacaagactcctactggtgtatataattattttaaacctaaagatgttattcctgttatcatatctataggaaaggataaagatgtctgtgaactattaatctcatcagacatatcgtgtgcatgcgtggagttaaattcatataaagtagccattcttcccatggatgtttccttttttaccaaaggaaatgcatcattgattattctcctgtttgatttctctatcgatgcagcacctctcttaagaagtgtaaccgatagctttttcgcaacgggtttgccgccagaacacagctgaagcttcgaggggctcgcatctctccttcacgcgcccgccgccctacctgaggccgccatccacgccggttgagtcgcgttctgccgcctcccgcctgtggtgcctcctgaactgcgtccgccgtctagCTCCTGGGCAACGGTACCGGATCCCTGCAGAAGCTTCTAAAAATTGAAATTTTATTTTTTTTTTTTGGAATATAAATGGTGAGCAAGGGCGAGGAGGATAACATGGCCATCATCAAGGAGTTCATGCGCTTCAAGGTGCACATGGAGGGCTCCGTGAACGGCCACGAGTTCGAGATCGAGGGCGAGGGCGAGGGCCGCCCCTACGAGGGCACCCAGACCGCCAAGCTGAAGGTGACCAAGGGTGGCCCCCTGCCCTTCGCCTGGGACATCCTGTCCCCTCAGTTCATGTACGGCTCCAAGGCCTACGTGAAGCACCCCGCCGACATCCCCGACTACTTGAAGCTGTCCTTCCCCGAGGGCTTCAAGTGGGAGCGCGTGATGAACTTCGAGGACGGCGGCGTGGTGACCGTGACCCAGGACTCCTCCCTGCAGGACGGCGAGTTCATCTACAAGGTGAAGCTGCGCGGCACCAACTTCCCCTCCGACGGCCCCGTAATGCAGAAGAAGACCATGGGCTGGGAGGCCTCCTCCGAGCGGATGTACCCCGAGGACGGCGCCCTGAAGGGCGAGATCAAGCAGAGGCTGAAGCTGAAGGACGGCGGCCACTACGACGCTGAGGTCAAGACCACCTACAAGGCCAAGAAGCCCGTGCAGCTGCCCGGCGCCTACAACGTCAACATCAAGTTGGACATCACCTCCCACAACGAGGACTACACCATCGTGGAACAGTACGAACGCGCCGAGGGCCGCCACTCCACCGGCGGCATGGACGAGCTGTACAAGTCTAGATCTGCCACCATGCTTGCATTTTGTTATTCGTTGCCCAATGCGGGcGATGTAATAAAGGGCAGAGTATACGAGAAGGATTATGCTCTATAcATTTATCTTTTTGACTATCCTCACTcTGAAGCTATCTTGGCAGAGAGTGTTAAGATGCATATGGATAGATATGTTGAATATAGGGATAAACTGGTAGGGAAAACTGTAAAAGTTAAAGTGATTAGAGTTGATTATACAAAAGGATATATAGATGTCAATTACAAAAGGATGTGTAGACATCAATAATAgGCGGCCGCCTAaacttgtttattgcagcttataatggttacaaataaagcaatagcatcacaaaAAAAAAAAAAAAAAAAAAAAAAAAAAAAAAAAAAAAAAAAAAAAAAAgaagtgtaaccgataataatgttattatatctagacaccagcgtctacatgacgagcttccgagttccaattggttcaagttttacataagtataaagtccgactattgttctatattatatatggttgttgatggatctgtgatgatgcgatagctgataatagaactcacgcaattattagcaaaaatatattagacaatactacgattaacgatgagtgtagatgctgttattttgaaccacagattaggattcttgatagagatgagatgctcaatggatcatcgtgtgatatgaacagacattgtattatgatgaatttacctgatgtaggcgaatttggatctagtatgttggggaaatatgaacctgacatgattaagattgctctttcggtggctggtaatttaataagaaatcgagactacattcccgggagacgaggatatagctactacgtttacggtatagcctctagataa

### Isolate 2*

atgattgcgttattgatactatcgttaacgtgttcagtgtctacctatcgtctgcaaggatttaccaatgccggtatagtagcgtataaaaatattcaagatgataatattgtcttctcaccgtttggttattcgttttctatgtttatgtcgctattgcctgcatcaggtaatactagaatagaattattgaagactatggatttgagaaaaagagatctgggtccagcatttacagaattaatatcaggattagctaagctgaaaacatctaaatatacgtacactgatctaacttatcaaagtttcgtagataatactgtgtgcattaaaccgttgtattatcaacaatatcatagattcggcctatatagattaaactttagacgagatgcggttaataaaattaattctatagtagaacgtagatccggtatgtctaatgtagtagattctaatatgctcgacaataatactctatgggcaatcattaatactatatattttaaaggtacatggcaatatccgtttgatatcactaaaacacgcaatgctagttttactaataagtacggtacgaaaacggttcccatgatgaacgtagttactaaattgcaaggaaatacaatcacaatcgatgacgaagaatatgatatggtgcgccttccgtataaggatgctaatattagtatgtacctggcaataggtgataatatgacccatttcacagattctattacggctgcaaaattagactattggtcgtttcaattagggaataaagtgtacaatcttaaactccctaaattttctatcgaaaataagagggatattaagtcgatagccgaaatgatggctcctagtatgtttaatccagataatgcgtcgtttaaacatatgactagggacccattatatatttataaaatgtttcagaatgcaaagatagatgtcgacgaacaaggaactgtagcagaggcatctactatcatggtagctacggcgagatcatctcctgaaaaactggaatttaatacaccatttgtgttcatcattagacatgatattactggatttatattgtttatgggtaaggtagaatctccttaatattgtttatggatacggtggaaggaatcattattttatttatattgatgggtacgtgaaatctgaattttcttaataaatattatttttattaaatgtgtatatgttgttttgcgatagccatgtatctactaatcagatctattagagatattattaattctggtgcaatatgacaaaattataaaaaatgaaaaaatatacactaattagcgtctcgtttcagacatggatctgtcacgaattaatacttggaagtctaagcagctgaaaagctttctctctagtaaagatacatttaaggcggatgtccatggacatagtgccttgtattatgcaatagctgataataacgtgcgtctagtatgtacgttgttgaacgctggagcattgaaaaatcttctagagaatgaatttccattacatcaggcagccacattagaagataccaaaatagctttttcgcaacgggtttgccgccagaacacagctgaagcttcgaggggctcgcatctctccttcacgcgcccgccgccctacctgaggccgccatccacgccggttgagtcgcgttctgccgcctcccgcctgtggtgcctcctgaactgcgtccgccgtctagCTCTAAATGGTGAGCAAGGGCGAGGAGGATAACATGGCCATCATCAAGGAGTTCATGCGCTTCAAGGTGCACATGGAGGGCTCCGTGAACGGCCACGAGTTCGAGATCGAGGGCGAGGGCGAGGGCCGCCCCTACGAGGGCACCCAGACCGCCAAGCTGAAGGTGACCAAGGGTGGCCCCCTGCCCTTCGCCTGGGACATCCTGTCCCCTCAGTTCATGTACGGCTCCAAGGCCTACGTGAAGCACCCCGCCGACATCCCCGACTACTTGAAGCTGTCCTTCCCCGAGGGCTTCAAGTGGGAGCGCGTGATGAACTTCGAGGACGGCGGCGTGGTGACCGTGACCCAGGACTCCTCCCTGCAGGACGGCGAGTTCATCTACAAGGTGAAGCTGCGCGGCACCAACTTCCCCTCCGACGGCCCCGTAATGCAGAAGAAGACCATGGGCTGGGAGGCCTCCTCCGAGCGGATGTACCCCGAGGACGGCGCCCTGAAGGGCGAGATCAAGCAGAGGCTGAAGCTGAAGGACGGCGGCCACTACGACGCTGAGGTCAAGACCACCTACAAGGCCAAGAAGCCCGTGCAGCTGCCCGGCGCCTACAACGTCAACATCAAGTTGGACATCACCTCCCACAACGAGGACTACACCATCGTGGAACAGTACGAACGCGCCGAGGGCCGCCACTCCACCGGCGGCATGGACGAGCTGTACAAGTCTAGATCTGCCACCATGCTTGCATTTTGTTATTCGTTGCCCAATGCGGGcGATGTAATAAAGGGCAGAGTATACGAGAAGGATTATGCTCTATAcATTTATCTTTTTGACTATCCTCACTcTGAAGCTATCTTGGCAGAGAGTGTTAAGATGCATATGGATAGATATGTTGAATATAGGGATAAACTGGTAGGGAAAACTGTAAAAGTTAAAGTGATTAGAGTTGATTATACAAAAGGATATATAGATGTCAATTACAAAAGGATGTGTAGACATCAATAATAgGCGGCCGCCTAaacttgtttattgcagcttataatggttacaaataaagcaatagcatcacaaaAAAAAAAAAAAAAAAAAAAAAAAAAAAAAAAAAAAAAAAAAAAAAagaagataccaaaatagtaaagattttgctattcagtggaatggatgattcacaatttgatgacaaaggaaacaccgcattgtattatgcggttgatagtggtaacatgcaaacggtgaaactgtttgttaagaaaaattggagactgatgttctatgggaaaactggatggaaaacttcattttatcatgccgtcatgcttaatgatgtaagtattgtatcatactttctttcagaaataccatctacttttgatctggctattctccttagttgtattcacaccactataaaaaatggacacgtggatatgatgattctcttgctcgactatatgacgtcgacaaacaccaataattcccttctcttcattccggacattaaattggctatagataataaagacattgagatgttacaggctctgttcaaatacgacattaatatctactctgttaatctggaaaatgtactattggatgatgccgaaataactaagatgattatagaaaagcatgttgaatacaagtctgactcctatacaaaagatctcgatatcgtcaagaataataaattggatgaaataattagcaaaaacaaggaactcagactcatgtacgtcaattgtgtaaagaaaaactaa

### Isolate 3

atgaacaaacctaagacagattatgctggttatgcttgctgcgtaatatgcggtctaattgtcggaattatttttacagcgacactattaaaagttgtagaacgtaaattagttcatacaccatcaatagataaaacgataaaagatgcatatattagagaagattgtcctactgactggataagctataataataaatgtatccatttatctactgatcgaaaaacctgggaggaaggacgtaatacatgcaaagctctaaatccaaattcggatctaattaagatagagactccaaacgagttaagttttttaagaagccttagacgaggctattgggtaggagaatccgaaatattaaaccagacaaccccatataattttatagctaaaaatgccacgaagaatggaaatatatttgtagcacaacgaatactcccaaactgcattcgtgttacactatataacaattacactacatttttatcataccactacttcggttagatgttttagaaaaaaataaatatcgcGctttttcgcaacgggtttgccgccagaacacagctgaagcttcgaggggctcgcatctctccttcacgcgcccgccgccctacctgaggccgccatccacgccggttgagtcgcgttctgccgcctcccgcctgtggtgcctcctgaactgcgtccgccgtctagCTCCTGGGCAACGGTACCGGATCCCTGCAGAAGCTTCTAAAAATTGAAATTTTATTTTTTTTTTTTGGAATATAAATGGTGAGCAAGGGCGAGGAGGATAACATGGCCATCATCAAGGAGTTCATGCGCTTCAAGGTGCACATGGAGGGCTCCGTGAACGGCCACGAGTTCGAGATCGAGGGCGAGGGCGAGGGCCGCCCCTACGAGGGCACCCAGACCGCCAAGCTGAAGGTGACCAAGGGTGGCCCCCTGCCCTTCGCCTGGGACATCCTGTCCCCTCAGTTCATGTACGGCTCCAAGGCCTACGTGAAGCACCCCGCCGACATCCCCGACTACTTGAAGCTGTCCTTCCCCGAGGGCTTCAAGTGGGAGCGCGTGATGAACTTCGAGGACGGCGGCGTGGTGACCGTGACCCAGGACTCCTCCCTGCAGGACGGCGAGTTCATCTACAAGGTGAAGCTGCGCGGCACCAACTTCCCCTCCGACGGCCCCGTAATGCAGAAGAAGACCATGGGCTGGGAGGCCTCCTCCGAGCGGATGTACCCCGAGGACGGCGCCCTGAAGGGCGAGATCAAGCAGAGGCTGAAGCTGAAGGACGGCGGCCACTACGACGCTGAGGTCAAGACCACCTACAAGGCCAAGAAGCCCGTGCAGCTGCCCGGCGCCTACAACGTCAACATCAAGTTGGACATCACCTCCCACAACGAGGACTACACCATCGTGGAACAGTACGAACGCGCCGAGGGCCGCCACTCCACCGGCGGCATGGACGAGCTGTACAAGTCTAGATCTGCCACCATGCTTGCATTTTGTTATTCGTTGCCCAATGCGGGcGATGTAATAAAGGGCAGAGTATACGAGAAGGATTATGCTCTATAcATTTATCTTTTTGACTATCCTCACTcTGAAGCTATCTTGGCAGAGAGTGTTAAGATGCATATGGATAGATATGTTGAATATAGGGATAAACTGGTAGGGAAAACTGTAAAAGTTAAAGTGATTAGAGTTGATTATACAAAAGGATATATAGATGTCAATTACAAAAGGATGTGTAGACATCAATAATAgGCGGCCGCCTAaacttgtttattgcagcttataatggttacaaataaagcaatagcatcacaaaAAAAAAAAAAAAAAAAAAAAAAAAAAAAAAAAAAAAAAAAAAAAAagaaaaaaataaatatcgccgtaccgttcttgtttttataaaaataacaattaacaattatcaaattttttctttaatattttacgtggttgaccattcttggtggtaaaataatctcttagtgttggaatggaatgctgtttaatgtttccgcactcatcgtatattttgacgtatgcagtcacatcgtttacgcaatagtcagactgtagttctatcatgcttcctacatcagaaggaggaacagttttaaagtctcttggttttaatctattgccattagttttcatgaaatcctttgttttatccacttcacattttaaataaatgtccactatacattcttctgttaattttactagatcgtcatgggtcatagaatttataggttccgtagtccatggatccaaactagcaaacttcgcgtatacggtatcgcgattagtgtatacaccaacagtatgaaaattaagaaaacagtttaatagatcaacagaaatatttaatcctccgtttgatacagatgcgccatatttatggatttcggattcacacgttgtttgtctgaggggttcgtctagcgttgcttctacgtaaacttcgattcccatatattctttattgtcagaatcgcataccgatttatcatcatacactgtttgaaaactaaatggtatacacatcaaaataataaataataacgagtacat

### Isolate 4*

gagaaagagaaagagatagttagtctagatatttttcttagtacaaaagtcaatgttttaaaatatatggacaagaatttgtctgtataaaaacttgtgtgaaattttgtaccaaagaaaaaatgtgagcagtatcccctacatggattttactagatcatttatataccaaaaaatattatacgatctacgttttattatatgattttaacgtgtaaattataaacattattttatgatatacaattgtctggtaacctagatgggcataggggatgttgataagctcgacgagtatatgttgttggacgttattgtttaagaaatagttgatgcatcagaaagagaataaaaaatattttagtgagaccatcgaagagagaaagagataaaacttttttacgactccatcagaaagaggtttaatatttttgtgagaccatcgaagagagaaagagaataaaaatattttatgactccattgaagagagaaagagaaaatgagaatgagaataaaaatattttagtgacaccatcagaaagaggtttaatatttttgtgagaccatcgaagagagaaagagaataaaaatattttatgactccattgaagagagaaagagaaaatgagaatgagaataaaaatattttagtgacaccatcagaaagaggtttaatattttttatgagaccatcaaagagagaaagagaataaaaatatttttgtaaaactttttttatgagaccatcaaagagagaaagaGAATAAAAATATTTTTGTAAAACTTTCAAAGAGAGAAAGAGAATAAAAATATTTTTGTAAAACTTTTTTTATGAGACCATCAAAGAGAGAAAGAGAATAAAAATATTTTTGTAAAACTTTTTTTATGAGACCATCAAAGAGAGAAAGAGAATAAAAATATTTTTGTAAAACTTTTTTTATGAGACCATCAAAGAGAGAAAGAGAATAAAAATATTTTTGTAAAACTTTTTTTATGAGACCATCAGAAAGAGGTTTAATATTTTTGTGATACCCTGAAAGGAAATAGGAATAGGAATAGGAATAGTGTCATAATCGTATCACACTATTGAGACAGAAAAAGAAGAAGTAACGAGAGGTAACTTTTTGTGAATGTAGTTAAGAACATTTTTGTTTTGCAAACCGGAATATAGTGTCCGGTACACTTTTAGACCATCGAAGAGAGAAAGAGAATAAAAATATTTTATGACTCCATTGAAGAGAGAAAGAGAAAATGAGAATGAGAATAAAAATATTTTAGTctttttcgcaacgggtttgccgccagaacacagctgaagcttcgaggggctcgcatctctccttcacgcgcccgccgccctacctgaggccgccatccacgccggttgagtcgcgttctgccgcctcccgcctgtggtgcctcctgaactgcgtccgccgtctagCTCCTGGGCAACGGTACCGGATCCCTGCAGAAGCTTCTAAAAATTGAAATTTTATTTTTTTTTTTTGGAATATAAATGGTGAGCAAGGGCGAGGAGGATAACATGGCCATCATCAAGGAGTTCATGCGCTTCAAGGTGCACATGGAGGGCTCCGTGAACGGCCACGAGTTCGAGATCGAGGGCGAGGGCGAGGGCCGCCCCTACGAGGGCACCCAGACCGCCAAGCTGAAGGTGACCAAGGGTGGCCCCCTGCCCTTCGCCTGGGACATCCTGTCCCCTCAGTTCATGTACGGCTCCAAGGCCTACGTGAAGCACCCCGCCGACATCCCCGACTACTTGAAGCTGTCCTTCCCCGAGGGCTTCAAGTGGGAGCGCGTGATGAACTTCGAGGACGGCGGCGTGGTGACCGTGACCCAGGACTCCTCCCTGCAGGACGGCGAGTTCATCTACAAGGTGAAGCTGCGCGGCACCAACTTCCCCTCCGACGGCCCCGTAATGCAGAAGAAGACCATGGGCTGGGAGGCCTCCTCCGAGCGGATGTACCCCGAGGACGGCGCCCTGAAGGGCGAGATCAAGCAGAGGCTGAAGCTGAAGGACGGCGGCCACTACGACGCTGAGGTCAAGACCACCTACAAGGCCAAGAAGCCCGTGCAGCTGCCCGGCGCCTACAACGTCAACATCAAGTTGGACATCACCTCCCACAACGAGGACTACACCATCGTGGAACAGTACGAACGCGCCGAGGGCCGCCACTCCACCGGCGGCATGGACGAGCTGTACAAGTCTAGATCTGCCACCATGCTTGCATTTTGTTATTCGTTGCCCAATGCGGGcGATGTAATAAAGGGCAGAGTATACGAGAAGGATTATGCTCTATAcATTTATCTTTTTGACTATCCTCACTcTGAAGCTATCTTGGCAGAGAGTGTTAAGATGCATATGGATAGATATGTTGAATATAGGGATAAACTGGTAGGGAAAACTGTAAAAGTTAAAGTGATTAGAGTTGATTATACAAAAGGATATATAGATGTCAATTACAAAAGGATGTGTAGACATCAATAATAgGCGGCCGCCTAaacttgtttattgcagcttataatggttacaaataaagcaatagcatcacaaatttcacaaataaagcatttttttcactgcaAAAAAAAAAAAAAAAAAAAAAAAAAAAAAAAAAAAAAAAAAAAAAAAAtattttagtgacaccatcagaaagaggtttaatatttttgtgagaccatcgaagagagaaagagaataaaaatatttttgtaaaactttttttatgagaccatcaaagagagaaagagaataaaaatatttttgtaaaactttttttatgagaccatcaaagagagaaagagaataaaaatatttttgtaaaactttttttatgagaccatcaaagagagaaagagaataaaaatatttttgtaaaactttttttatgagaccatcaaagagagaaagagaataaaaatatttttgtaaaactttttttatgagaccatcaaagagagaaagagaataaaaatatttttgtaaaactttttttatgagaccatcaaagagagaaagagaataaaaatatttttgtaaaactttttttatgagaccatcagaaagaggtttaatatttttgtgataccctgaaaggaaataggaataggaataggaatagtgtcataatcgtatcacactattgagacagaaaaagaagaagtaacgagaggtaactttttgtgaatgtagttaagaacatttttgttttgcaaaccggaatatagtgtccggtacacttttttaattcgtggtgtgcctgaatcgttcgattaaccctactcatccaatttcagatgaatagagttatcgattcagacacacgctttgagttttgttgaatcgatgagtgaagtatcatcggttgcaccttcagatgccgatccgtcgacatacttgaatccatccttgacctcaagttcagatgattccttgcacatgtctccgatacgaacgctaaactctagattcttgacacattttgtatcgacgatcgttgaaccgatgatatcttcgtaactcactttcttatgagagatgttagacccgagtactggatgggtcttgatgtcgctgtctttctcttcttcgctacatctgatgtcgatagacacctcacagtctttgatcatagccagagcttcttcacgagtgatcgcgggagagtccttaccttgtcctggggacacgctggacaatctagcattcactgtgtttccatcagcggattctgagatggatttaatctgaggacatttggtgaatccaaagttcattctcagacctccaccgatgatggagtaataagtggtaggaggatctacatcctcgactgatgtggaatcatcttctgattccacctcgggatctggatctgactcggactctgtaatttccgttacggattggcaaatcttatcattggtcggtgtttggtcttgctttgtgactttgataataacatcgattcccatatgatgtttgttttcttcttccgtacacgaggaggaggatgaggatgattgctgaagactggcaggcacatgcat

### Isolate 5

ttatagtataaagtaataaaaaatagttaatgtgatgactagcgccaccaacgccaacaacatttgataatttctacttactagacgtaccgtaaaaatataaattactataacaaataatagtatatcaataaacaacctaattaatggtcgaagtatagcaggacattgatgctctagaccgtgtataacaaaatctacaaatttttcatccgctatattttgtttcactatatcgtctagacgatcagcgataacttccatgttaatctattaaaatattatcaatatattttcagttttgcatatccgtggtagcaataaccatcggagaagttctaaagaatgtgtccatgtaagtagtccaatggacgttttccttattggctaaaataagtttgattttatcattggtggatgtgaacagcatacgcttggcatagtacataaacaacgctgccaatattataacaccgataacaatcatataaaactgaactcctgtaccagcaacttgtctaggtgctatttgagtagtggccttagtagtcaattgcatcaacgccttaatggcacaatttcctttgctagatcctgtattaataaattccaaatttgttggagatcctggggctccgtaacattcatctatgattacgttttgtatctttaatttgttatcgacgaccgcgctagaattacaagtttgttttacataattttcaaaatctctaacaacagtgtttacactcgtctgaatgtttaacgcagcagtaaacatagctggcacgtatgctttttgttccggtgttaatccactatatgtttctgtagcggctgataacacagcatccaactgagcatccgcgtccgcagagcacatatttttaacagtgaggttacatccatggttttgtcggatataaaaatttccgatttctatatcacattttgtttgagcactagcattcgcttcttgttctaatttagacgagatacgttcgctgagtgtattcaccgtcgtctgtatgcttgctgcggcacccatttaaatagctacaattagtatccatattaccaagagagataataaactgatcaaatgcaattttaggtcgaacgagtgtttaatattatgttgaactacttttgctctattgacgggaacagtagaaaatctatcactattgcttaatccacatgacaatcttaatgaggaagttttatccatctgtaagttgttcacgctagtattacatcgtacaatattgcaaagtcctaaattattataattacgtgttagcaagaaattaacattggcattcgaacactctggatcccaacattctcgaggttccgcatattttaatgactcttctaacttatctctagtgggataactacatctcatatatttctgtttaaagtccgcagactgttgtcttagaatataatcgatcatctctttgctatcttctgtattgtgtgcgcgtaaatgatgcaaaaatgattcacatattggtacactagcatctttactacataatgtttgcatcttattatacagatttattaacgattgttgaccctctacagttctattactcctattaaaggctgaaccgatccactgatggcatatgtttctatcgaacgtatccccctgacaccagtcgaataaatcaacatcgcattttccagtgtcgtgaacgtctggccaacacgattctaatactgcaccctcttcatacttatctgatatctttccatcctttttccaataatgagtacgattaaaagtgcgacagcaattgggtgcggttccattatacgattgtcttaacaatgccgaaagaccaccgggtcctgtgttgactaatctaaactctggatatcttttctttactagctttttcgcaacgggtttgccgccagaacacagctgaagcttcgaggggctcgcatctctccttcacgcgcccgccgccctacctgaggccgccatccacgccggttgagtcgcgttctgccgcctcccgcctgtggtgcctcctgaactgcgtccgccgtctagCTCCTGGGCAACGGTACCGGATCCCTGCAGAAGCTTCTAAAAATTGAAATTTTATTTTTTTTTTTTGGAATATAAATGGTGAGCAAGGGCGAGGAGGATAACATGGCCATCATCAAGGAGTTCATGCGCTTCAAGGTGCACATGGAGGGCTCCGTGAACGGCCACGAGTTCGAGATCGAGGGCGAGGGCGAGGGCCGCCCCTACGAGGGCACCCAGACCGCCAAGCTGAAGGTGACCAAGGGTGGCCCCCTGCCCTTCGCCTGGGACATCCTGTCCCCTCAGTTCATGTACGGCTCCAAGGCCTACGTGAAGCACCCCGCCGACATCCCCGACTACTTGAAGCTGTCCTTCCCCGAGGGCTTCAAGTGGGAGCGCGTGATGAACTTCGAGGACGGCGGCGTGGTGACCGTGACCCAGGACTCCTCCCTGCAGGACGGCGAGTTCATCTACAAGGTGAAGCTGCGCGGCACCAACTTCCCCTCCGACGGCCCCGTAATGCAGAAGAAGACCATGGGCTGGGAGGCCTCCTCCGAGCGGATGTACCCCGAGGACGGCGCCCTGAAGGGCGAGATCAAGCAGAGGCTGAAGCTGAAGGACGGCGGCCACTACGACGCTGAGGTCAAGACCACCTACAAGGCCAAGAAGCCCGTGCAGCTGCCCGGCGCCTACAACGTCAACATCAAGTTGGACATCACCTCCCACAACGAGGACTACACCATCGTGGAACAGTACGAACGCGCCGAGGGCCGCCACTCCACCGGCGGCATGGACGAGCTGTACAAGTCTAGATCTGCCACCATGCTTGCATTTTGTTATTCGTTGCCCAATCCTCACTcTGAAGCTATCTTGGCAGAGAGTGTTAAGATGCATATGGATAGATATGTTGAATATAGGGATAAACTGGTAGGGAAAACTGTAAAAGTTAAAGTGATTAGAGTTGATTATACAAAAGGATATATAGATGTCAATTACAAAAGGATGTGTAGACATCAATAATAgGCGGCCGCCTAaacttgtttattgcagcttataatggttacaaataaagcaatagcatcacaaatttcacaaataaagcatttttttcactgcaAAAAAAAAAAAAAAAAAAAAAAAAAAAAAAAAAAAAAAAAAAAacttgtctaggtgctatttgagtagtggccttagtagtcaattgcatcaacgccttaatggcacaatttcctttgctagatcctgtattaataaattccaaatttgttggagatcctggggctccgtaacattcatctatgattacgttttgtatctttaatttgttatcgacgaccgcgctagaattacaagtttgttttacataattttcaaaatctctaacaacagtgtttacactcgtctgaatgtttaacgcagcagtaaacatagctggcacgtatgctttttgttccggtgttaatccactatatgtttctgtagcggctgataacacagcatccaactgagcatccgcgtccgcagagcacatatttttaacagtgaggttacatccatggttttgtcggatataaaaatttccgatttctatatcacattttgtttgagcactagcattcgcttcttgttctaatttagacgagatacgttcgctgagtgtattcaccgtcgtctgtatgcttgctgcggcacccatttaaatagctacaattagtatccatattaccaagagagataataaactgatcaaatgcaattttaggtcgaacgagtgtttaatattatgttgaactacttttgctctattgacgggaacagtagaaaatctatcactattgcttaatccacatgacaatcttaatgaggaagttttatccatctgtaagttgttcacgctagtattacatcgtacaatattgcaaagtcctaaattattataattacgtgttagcaagaaattaacattggcattcgaacactctggatcccaacattctcgaggttccgcatattttaatgactcttctaacttatctctagtgggataactacatctcatatatttctgtttaaagtccgcagactgttgtcttagaatataatcgatcatctctttgctatcttctgtattgtgtgcgcgtaaatgatgcaaaaatgattcacatattggtacactagcatctttactacataatgtttgcatcttattatacagatttattaacgattgttgaccctctacagttctattactcctattaaaggctgaaccgatccactgatggcatatgtttctatcgaacgtatccccctgacaccagtcgaataaatcaacatcgcattttccagtgtcgtgaacgtctggccaacacgattctaatactgcaccctcttcatacttatctgatatctttccatcctttttccaataatgagtacgattaaaagtgcgacagcaattgggtgcggttccattatacgattgtcttaacaatgccgaaagaccaccgggtcctgtgttgactaatctaaactctggatatcttttctttactttatcctctttatcttttgctagtggtcctatatgcacatattctaaaagcttagcgggagctatcacgtcatgcattttatccacgtttaataacatctcatcagtgggtactcccggaggcggatcccgtttagggagctcaacacttactccgccacccatatttatctcattgaaagtattaatctaaaaacgccataaagatgttgatcttaaaggattgaactctatccgaaaacaacattcctagaatgttatcgtcattatccattacgattctagtttcaaaaacattgactctctttttgaatcctcgtagtttgttgagagacgagatagctattttgaaagaaaacttttgtagttcttgagaacattcagtcatagaatattccctggaaaatgcatcagtattaccaggagtcttcataataatattgtcatctttaaacataatagccagatgctgatgctgactaatacacttgataaagcccaacaaccattctaaatgaatgacggttctaccacaacatttttcttcatacttgtgaaaattaaacacataagatttcttttgatctatacttagacaaatagtagtatctgtcctaataggcatcagttccttattacaatcgacacttactacatgataactagaaagttttactagattattttccagatcaggttctatatctatgatggcatcattgtgaaaactacacaaacacgattttaccttggacacaggaagattaaacacaatgttttcggctcctcggtagaacactgatgcactgagaggtataatggcccaaatgtttacagatccgcccaaggcggcaaaaatatacattaactcatccgtcgagtctacatttatagacacttcttcactgaactctgaaaaatatgccacaatttggcgcagtttatcgatttttatacggatgctcattttaaatttttgtaaattatttaaagttaaatggctgcagaacagcgtcgttctacaatttttgacatagtttcaaaatgtatagtgcaatctgtattaagagatatatctattaattctgaatacatagagtccaaagctaaacaattgtgctattgtccggcatcgaaaaaggaatcagtgattaatggtatctacaattgttgcgagtcaaatatagaaataatggacaaagagcagctattaaaaatattggacaatcttcgatgtcattcggctcatgtatgtaacgccacagatttctggagactatataattcgttaaaacggtttactcatactaccgcattctttaatacatgcaagcccactattctagccacg

### Isolate 6

atgccttctttgttctcctccctactaacgaccttagttttccatattttgatttattatcaaattaatttagtaactgtaaatataattatgaattgtttccaagaaaaacaattttcaagagaaaatttattaaaaatgccgtttagaatggttttaacgggaggatctggatctggaaaaactatctatttactatctctgttttctacactagttaaaaaatataaacatatattcttgtttacacccgtttataacccagattatgatggatacatttggccaaatcatattaatttcgttagtagtcaggaatctctagaatataatctgatacgaactaaaagtaacatagaaaaatgtattgctgtcgcacaaaatcataaaaaatcagcacactttttacttatttttgatgatgtaggcgataaactatcaaaatgcaatactctaatagaattcttaaactttggaaggcatttaaacacgtctattattctactatgccaaacttatagacacgtaccaatattaggacgggctaacattacgcatttttgtagttttaacatttccatctcagacgcggaaaatatgctacgatcgatgcctgtaaaggggaaacgaaaggatatattaaacatgttgaatatgatacagacagctagatccaataatcgattggctattattatcgaagactccgtattttgtgaaggtgaattacgtatatgtaccgataccgccgataaggacgtcatagaacaaaagttaaacatcgatattttagtaaatcaatattcgcacatgaaaaagaatctaaacgctatattagaaagtaaaaaaacctttttcgcaacgggtttgccgccagaacacagctgaagcttcgaggggctcgcatctctccttcacgcgcccgccgccctacctgaggccgccatccacgccggttgagtcgcgttctgccgcctcccgcctgtggtgcctcctgaactgcgtccgccgtctagCTCCTGGGCAACGGTACCGGATCCCTGCAGAAGCTTCTAAAAATTGAAATTTTATTTTTTTTTTTTGGAATATAAATGGTGAGCAAGGGCGAGGAGGATAACATGGCCATCATCAAGGAGTTCATGCGCTTCAAGGTGCACATGGAGGGCTCCGTGAACGGCCACGAGTTCGAGATCGAGGGCGAGGGCGAGGGCCGCCCCTACGAGGGCACCCAGACCGCCAAGCTGAAGGTGACCAAGGGTGGCCCCCTGCCCTTCGCCTGGGACATCCTGTCCCCTCAGTTCATGTACGGCTCCAAGGCCTACGTGAAGCACCCCGCCGACATCCCCGACTACTTGAAGCTGTCCTTCCCCGAGGGCTTCAAGTGGGAGCGCGTGATGAACTTCGAGGACGGCGGCGTGGTGACCGTGACCCAGGACTCCTCCCTGCAGGACGGCGAGTTCATCTACAAGGTGAAGCTGCGCGGCACCAACTTCCCCTCCGACGGCCCCGTAATGCAGAAGAAGACCATGGGCTGGGAGGCCTCCTCCGAGCGGATGTACCCCGAGGACGGCGCCCTGAAGGGCGAGATCAAGCAGAGGCTGAAGCTGAAGGACGGCGGCCACTACGACGCTGAGGTCAAGACCACCTACAAGGCCAAGAAGCCCGTGCAGCTGCCCGGCGCCTACAACGTCAACATCAAGTTGGACATCACCTCCCACAACGAGGACTACACCATCGTGGAACAGTACGAACGCGCCGAGGGCCGCCACTCCACCGGCGGCATGGACGAGCTGTACAAGTCTAGATCTGCCACCATGCTTGCATTTTGTTATTCGTTGCCCAATCCTCACTcTGAAGCTATCTTGGCAGAGAGTGTTAAGATGCATATGGATAGATATGTTGAATATAGGGATAAACTGGTAGGGAAAACTGTAAAAGTTAAAGTGATTAGAGTTGATTATACAAAAGGATATATAGATGTCAATTACAAAAGGATGTGTAGACATCAATAATAgGCGGCCGCCTAaacttgtttattgcagcttataatggttacaaataaagcaatagcatcacaaaAAAAAAAAAAAAAAAAAAAAAAAAAAAAAAAAAAAAAAAAAAAAAagaaagtaaaaaaacaaaattgtgcaatagcgatcaatcatcgtcgtcaaaaaatgtatcatcataattattaacttttgtaacaatagtcctatttagagaaagtctatcgatagacgatcccaatttgtaaattactccactacttgtcactccatgatatgataaatccatgtaaaatagcatcatctttagatcattaattgttaccttccccaatacaaccaaatcatcatgatatatacctcctccagacaagtatttaacaacggtagaatgctttggcttataaaatacaaatgacattcccttatgtttaatcttaatcttttctttagttattgaatcgttacaattataaaatgatgttttttccaaaaacctaagtgtatttaaaatagatgccat

### Isolate 7

atggagggatctaaacgcaaacacgatagtcggcgaccacaacaagaacaggagcagcctcgtccacgtacaccgccatcatatgaagaaattgcaaaatatggacactcatttaacgtgaaaagatttacgaatgaagaaatgtgtcttaagaatgattatccacgaGctttttcgcaacgggtttgccgccagaacacagctgaagcttcgaggggctcgcatctctccttcacgcgcccgccgccctacctgaggccgccatccacgccggttgagtcgcgttctgccgcctcccgcctgtggtgcctcctgaactgcgtccgccgtctagCTCCTGGGCAACGGTACCGGATCCCTGCAGAAGCTTCTAAAAATTGAAATTTTATTTTTTTTTTTTGGAATATAAATGGTGAGCAAGGGCGAGGAGGATAACATGGCCATCATCAAGGAGTTCATGCGCTTCAAGGTGCACATGGAGGGCTCCGTGAACGGCCACGAGTTCGAGATCGAGGGCGAGGGCGAGGGCCGCCCCTACGAGGGCACCCAGACCGCCAAGCTGAAGGTGACCAAGGGTGGCCCCCTGCCCTTCGCCTGGGACATCCTGTCCCCTCAGTTCATGTACGGCTCCAAGGCCTACGTGAAGCACCCCGCCGACATCCCCGACTACTTGAAGCTGTCCTTCCCCGAGGGCTTCAAGTGGGAGCGCGTGATGAACTTCGAGGACGGCGGCGTGGTGACCGTGACCCAGGACTCCTCCCTGCAGGACGGCGAGTTCATCTACAAGGTGAAGCTGCGCGGCACCAACTTCCCCTCCGACGGCCCCGTAATGCAGAAGAAGACCATGGGCTGGGAGGCCTCCTCCGAGCGGATGTACCCCGAGGACGGCGCCCTGAAGGGCGAGATCAAGCAGAGGCTGAAGCTGAAGGACGGCGGCCACTACGACGCTGAGGTCAAGACCACCTACAAGGCCAAGAAGCCCGTGCAGCTGCCCGGCGCCTACAACGTCAACATCAAGTTGGACATCACCTCCCACAACGAGGACTACACCATCGTGGAACAGTACGAACGCGCCGAGGGCCGCCACTCCACCGGCGGCATGGACGAGCTGTACAAGTCTAGATCTGCCACCATGCTTGCATTTTGTTATTCGTTGCCCAATGCGGGcGATGTAATAAAGGGCAGAGTATACGAGAAGGATTATGCTCTATAcAATGTTGAATATAGGGATAAACTGGTAGGGAAAACTGTAAAAGTTAAAGTGATTAGAGTTGATTATACAAAAGGATATATAGATGTCAATTACAAAAGGATGTGTAGACATCAATAATAgGCGGCCGCCTAaacttgtttattgcagcttataatggttacaaataaagcaatagcatcacaaaAAAAAAAAAAAAAAAAAAAAAAAAAAAAAAAAAAAAAAAAAAAAAgaatgattatccacgaattatatcatataatcctccaccaaaatagagaatatatatatcatcatttcatgatgtatactactgacatagtttcaatgtgaacttttcactttcttgccggttatgaagaatattttttattttaatggtcattactaatcgtatattataattgaaaatgaattagtttaatatgacgctcgtcatgggattctgctgtggtagattctgtgacgctaagaataagaataagaataagaataagaataagaataagaataagaataagaaggaagatgtagaagagggaagagaaggatgttacaattataagaaccttaatgatctggatgaatccgaagcacgtgtagaatttggaccattatatatgataaatgaagaaaaatcagacataaatacattggatataaaaagaagatatagacacacgatagagtctgtatatttctaa

### Isolate 8

atgatgacaccagaaaacgacgaagagcagacatctgtgttctccgctactgtttacggagacaaaattcaaggaaagaataaacgcaaacgcgtgattggtctatgtattagaatatctatggttatttcactactatctatgattaccatgtccgcgtttctcatagtgcgcctaaatcaatgcatgtctgctaacgaggctgctattactgacgccgctgttgccgttgctgctgcatcatctactcatagaaaggttgcgtctactttttcgcaacgggtttgccgccagaacacagctgaagcttcgaggggctcgcatctctccttcacgcgcccgccgccctacctgaggccgccatccacgccggttgagtcgcgttctgccgcctcccgcctgtggtgcctcctgaactgcgtccgccgtctagCTCCTGGGCAACGGTACCGGATCCCTGCAGAAGCTTCTAAAAATTGAAATTTTATTTTTTTTTTTTGGAATATAAATGGTGAGCAAGGGCGAGGAGGATAAACATGGCCATCATCAAGGAGTTCATGCGCTTCAAGGTGCACATGGAGGGCTCCGTGAACGGCCACGAGTTCGAGATCGAGGGCGAGGGCGAGGGCCGCCCCTACGAGGGCACCCAGACCGCCAAGCTGAAGGTGACCAAGGGTGGCCCCCTGCCCTTCGCCTGGGACATCCTGTCCCCTCAGTTCATGTACGGCTCCAAGGCCTACGTGAAGCACCCCGCCGACATCCCCGACTACTTGAAGCTGTCCTTCCCCGAGGGCTTCAAGTGGGAGCGCGTGATGAACTTCGAGGACGGCGGCGTGGTGACCGTGACCCAGGACTCCTCCCTGCAGGACGGCGAGTTCATCTACAAGGTGAAGCTGCGCGGCACCAACTTCCCCTCCGACGGCCCCGTAATGCAGAAGAAGACCATGGGCTGGGAGGCCTCCTCCGAGCGGATGTACCCCGAGGACGGCGCCCTGAAGGGCGAGATCAAGCAGAGGCTGAAGCTGAAGGACGGCGGCCACTACGACGCTGAGGTCAAGACCACCTACAAGGCCAAGAAGCCCGTGCAGCTGCCCGGCGCCTACAACGTCAACATCAAGTTGGACATCACCTCCCACAACGAGGACTACACCATCGTGGAACAGTACGAACGCGCCGAGGGCCGCCACTCCACCGGCGGCATGGACGAGCTGTACAAGTCTAGATCTGCCACCATGCTTGCATTTTGTTATTCGTTGCCCAATGCGGGcGATGTAATAAAGGGCAGAGTATACGAGAAGGATTATGCTCTATAcATTTATCTTTTTGACTATCCTCACTcTGAAGCTATCTTGGCAGAGAGTGTTAAGATGCATATGGATAGATATGTTGAATATAGGGATAAACTGGTAGGGAAAACTGTAAAAGTTAAAGTGATTAGAGTTGATTATACAAAAGGATATATAGATGTCAATTACAAAAGGATGTGTAGACATCAATAATAgGCGGCCGCCTAaacttgtttattgcagcttataatggttacaaataaagcaatagcatcacaaaAAAAAAAAAAAAAAAAAAAAAAAAAAAAAAAAAAAAAAAAAAAagaaaggttgcgtctagcactacgcaatatgatcacaaagaaagctgtaatggtttatattaccagggttcttgttatatattacattcagactaccagttattctcggatgctaaagcaaattgcactgcggaatcatcaacactacccaataaatccgatgtcttgattacctggctcattgattatgttgaggatacatggggatctgatggtaatccaattacaaaaactacatccgattatcaagattctgatgtatcacaagaagttagaaagtatttttgtgttaaaacaatgaactaa

### Isolate 9

gtaaaattaaattaattataaaattatgtatatgatttactaactttagttagataagttagtaatacataaattttagtatattaatattatattttaaatattttatttagtgtctagaaaaaaatgtgtgaccaacgaccgtaggaaactctagagggtaagaaaaatcaatcgcGctttttcgcaacgggtttgccgccagaacacagctgaagcttcgaggggctcgcatctctccttcacgcgcccgccgccctacctgaggccgccatccacgccggttgagtcgcgttctgccgcctcccgcctgtggtgcctcctgaactgcgtccgccgtctaATATAAATGGTGAGCAAGGGCGAGGAGGATAACATGGCCATCATCAAGGAGTTCATGCGCTTCAAGGTGCACATGGAGGGCTCCGTGAACGGCCACGAGTTCGAGATCGAGGGCGAGGGCGAGGGCCGCCCCTACGAGGGCACCCAGACCGCCAAGCTGAAGGTGACCAAGGGTGGCCCCCTGCCCTTCGCCTGGGACATCCTGTCCCCTCAGTTCATGTACGGCTCCAAGGCCTACGTGAAGCACCCCGCCGACATCCCCGACTACTTGAAGCTGTCCTTCCCCGAGGGCTTCAAGTGGGAGCGCGTGATGAACTTCGAGGACGGCGGCGTGGTGACCGTGACCCAGGACTCCTCCCTGCAGGACGGCGAGTTCATCTACAAGGTGAAGCTGCGCGGCACCAACTTCCCCTCCGACGGCCCCGTAATGCAGAAGAAGACCATGGGCTGGGAGGCCTCCTCCGAGCGGATGTACCCCGAGGACGGCGCCCTGAAGGGCGAGATCAAGCAGAGGCTGAAGCTGAAGGACGGCGGCCACTACGACGCTGAGGTCAAGACCACCTACAAGGCCAAGAAGCCCGTGCAGCTGCCCGGCGCCTACAACGTCAACATCAAGTTGGACATCACCTCCCACAACGAGGACTACACCATCGTGGAACAGTACGAACGCGCCGAGGGCCGCCACTCCACCGGCGGCATGGACGAGCTGTACAAGTCTAGATCTGCCACCATGCTTGCATTTTGTTATTCGTTGCCCAATGCGGGTGATGTAATAAAGGGCAGAGTATACGAGAAGGATTATGCTCTATATAATGTTGAATATAGGGATAAACTGGTAGGGAAAACTGTAAAAGTTAAAGTGATTAGAGTTGATTATACAAAAGGATATATcGATGTCAATTACAAAAGGATGTGTAGACATCAATAATAgGCGGCCGCCTAaacttgtttattgcagcttataatggttacaaataaagcaatagcatcacaaaAAAAAAAAAAAAAAAAAAAAAAAAAAAAAAAAaaaaatcaatcgctttatagagaccatcagaaagaggtttaatatttttgtgagaccatcgaaggagaaagagataaaacttttttacgactccatcagaaagaggtttaatatttttgtgagaccatcgaaggagaaagagataaaacttttttacgactccatcagaaagaggtttaatatttttgtgagaccatcgaaggagaaagagataaaacttttttacgactccatcagaaagaggtttaatatttttgtgagaaaggagaaagagataaaacttttttacgactccatcagaaagaggtttaatatttttgtgagaccatcgaaggagaaagagataaaacttttttacgactccatcagaaagaggtttaatatttttgtgagaccatcgaaggagaaagagataaaacttttttacgactccatcagaaagaggtttaatatttttgtgagaccatcgaaggagaaagagataaaacttttttacgactccatcagaaagaggtttaatatttttgt

### Isolate 10

atggatatctttaaagaactaatcgtaaaacaccctgatgaaaatgttttgatttctccagtttctattttatctactttatctattctaaatcatggagcagctggttctacagctgaacaactatcaaaatatatagagaatatgaatgagaatacacccgatgacaataatgacatggacgtagatattccgtattgtgcgacactagctaccgcaaataaaatatacggtagcgatagtatcgagttccacgcctccttcctacaaaaaataaaagacgattttcaaactgtaaactttaataatgctaaccaaacaaaggaactaatcaacgaatgggttaagacaatgacaaatggtaaaattaattccttattgactagtccgctatccattaatactcgtatgacagttgttagcgccgtccattttaaagcaatgtggaaatatccattttctaaacatcttacatatacagacaagttttatatttctaagaatatagttaccagtgttgatatgatggtgggtaccgagaataacttgcaatatgtacatattaatgaattattcggaggattctctattatcgatattccatacgagggaaactctagtatggtgattatactaccggacgacatagaaggtatatataacatagaaaaaaatataacagatgaaaaatttaaaaaatggtgtggtatgttatctactaaaagtatagacttgtatatgccaaagtttaaagtggaaatgacagaaccgtataatctggtaccgattttagaaaatttaggacttactaatatattcggatattatgcagattttagcaagatgtgtaatgaaactatcactgtagaaaaatttctacatacgacgtttatagatgttaatgaggagtatacagaagcatcggccgttacaggagtatttacgattaacttttcgatggtatatcgtacgaaggtctacataaaccatccattcatgtacatgattaaagacaccacaggacgtatactttttatagggaaatactgctatccgcaataaatataaacaaatagacttttatcacgtttatctatgtctaaatattacaaatagtaatagtataaactaaagctgataatacttaaaaaaataataatatcatttacaattaatagtataaactaaaaattaaacaaatcgttattataagtaatatcaaaatgatgatatacggattaatagcgtgtcttatattcgtgacttcatccatcgctagtccacccattacggaagataaatcgttcaatagtgtagaggtattagtttccttgtttagagatgaccaaaaagactatacggtaacttctcagttcaataactacactatcgataccaaagactggactatccacacctgatggtttggatataccattgactaatataacttattggtcacggtttactataggtcgtgcattgttcaaatcagagtctgaggatattttccaaaagaaaatgagtattctaggtgtttctatagaatgtaagaagtcgtcgacattacttacttttttgaccgtgcgtaaaatgactcgagtatttaataaatttccagatatggcttattatcgaggagactgtttaaaagccgtttatgtaacaatgacttataaaaatactaaaactggagagactgattacacgtacctctctaatgggggttgcctgcatactatcgtaatggggtcgatggttgattattgattagtatattcctgctttttcgcaacgggtttgccgccagaacacagctgaagcttcgaggggctcgcatctctccttcacgcgcccgccgccctacctgaggccgccatccacgccggttgagtcgcgttctgccgcctcccgcctgtggtgcctcctgaactgcgtccgccgtctagCTCCTGGGCGTGAGCAAGGGCGAGGAGGATAACATGGCCATCATCAAGGAGTTCATGCGCTTCAAGGTGCACATGGAGGGCTCCGTGAACGGCCACGAGTTCGAGATCGAGGGCGAGGGCGAGGGCCGCCCCTACGAGGGCACCCAGACCGCCAAGCTGAAGGTGACCAAGGGTGGCCCCCTGCCCTTCGCCTGGGACATCCTGTCCCCTCAGTTCATGTACGGCTCCAAGGCCTACGTGAAGCACCCCGCCGACATCCCCGACTACTTGAAGCTGTCCTTCCCCGAGGGCTTCAAGTGGGAGCGCGTGATGAACTTCGAGGACGGCGGCGTGGTGACCGTGACCCAGGACTCCTCCCTGCAGGACGGCGAGTTCATCTACAAGGTGAAGCTGCGCGGCACCAACTTCCCCTCCGACGGCCCCGTAATGCAGAAGAAGACCATGGGCTGGGAGGCCTCCTCCGAGCGGATGTACCCCGAGGACGGCGCCCTGAAGGGCGAGATCAAGCAGAGGCTGAAGCTGAAGGACGGCGGCCACTACGACGCTGAGGTCAAGACCACCTACAAGGCCAAGAAGCCCGTGCAGCTGCCCGGCGCCTACAACGTCAACATCAAGTTGGACATCACCTCCCACAACGAGGACTACACCATCGTGGAACAGTACGAACGCGCCGAGGGCCGCCACTCCACCGGCGGCATGGACGAGCTGTACAAGTCTAGATCTGCCACCATGCTTGCATTTTGTTATTCGTTGCCCAATGCGGGcGATGTAATAAAGGGCAGAGTATACGAGAAGGATTATGCTCTATAcATTTATCTTTTTGACTATCCTCACTcTGAAGCTATCTTGGCAGAGAGTGTTAAGATGCATATGGATAGATATGTTGAATATAGGGATAAACTGGTAGGGAAAACTGTAAAAGTTAAAGTGATTAGAGTTGATTATACAAAAGGATATATAGATGTCAATTACAAAAGGATGTGTAGACATCAATAATAgGCGGCCGCCTAaacttgtttattgcagcttataatggttacaaataaagcaatagcatcacaaaATTCACAAATAAAGCATTTTTTCACTGCAAAAAAAAAAAAAAAAAAAAAAAAAAAaagagtaacagctgcccccattcttaataatcgtcagtatttaaactgttaaatgttggtatatcaacatctaccttatttcccgcagtataaggtttgttgcaggtatactgttcaggaatggttacatttatacttcttctatagtcctgtctttcgatgttcatcacatatgcaaagaacagaataaacaaaataatgtaagaaataatattaaatatctgtgaattcgtaaatacattgattgccataataattacagcagctacaatacacacaatagacattcccacagtgttgccattacctccacgatacatttgagttactaagcaataggtaataactaagctagtaagaggcaatagaaaagatgagataaatatcatcaatatagagattagaggagggctatatagagccaagacgaacaaaatcaaaccgagtaacgttctaacatcattatttttgaagattcccaaataatcattcattcctccataatcgttttgcatcatacctccatctttaggcataaacgattgctgctgttcctctgtaaataaatctttatcaagcactccagcacccgcagagaagtcgtcaagcatattgtaatatcttaaataactcat

*Note that some isolate 4 viruses carry a second insertion that precisely matches that of the isolate 2 insert. We believe that this is a result of recombination between two separate strains during plaque purification of both.

Nanopore reads showing K3L-CNV:

### Isolate 4

TGTTGATAATGGGAAGATGGGATATATGATGTTGCATGAACGTTGATAGTTCACTGGTCATCAATATGCCGACATTCAATGCCCAGAATCATGGTTAATTTCGCCAATACATACCACGATACTTCCAGCGTGAGAAGTCGCGTAAACAATATCCTCATCTCCTCTGCGGACGCTATTAATTGTGATAGTTGAATACCAGCCAGCGCTTGTAATTTCGGCGTCGTTAGCGCAGTCACTGGTTTGAATGATAGCAGCAACGTCAACATATTTCATCATTTTATGAGGATTCGTGCTATGCCAGCCAGCGTGTGTTTATAACAATATTTCGTAGCCCATATTATTAATTCAGTAATATAAACTATCAATATTGTGTTACTGTAAAATGTCATGCGGTCAAATGTTATTAGTCGTACACGCATGCCAATATAATATAGTCGTAATTATGCGCAACACGGAAGATATCACCATTTGTCATTGTCGCTTCATCATATCTTGAGCTTAGAACATCATCAATCCATTTCATGCCGGTGACACGGTATGCACAAGCGATTGATTGCACGCAATACAACAATGAAGTATGAATACGGGGTACAAACAACCGAAATTTCAACATTACTATTTATGATATACACGTTATGTCGCCATTGCTTCATACCATGGATTGTGAATCAATCGTAAACAATTTTATGGCATCGATAACGACGAGTCATTTATGCAGGACGCGCAAGATATTCCGTACAGCATCATACGCAACAAAGCGTACGGGCGAAATAAAAGACATTTTCAACCGGAACTTTACACGCTACCAAACAAAGTGACCAATCAATCTAGCGGTAAAAGTCCGCCATTCACGTATGACAGCCAAAGGCGCCGCCCATTTTGGGTACGCTTTCAAACAGCCCACATAGTACAGACAAGTTTTTATATTTCATTAACATGGTTGCGAAGGCCTCGTGGGTCATGCAGTTGAGTATTACGACTGAGTCAGTGACACTTTCATTATCGATACCATGAACTTCAGTATACGATTATTTATGTAGCTGCGATCAAGATGAAAAATTTGCGTCGGTGCGGTATGTTAGACTTCGTTAAAAGCTTGGAAAGTGAAATGACAGAATCGCATGGGCGTCGCATTTTGAAATTTAGACTTTTTCAATACATTCGGACATTGGCGCATTTGCTAAAGATGCGTAATGTCACTGCAGTAGGCCTTACATACGACGTTACAGATGTTACGAGGAGTGACATCGGCCGCCACAGGAGAGTTTACGATTAATCTCTTTCGATGGCACTACGTACATGATTAAAGACACCACAGGACGCAAGGTTTTATAGAAATACGTTATCCGCACAAATATGTTCGAATAAAATATTTCATCACGTTTGTCTAAATATTACAAATAAAGTGGGTCAGTAGCAACTAAAAAGCCGACAATGGTTTCAAAAATAATATCATTTTACTACCACAGGAAACTGTACTAAACAAACGTTATTAGTAATATCAAAATGATGACATACGATTATCAGCGCAGACTTTGAAGATGTAACAAGTCAATAGCGTAGAGGCATAGTTTCGAAAGTTTGAGATTTATCAAAAGAATATACGGTAATCAGTTCACAATCACAATTGTCGATAAATAAAGACTGCCATTCATCACCCGACGGGATGAGTGTCATCGACTGGTCAGATAGCTGGTCACGCATTATAGGCCATCATTCGAGTTTCCAAAGAAATGACCAGGCATAGCGTAAGAAGTCGCACGACAGGACTCGAGTATAAATTATCGCGCATAGCCAAGTTGTTACTTATCGTATGCTCCATTTGCCGACATTAGCATATGCCCATATGCAACATTTCGCTGTTTCATAAAATGAAGGTATAGTCATTAATATGACGCTTAGTTTACATCGGCAATCATACCATTGCCCCGGCCATACTATTTGGACAGCATTAGCGTTCGTGGCAAAGAAAACATGCATGGCGACATTGGACTGGTTATTATCAATAAAGAAGAGTGTAGTGTCAAAAGTAAAGTCAACAAAAGTAACGGTATCGTTGGCCGCTGAGCTTGACTGACAGACATACGAGGGTGTTAGAATGTCCATTAAAATATGTGACATCAACGGAACATAATATATAAATGCACATTGCCGTTGGCATAATTGCCGATTTCCACGAGTTAAAGTTATTCAATATGAACTTCACGCGACCATACTGTCGATGCATACACTGTTATTACCACGAAGAATCATGCGACTCTCTACATCTCATCGATGATAACATGTACAATATTTCTCGAATAAGTCTTTTAAATAATATGAAAAACTACATGCAGATGCAAAGATGTTTAATGATGATACGATTTCATCTTCAGCGAGAAGATGTCGTGCCAGAATACTTGTCATAACCGTTTAATAATATCAATTTAGTCACGATAATACAACATTTGTTAAGTTCAAACTTTCCACAGCCAGACTGCTTGTAGAATGCCGTGACATAAACAGTCCAGTAGAAGGTACATTCCCACACACTGTTAATTCGCTGGTCGAATTATGATTTGACCATTTTAAAACCTTTCAAGGAGTATCAGCCCACTCATTATCATGATGAAGTTGCGAATTATATGCTTTCTCTCGATAGTATAGATAGCTTAGCTAATGTATAACATAATTCACCATAGCAGTTTCACCGATTTAAAACGATAAACTATTGATTACGCACTGTATAAGTTGCCGGGAAAAGGAGCAGACCCCAATATGTAGATTATAGAGGGCTTCTTCTGCCACGCCTCGCATCTATATGTTCATTCTCATGGAAAAGATGGTCGCAGTGCCATTTGTTGTAAGAGTTAAAACAGAGTCCTGAAAGTACCTCATAACTACGCAGCTCCCTCAATATTGCTCTCACCTCCTTCATCTCACAACCCACAAAAATGGAAATAAAAGCCCTCATTGGGCATAAACGCCTGGGTTGCCTTCTCGCCTGCTCAGACCCTCGCCAGCCGGCGACCCCTACGTGTCGCATGTCTGACTGGTCACACCTCGTGTTTCATAGAGACCCTGCAGCTCCTTCAGCCTCATTCACAACGAGCTTGGCGGTGAAGCCGAGTACTGCGGACAATTGCGTTATTAAATGTTTCATCAGAATGAAACCGCCGAAATATTGCATCATTGACTTTGGTCATCAATATCGTTCCCGGCCAACGATGGGTATAGTCGACATACATGCCGCAGGGTAATCCATAGTCATGCCACTGGGTTGACTTTTTGTCATTCATGAGACGATACGCTTTCGTGAAATAGAAGGACAGCGTTGGCGATTATTGCGGAAAGCACCAGCTTCATGCGTCGATGAAGATGCAAACTGAGCATTGAAACGTCACGGGGTCTCATCGAGCAGACAGCATCTATAAGTCGGTCGCACATGTCTCCACACGCGATACAGACAAATTCGACTGCCTGCGAGTCGAGAGAGTCAGGAAACAACGAAGCGATAACTCGCCAGCGATGCCAGTGTTACAGACATGCAAAATTCAGACAGTACTCCACGATGTCCGGTACATTTCATTTCAACGAATGATTATGATGTCCGTTCAGAAAGATGCGAGGCGCTTAGAAAATTCCCCATATACGAGGAAACGTTAATCTTCTTACGTCATGGTTTCGTCGATGCAATGCGGTACAGCCACATCTGGCAAGGTCGCGTCTCCACTGGCTTGGTCATTTCAAGAGATGAATAACAAATCCAGTTACGTTGATAAACTACTGATGCGACAGGGTCGTTCAGATGGACGACAACAATAATTTTGTTAATTAAACGTAAGGTTATCGCGGAAATCAGATACGCGGCTAAAATCACACAATTATGCTATTGTCGAGATTTACTGCAAGGTCACTGTTATGAAACCATACTACGGCTTGATATACCACAAGTACATGGACGATGACATGACGACAATAAAATCATTATAGTTACGCGGTCGATGTCAATAATATACAATTTCAAAGCGGTTGGTTATGGAGCATCAATGCCGATAATAACAGCGTCATTCAGTAGCAGTTGCCAAAGAGAAAAATAAAATCTATGGAGAAGCCGATTTTATTTCACAGTCATCCAGAGACAGACGAGCCTGGGTCGATGCGCATGCTGCCTATATCTCGAAATCATTCGGGTATTGTCACGCCCAGATGATTTTTGCAAAAGATATCGCCGGTCATAACTGTGCTATAGAGCAGACATTAATAGAATGAAGAATGCCTGGCTATTGGTCAGGCGTTCCATGCTTGATATATGCTTAAACGAAAGAAACGCCACAGACTGAGGATACGCAAGCATTACATCAAATGGTACAGTAATCCACCCAAAGTCCATCACAGAAAACCGTACACGAATATTACAGACGATATTCCTGATTTTCGAAATTTCACCATTCATTGCCATCACGCGGAGCGAGCTGGTCGTCTCAGCCAGCTTCATGATACAGTTGAACGATGATAAAAGATATCGGCATCATAAACACGGGAGTTTGTTTATATGCAAAATCCGCATAAACATTAAAAAATATATTGTACCTGTTTCTTTCTTTTCGTACATCTTCTACTGCGAAAAGTAAATGATGCAAAGGAGATGGGCGATGAGATTGAATAAACCGCATTGCGTTATACCTAACGTAGCAGGACGATGTCACGCTTTCAATGTTTGTCAACATAAGGGTTTACGTTTCGATGTGGACAAAGACCATGTTTAAGCGTTGTTTATTCGAATAAACAAAATCCTCCATTTACCGCCATTGTTTAAATGCCACGGTTTGGTCCATTCGTACAGCATTGGCACAAATACAACAGCAACCCGATAGCGATGCAGGTGCCACTACTCGAGCCCATTGTCATCAATGAGCTGTCAAAGGTTTCTTTCGATACACGACGTTCTTTAAAACATCGATCTGTTGCTGCAAGTTATTTTACCGTCGGGAAAGCAAATTGATTCATTTAATTGGCTCTTTCATTGGTCATTGCAAAACACATGTTCACGTTCTCGTGCTCATTGTTGTGTTACATAGAAATTGCCGCGTGAGCAAACAGATAACGTAAGTCAGCACTACCTTCAACAGCATGTCAAACATGTTCTTCGCATGCTCTTTTGCCCGAGTGACAGCGGCCACCGTGCCAATTGTCATGTAAGTTACAGCTCTCTGCTCGGCATCAAAGGAAGAAGGACACCACTTTGCCCATCATAAAACATATCGCGATACACTCAATATACCACGGCATTCGCCAGAGAAGACGGCCACAATACGCTCAAAATTAAAATATACATATGAATATCAACCTTCCGATGATCGCTCGACTTTCTGAACAGAAACATCGTTATTAAGCAAGTGATGTTGTGACGACACACACAGACTGTATTACCCATGATTAAATGTTACAAGATGCTCAGAGTTCTCGTCGATTACTGCAGCCGAAGTCATCGTTTACAGTGATTTATCATTTTATAAAATTGCTTAAATGAGATTATAAAAGTGCATTATTGAAAATATCATTCTCGACAAGATAGATTCTGCCTGTCGCGACATAAGTCGCTCAAAGTTATATTACCCCGTATGCTTGCTTCAAAGTTGTCTTGAAATATGGGGACAGAGTCAATATTACTTCCTTCTCATACGACGATCCGGTCAGTGTCATCTGTCGATGGAAAGCGTAAACCCAAAATGACAAAGTTACCAATAAATTGGGAAAGTTCACATCTGGCCGTATCATGCGCACACACCAAAGAAAGGCAGCTGTATTTGAAAGAAATCATTGCGTCATTATTCATGACAGAAGAAAATGAATTGTTGGAGAAGTAAAAGACAATATTACAATTATGAATATACTGTATATTGCTCTCTCATGCATATCAAAGACAAACATCCGATGGAAAGTGGAAGACAATTTAACAAACGTTACGGGTTTCAGTCTGGGGAACATGCTTCCAGACAGCAGGTTAAATGGTCGCCAAAGCCGTGTAACACTGCAATAATACCGTTGTCGCCGTATGGAAGACGAGTCAAACATCTTCACAATAAGTCAGGCCTCCGTGAGCTCTCCCCGGATATTGCCCGCATGCTTTCAACAAAATGTGAAGGATTACGGAGAACAGCCCGGCTCGTGTTCGGAATATCAAACTTCTTCGGCAGAATCACAGAAATGTTTCCACCATACGATGCGGAAATTAACCAACGCCGACGACACTGGCGCTCGCGACCACAATGCCGGAAATTATTGCTTTGCCAAAAGATCTCAAGAGCCAAGATGGATATTCTCAGGATTTTCACAGGACCCCGGAATTTCGAGTCAAAGGCAACAAAGATTCTTCTCCAAATAATCTAGTAAAGACAGAACATGTCAGCAGGAGACTGCCTATCATTCATACTATGCGGGTCAATAATACTACATTTGAAAACCGATACAATCTCTTACTTTTCTATGTCGAGGTTATTATGCTCGGTATTAGCGATGACCCCGCTATAGTAAGTTTTTCACCATAACAATACAATAATAATTTCGTAAAAGCAGAAAATATATTCTGGTTCTGTACGGCAGCGAAGCAGAATATGTTAACAATAAAGGGCCGGCGAAAGAATATACGTCTCGGTGATGGATGTTCGCAATAGCCTCGCAAATCATCTCATTCTTCCTTTTCGCATGGAAGAAGAAAACAAACATCATGAAGGGGAACGATGTTACATCAAAAGTCACAGTCACTTTATCATGATAAGATTTGCCGACCGCATAGAGTTCTGAGTCAGTTCAGATCTCTTTATAGTGGGAACAGAAGATGATCAACATCAAATGAGATTCTCATCATCTGATACCATCATCGGTGAGGTTCAGCGAACGGATCACCAATGTCCAGACACCATTCAGAATCCGCCGATTGAAACACAGTGAATGGCACAGTCTGGGTACCTCGCTTTCTCTCGCGATCACCAGAAAGAAGTTTCGCTGTCAAAGACTGTGAGGTGCTCCACGACACGACAAGACAGCGACACCATTCAGGTTCAACATCATACAGTGAAAGATCGGTTCAACATCGACAAAATGCATCAGAATCTCAGAAGTTTGAAGTAGACAGTAATCACCTGACCTGAGTCGGCGATGGATTCTACGTCGACGATCGGGCATTCGAAGGCAACCGACTCGTCGATCAACAAAGTGTGTTCGACGATAATTTGTTCATTCGAAATTGGATAATCGAACGATCAGGCACACCACGAATTGCAAAATTATATTTGCTTCAAACAAAATGTTCTAATTGCCATCTTCACAAAAGTTATTTCGTTACTTCTTTTCTGTTCAATAGAAGATTGTCAATTATTCCTGTTCCCCTATTCTTTGCTTCCTCGGCCAAAAATATTAAACCTTCCTTTTCGATGGTTCATAAAAAGCTTCAAAATAGTTTTATTCTTCTTTGATAACTGATAGGGTTTTATTGTGTATGATTTTGATCCATAGGGAAGTTTGTAACATTTTTAAGTCCAGGGAAGTTCTTGACAGTTTTATGTCTCTGATGAACCATAGGGTCTTTATTTTATCTTCATAGGCCTATGTGACAAAATATTTATTCTGAGTCCTCATTGTAACGTTTATTTGAATTTCTGATTGCCGTGAGGTCGTCTATTGATAAACCATTGTCATCCCAGGGACTGACCATAATTTATCCTGACCCTTATGTTTTCCCGTCATTTGTTCATATCAACATGTCTATTCATGGCCTGACAAACAAGATTAAAGTGAGACAGTCATAGCATAATACTTCGCCTTTATCACATCACCGCACAAGGGACGAATGTCGAATGCAGCATGGCGGCAGATCTGACTGCAGTTTCGTTCATGCCAGCGTACTGTACGTTCATCATGGTAGTCTCGCCGCGGGGGTGGGTGACGTCGACGCCGGGCAATGGCTTCGGGTTGTGCAGACCAGCGTCGCAGTGGCCGCCGCCCTGTTTTCAAAGTTGACTGCCCTTTCAGGGCTGACCGAGCCTCCAGTTCATTGTTTTCCCGCGAGTTTGTAAGTCGTCGGAGGAGAAAACAGCAAAGCCTGTAGTAATCCGTCCTGCAGGAGTCTCGGGTCGCGCCCACGCGCCCTCATTCTTCCTCGGTAGTCGGGGATGTCGGCAGCAAGCCACGTAGTTCGAGCCGCATTACGAACTATGAAAGGTGTTGGCCAGTTTTGCGCGAATTTGCCAGGTCTCGCCCGCTCGCCTGGGCGCATTTTTGATGATGGCCATGTTTCCTCGCCCCGTCACCACTTCATATTCAAAAAAAAAATAAAATTAATTTTTAGAGTTTTTCTGTAGGGGATTCGGAATAAGCAGTTCAGGAGATTTATGATGAGCCAGAGGTACCACAGGGGAGGTGGCAGAAGTTAGTAAGGGCGGGCAGAGATGTGAGCTCGAGTTGCCGTGTTCTGGAAAGTGCAGTAACCCGTTGCGAAAGACCAAATATTCTCATTATTTTCTTTCAATGGAACATAAAAATATTATCTCTTCGATAAAGCGTACCGGACATCTGCTGCTTCAAACAAAATGCCTTTCATCCACATCACAAAGTTGCCTTCGTCATTTCTTTCTCTTTTTCTCTGTTCACAGGTGATGAATATTTCATCAATTATTCCTTGCCCTTATTCCTGAGCCTCTCAATAAAACATAAACCTCTTCGTGAGTCCAGGTTTTTATAACCAGCCTTATTAAAGATGTTCAGGGAAGTCTATGTTTTATGTTGTCTCATTAGGTTCTTGATATCCATAGGTTTTATTGTCCATAGGGTTTTGACAGGGAAGATGGAGGAACATGAATTCTGATTTATGTCAATTAAATATTTAACAGCGATTTAACTGACATGGTTCACCATCGATCCTTTTGTAATCGACACTATATCCTTTGTAGCATCAACTTCAATCACTTTACTTTGGCTGTTTTCTCATAACATACCCACCATATGCATCTAACATTTCAGTGCGAGACAGTCGTTATCATAGCGTATTGGCTGGCCCTTCGTACTGCCCTTGATACCGCATTGCCAAATTTAACAACAAAATGCAAAGTTTGCTCGTACAGTCAAACACGGTGTGATGTAGCATGGCGCCCGACGATGTAGTTCCGCCGTGGAGACCAATCCGATGCCGATAGATGCCGGGCATAAGCCGCACGCTTCAGCACCAGCGTCGCAGGCCGCGCTTTCTCCTCGCTCAGGTGTTCCGTTCTCGCCTCATTTATTTCGACAGCGTAGTCAGTTGTCGCAAAGTCACCTGTAGATGACCGTCGTCCCGCAGGGAGAGCAAAACAATGTCCTCGAGTCACGCGCCCTGCCTCGGGAAGCGACAGCAATGGGGGATGTCGGCGGGCAAGCCATCTGGAGTCGCAATACTTGTGAGGGGGACACAGGCTAGCAGCAGGGGGCATCAGCTTGCAGCCCAGCCTGCAGGGGCGGCCCTCGCCCTCGCCTCGATCTCGACCACCATTTGTGACCTGAGCATGAACTTCCCGGATGATGCACATGTTATACCTCCCCGCCCCGCTCAAATTACTATATTCCAAAAAAAAAAAATAAAAATTTGGTCTAGAAAGTCAGCAAGACGGGACGCAGCTCAGAGGCATGGGAGGCAACGCGAGCCAACCGGTGGATGGCTCAGTAGGGCGGCGGGCGCGAGCCCCTCGAAGCTTCAGTTGGAAGAGTGGCAACTCCGTTGCGAAAAGTTGACTGCCTTTCTTCTTCTCATCGGAGTCATAAAAATATTTTTGTTCTTCTTTTTCGATGTTCAAAAATGGACATCGTTTGCAAAAGTGGGGTCAGCCTAACAGATGATCATGGACACTACTCCTGTTCTCACTTTCACTAGATTGGGAAGTTATTTTCTTATTGAACTAGGAAAGTTTATTCTGATCCTGATAGAAAGTGGCCATTTTTATTCTGATGAACTGATAGGAAGCTTATTAAATATTTTTTAGTTAGTTTTTGATTATGTGGATGATAGATAGGGTTTTATTAAATATTTTTTTCTTCTTTCTGATCCAGGTTGTATTTTGGATGAACTGTATTGAACCCTCTCTGATGGTGTCACCAAATATTTTGCAGATGAATTTAGGACAAGTTATGCTGTTTGGGTCCATAGTCAGTTTATGATTGTTATTATCCTTAGACTGATCTGAACTCAGGGATCCTTGCCCATCAACCACATGTATTCATTTCGCCAAGATATGAGATAGTCGTCGAGGCAGAGCGGTCTCTGCCTGCCCTTGTTGAATACCGCATTGGGCGCATAGTCAAGCATGGTGACAGATGCCGGTGGAGTGAAGCTCATTTTGCGTGGTTCTCGTTGCGGGAGATACCACGCGGCCGGGTGCCGCACTGTTTTCTTGCTCCGCAGGTGGTTCGACCAGCGTCGGGCGGCCGCCGTCCTCAGCTGGTCGCTGATCTCGCCCTTTCAGGGCGCCGTCTCCGGGGTAGATCTCGGAGAGTTCGCTGTAGTTGCCGCGTAGTTTCATCTGTATCCTCGTCGTCCGCATGAGTCATGCTCACTCGAAGTTGCACGCGGGTGCTGCCGGCCTTTGAGCCGCACATGACCGAGGGGACATAGTCACCTCGTCGGCCTGCGCTGGGGCGGCTCGCTCTTTCGATTTGATGAACTTCTGATGATGGCCATACCTTCGGTCTGCTCACCCGCCTATATTCCAAAAAAAGGGCGCTTGGGATTCAGGTCACGTCTGACGATGATTTAGAACAACGGGAGAGCAGGTTATCTTCTCGAGTTTTCTCAGTTGTTCGGCGGTAAATCTGTTGGTGAAAAGACTTTGTGTCATGCTTTTGATTATCTTCATTTTTCTTTCTTCATCGGAGTCGCCTTTTATTCTTCTCGCCGATTTGTCTAAAAGCAGTACCGACGATCTAGTTCTCGGTTTCATAAAATTCAAGACCATCTCACGTCTGCGCTTCTTTCTTTTTCAATCCGTGATTATGATCATTCATTCCTAGTTCCTGTTCTATTTTCTTCGTCAAATATTAAAGTTTTTATAAAAATATTTTTATTCTTCCTGATGAAACCTGCCCTTTTGATGACTGATAGTGCCTTACAAAATATTTATTCTTCTTTTTCTTTCTTGATGAGTCTTCTGATAGGGTTTTATAGGGTTTTATTCTGATCTCATAGGGGTTTATTAAAATATATTCTTTTCTGATGAGCTTATGTAACATATGGATTTCTTTCATTAGTAAAGTTTTATTAAAATATTTTATTCTTCTTTTGATGTTCATAAAGTCTTTATTTCTTGATGTCTCATAAAATATTAGTCCTTTCTGATGGAGAAACACTAAAGTTTTATTTTATTCTGCCACTGAGTCATAAATATTTTATTTTTGATCTCATTGTGGGCTAGTCCCTTCTCAAACATTGGTAATATTTATTTCATTCTTTTGACATTTTCTTTTCTTTCCATAGAGTTAAATATTTCTTTTGTTCTCTTGATGGTCTCATTGTTAGTCCCTTGATATTTAAGTTTATCTTCTGATCCCCACTGTAATATTTTATTTCTTCTGATGCATCAACCATTTCAACATCCAACAGCTCCACGATCAACAGCCTCGGCTATCTATATCACTGCACGTCAATGCCGGTTAAAACATATAACGTGATCGCATATGGCGGCGTAAATGATTCAGTAAACCACGGGGAGACGCTCACGGCCTTTTCTTGGCGCTGTGTCAATGTTTACACAGCATCAAATTTTGTCCGGCGATACTAAAACGAGCCGACTCGCGCCATGGAAAACAGCCCAGACTGGTCCGTCGATGGTTCATGCAAATATTGAAAAGTTTCTTCTTCGATATTCCCCGTCAGAGGCGAACCTTCTTTTCGATGTCGTAAAGCCTATTTTTCGATGGTCTCGCCAGTCCTTTTTCGGGATAAGTCTCGCAAGATGGTTCACGTTGAGGCCAACCTCTCGGATAAAAGCATTCTTCGATGTGAGCCCTCACGTAAATATTGAAATTTTCGATGGAGTCGTAAAGCTTTATCTTCTCGAGGAAGTTCTCTCTTCAACCTTTCGATGGAGTCGTTTTATTCTTTTCGATGGTTAAATTCTGATGTCGTGCTGATGTCTCGTCAAAAATATTAAACTCTCGATGGAGTCGTAAAGTTTTATCTCTTTCTTTCGATAAAAATATACATCCCTTTCGAGTGTTTTATCTCTTCTTTCGATGGTCTCGGTCAAAATATAACTCTTTCGAGTCGTAATATTCTCTCTCGATGGTCTCATATTAACCTTCTTTTCGATGTCGTGCTTATCTTCTTTCTTCATTTGTCCCCTCACATATCCTGTTGATAGTCAGTTTTATTTTTTGATATTAAATAGTCCTGAGTCAGCTTTATCCACAGCTGAACTTTGATGATTGTCTTTGTCTCACAGTTAGTCCTGAGTCAGGACCATTAAATCCTTTCTTATTAGTCCTTGATGAAAGTCAGATCTTTTCTGATTGAATCTCATTGATCAAAATGACCTCACACAGCATCCCTTTCGAAACCATCAGGGTCGTGTTTTATTTTCTCCTGATTGTCCAAAATAGTTAGTGATCTTGATGTAGTCAGCTTATCTTTCTCATCAGGAAGACAGTCCTCTTTTCTGATGTTCCACAGAACAGTCCTGATGTGGGCTTTTTCTGCCTGATCCCTGATGGAGCTTATTCTCAAAGTCTCAACCTCTCATCAGCCAACTCTTGTCAGCCTATGCCCTTTTCCTCTCGATGTCTCACCTCTTTTCTGATGGTCGTTTTATCTTCTCTCACTGATGTCAAGCCTGATCTTCATTAAAATAACCTTCTTTTTCGATGGAGTCGTGCCGTCTTCTTTCTTTCGATGGTTCAATAAAATAGATGGAAATGTGCCATCTTCTTTCGATGAAACTTCATTGTACATAGTCCTTTTTCGATCTTATATGATGATATTTTTTTAGACACTAACAAATATTTAACATAATATTAATATACTAAAATTTATGTATTATTAATTTATTTAACTAAAGTTATAAATTATATACATACTCTTTATAATTAATTTGACTTTACTATTTATTTAGTGTCTAGAAAAAATGTGCGGACCAACGACCGTAGGAAACTCTAGAGGGTAAGAAAAATCAATCGCTTTATAAGACCATCAGAAAGAGGTTTTAATATTTTGTGAGACCATCGAAGGAGAAAGAGATAAAACTTTTTACGACTCCACTCAGAAAGAGGTTTAATATTTTGTGAGACCATCGAAGGAGAAAGAGATAAAACTTTTTACGACTCCATCAGAAAGAGGTTTATTGTTTTCAGAGACCATCGAAGGAGAAAGAGATAAAACACGACTCCACTGGGGAAAGAGGTTTCTAATATTTGTGAGACCATCGAAGGAGCAAAGAGATAAACCTTTTTTATGACTCCATCAGAAAGAGGTTTAGCATTTCAGAGACCACTGAAGGAGAAAGAGGATAAAACTTACGACTCCATCAGAAAGAGGTTTTAATATTTTTAGAGACTACTCTAGCTGAGAAAAGAGATAAAACTTTTTGACTCCACCCACCAGAAAGAGGTTACTATTTTGTGAGACTACTCGAAGGAGACTCCATCAGAAAAGAGGTTGCAATTATTTTGAGGAAAGGAGAAAGATAAAACTTTTTTACGACTCCATCAGAAAGAGGCTTGAATATTTTTGTGGAGAGTGAGAAAGAGATAAACTTTTTACGACTCCATCAGAGAGGCTTCTAATATTTTTGTGAGACCATCGAAGGAGAGAAAGATAAAACTTTTTACGAATTCCATCAGAAAGAGGCTTAATTATTTTTGTGAGACCATCTAGAAGGAGAAAGAGATAAACCTTACACTCCATCAGAAAGAGGAATAATATTTTCAGGAGACCATCATCGAAGAGAGAAAGAGATAAAACTTTTAATGACTCCACTCAGAAAGAGGTTTAATATTTTAGAGACCATCGAAGAGAGAAAGAGATAAAACTTTTACGACTCCATCAGAAAGAGGTTTGGCATTTTGTGAGACCATCGAAGAGAAAGAAAGAGATAAACTTTTTACGACTCCATCAGAAAGAGGTTTATTATTTTAGAGACTATCAAAGAGAGAAAGAGATAAACCTTTTACGACTCCATCAGAAAGAGGTTTAATATTTTTGTGAGAGACCATCGAAGAGAGAAAGAGATAAAACTTTTTACGACTCCATCAGAAAGAGGGTTTAATATTTCAGGAGACCATCGAAGAGAGAAAGAGATAAAACTTTTTACGACTCCATCAGAAAGAGGTTTAATATTTTTGTGAGACCATCGAAGAGAGAAAGAGATAAAACTTTTTTACTGACTCCATCAGAAAGAGGTCAATATTTTTGTGAGACCATCGAAGAGAGAAAAGAGATAAAACTTTTTTTACGACTCCCATCAGAAAGAGGTTTAATATTTTTGTGAGACCATCGAAGAGAGAAAGAGATAAAACTTTTACGACTCCATCAGAAAGAGGTTTAATATTTTGTGAGACCATCGAAGAGAGAAAGAGATAAAACTTTTACGACTCCATCAGAAAGAGGTTTAATATTTCTGTGGAGACCATCGAAGAGAGAAAGAGATAAACTTTTTTACGACTCCACCCATCAGAAAGAGGCGTTTAATATTTTGTGAGACCATCGAAGAGAGAAAGAGATAAAACTTTTTACGACTCCATCAGAAAGAGGTTTAATATTTTGTGAGACCATCGAAGAGAGAAAGAGATAAACTTTTTACGACTCCATCAGAAAGAGGTTTAATATTTTGTGAGACCATCGAAGAGAGAAAGAGATAAACTTTTTACGACTCCATCAGAAAGAGGTTTAATATTTTGTGAAAGACCATCGAAGAGAGAAAGAGATAAAACTTTTTACGACCCATCAGAAAGAGGTTTAATACTTTTGTGAGACCATCGAAGAGAGAGAAAGAGATAAACTTTTACGACTCATCAGAAAGAGGCTTAATATTTTTGTGGGAGACCATCGAAGAGAGAAAGAGATAAAACTTTTACGACTCCATCAGAAAGAGGTTTAATATTGCAGGGAGACCATCGAAGAGAGAAAGAGATAAAACTTTTTACGACTCCATCAGAAAGAGGTTTAATATTTTTGTGAGACCATCGAAGAGAGAAGAGATAAACTTTTAATGACTCCATCAGAAAGAGGTTTAATATTTTGTGAATTATCGAAGAGAGAAAGAGATAAAACTTTTTACGACTCCATCAGAAAGAGGTTTAATATTTTGTGAGACCATCGGAAGAGAGAAAGAGATAAACTTTTTACGACTCCATCAGAAAGAGGTTTAATATTTTGTGAGACCATCGAAGAGAGAAAGATAAAACTTTTTACGACTCCATCAGAAAGAGGTTTAATATTTCGTGAGACCATCGAAGAGAGAAAGAGAAAGAGATAGTTAGTTCAGATATTTTCTTGTACAAAAGTCAATAATGTTTTAAAAATATAAGGGACAAGAATTTGTCTGTATAAAAACTTGTGTGAAATTTTGTACCAAAAGAAAAATGTGAGGTATCCCTCATATGGATTTTACAGATCATTTATATAACAAAAATATTATACGATCTACGTTTTATTATATGATTTTAACGTAAATTATAAACATTATTTTGGCGACATACAATTAATTGTAAACCTAGATAAGCATAGGGGATGTTGATAAGCTCGACGAGTATATGCTGTTGGACGTTGTTTAAGAAATAGTTGATGCATCAGAAAGAGAATAAAAATATTTTAGTGAGACCAATTGCGAAGAGAGAGAAAGAGAATAAAATATTTTATGACTCCATTGAAGAGAGAAAGAGAAAATGAGAATGAGAATAAAAATATTTTGAAGGACACCACTGTGAAAGAGGTTTAATATTTTTGTGAGCAGACCATCAGAAGAGAAAGAGAATAAAAAATATTTATGACTCCAATAGTTAGAGAAAAGAGAAAATGAGAATGAGAATAAAAATATTTTAGTGACATTCATCAGAAAGAGGTTTAATATTTTTATGAGACCACTAAAGAGAGAAAGAGAATAAAAATATTTTGTAAACTTTTATGAGACCATCAAAGAGAGAAAGAGAATAAAAATATTTTGTAAAACTTTTTTGTGAGACCATCGAAGAGAGAAAGAGAATAAAAAATATTTTTGTAAACTTTTTATGAGGACCATCAAAGAGAGAAAGAGAATAAAATATTTTGTAAAACTTTTATGAGACCATCAAAGAGAGAAAGAGAATAAAATATTTTGTAAACTTTTTTTATGAGACCATCAAAGAGAGAAAGAGAATAAAAATATTTTGTAAACTTTTTATGAGACCATCAAAGAGAGAAAGAGAATAAAAATATTTTGTAAACTTCTTTTTTATGAAAGAGAATAAAATATTTTTGTAAAACTTTTTATGAGACCATCAGAAAGAGGTTTAATATTTTTGTGATAATTCAGGCGAAATAGAATAGGAATAGGGACAGTGTCATAAATCATCACACTATTGCTGAGACAGAAATATAGTGTCCGGTACACTTTTAGACCATCGAAGAGAGAAAGAGAATAAAATGGCGACTCCATTGAAGAGAGTAGAGAAAATGAGAATGAGAATAAAAATATTTGGCCTTTTCGCAACGGGTTTGCCGCCAGAACACAGCTGAAGCTTCGAGGCTCGCATCTCTCTTCACGCGCCCGCCCTACCTGAGGCCGCCATCCACGCCGGCTGAGCTGCTGCTCCTGCCGCCTCCCGCCTGTGGTGCCTCCTGAACTGCGCGTCCGCCGTCTAGCTCCCGGGCAACGGTACCGGACTCGCAGAAGCTTCTAAAATTGAAATTTTTATTTTTTTTTTTTGGAATATAAATGCAGGAGCAAGGGCGAGAGGATAACATGGCCATCATCAAGGAGTTCAGCGCGCTTCAAGGTCCACATGGAGGGGCTCCGTGAACGGCCAATGAGTTCGAGATCGAGGGCGAGAGGGCGAGGGCCGCCCCTACGAGGGGCACCCAGACCCCTGCCAAGCTGAAGGTGACCAAGCGGGGTGGCCCCTGCTCTTCGCCTGGGGACATCCTGTCCCTCAGTTCATGTACGCTCCAAGGCTCACGTGAAGCACCCCAAGGCCGACATCTCCGGACTACTTGAAGCTGTCCTTCCCCGAGGGCTTCAAGTGGGAGCGCGTGATGAACTTCGAGGACGGCGGCGTGAGACCGTGACCCAGGACTCCTCCTGCAGGATTTTAAGGAGTTCATCTGGCTAAGGTGAAGCTGCGCGGCACCAACTTCCCCTCCCGACGGCCCCGTAATGCAGAAGAAGACCATGGGCTGGGAGGCCTCCTCCGGAGCGGATGTACTTCGAGGACGGCGCCCCTGAAGGGCGAGATCAAGCAGAGGCTGAAGCCGGAAGGATATGGCCACTACGACGCTGGAGAGGTCAAGACCACCTACAAGGCTGGGCAGTCTCCGTGCAGCTGCCCGGCGCTCAACGTCAACATCAAGTTGGACATCACCTCCCCACAACGAGGACTGGCTACCATCGTGGAACAGTACAACGCGCCGAGGGCCGCCACCCACCGGCGGCATGGGATGAGCTGTGCCAGTCTAGATTCGCCACCATGTTTGCATTTTGTTATTCGTTGCCCAATGCGGGTGATGTAATAAAGGCAGAGTATACGAGAAGGATTATGCTCATATATTTATCTTTTTTGACTATCCTCACTTTGAAGCTATCTTACAGAGAGTGCGTTAAGATGCATGGATAGATATGTTGAATATAGATAAACTGGTAGGAAAACCCGTAAAAGTTAAAGTGATTAGAGTTGATTATACAAAAGGATATATAGATGTCAATTACAAAGGATGTAGACATCAATAATAGCGGCCGCCCAACTCTGTTTATTGCAGCTTATAATGGTTGGGCAACAAAAGCAATAGCATCACAAATTACTGAATAAAGTGATTTTTTCACTGGGTAACATTTTTGAGTGATTAATCAGATCAGAAAGAGGGTTTTAATATTTCAGGGAGATCCATCGAATAGAGAAAGAGAATAAAAAATATTTTTGTAAACTTTTTATATATCCACTAATTATGAGACCATCGAAGAGAGAAAGAGAATTGTAACATTTTAGAACTTTTTATGAGACTGACCAAAGAGAAAGAGAATAAAATATTTTTAGAACTTTTTATGAGATCCCGACCGAAGAGAGAAAGAGAATAAAAATATTTCCCAACTTTTTTATGTGAGACCATCAGAAAGAGTTTTAAATATTTTGTGATACTCCTGGAAAGGAAATAGGAATAGGAATAGGAATAGTGTCATAATCGTATCACACTATTGAGACAGAAAAAGAAGAAGTAACGAGAGGTAACTTTTGTGAATGTAGTTAAGAACATTTTTGTTTTGCAAACCGGAACATAGTGTCCGGTACACTTTTAGACCATCGAAGAGAGAAAGAGAATAAAAATATTTTATGACTCATTGAAGAGAGAGAGAAATGAGAATGAGAATAAAATATTTTAGTCTTTTTTCGCAACGGGTTTGCTGCCAGAACACAGCTGAAGCTTCGAGGCTCGCATCCTCCTTCACGCGCGCCCGCCGCCCTACCCGAGGCCGCCATCCACGCCGGTTGAGTCGCGTTCTGCCGCCTCCCGCCTGTGGTGCTCCTGAACTGTGTCCGCCGTCTAGCCTGGGCAACGGTACGATCCCTGCGCAGAAGCTTCAAAAATTGAAATTTTTTATTTTTTTTTTGGAATATAAATGGTGAGCAAGGGCGAGGAGGATAACATGGCCATCATCAAGGAGTTCATGCGCTTCAAGGTCCACATGGAGGGCTCCGAACGGCCACGAGTTCGAGATCGGAGGGCGAGGGCGAGGGCCGCCCCTACGGAGGGGCACCCAGACGGGCCAAGCTGAAGGTGACTGGGCGGCGGCCCCCTGCTCCTTCGCTCCGGGGCCATCCTGTCCCCTCAGTTCATGCAGTACGGCTCCAAGGCTCACGTGACCCCGCCGACATCCCTCCTCTCGACTACTTGAAGCCGTCCTTCCCCGAGGGCTTCACAAGTGGGAGGCAAAATGAACTTCCTGAGGACGGCGGGCGTGGTGACCGTGACCCAGGACTCCTCCCCTGCAGGACGGCGAGTTCATTCATAAGGTGAAAGCTGCGCGGCACCAACTTCCCTCCCGACGGCCCCGTAATGCAGAAGAAGACCATGGGCTGGGAGGCCTCCTCCGAGCGGATGCACCCCGAGGACGGCGCCCTGAAGGGCGAGAGATCAAGCAGAGGCTGAGCTGAAGGACGGCGGCCAATACGACGCTGAGGCCAAGACCACCTACAAGGCCATAGCCCGCAGCTGCCCGGCGCCTACAACGTCAACATCAAGTTGACATCACCTCCCACAACGATGACCACAGCCTTCACGTGGAACAGTACGAACGCGCCGAGGGCCGCCGCCACAATCTACCCGGCGGCATGGACGAGCTGTACAAGCTCCATCTGCCACCATGCTTGCATTTTGTTACCTGTTGCCCAATGCGGGTGATGCAATAAGGCAGAGTATACGAGAAGGATTATGCTCATATATTTTATCTTTTGACTATCCTCACTTTGAAGCTATCTTGGCAGAGAGTGTTAAGATGCAGGCGGATATGTTGAATATAGGGATAAACCAGGAAAACTGTAAAAGTTAAAGTGGATAATTATAATAAAGGATATATATAGATGTCAATTCACAAAGGATGTAGACATCAATAGGCGGCCGCCCAAACTTGTTTATTGCAGCTTATAATATTACAAATAAAGAATAGTGATCACAAATTTCATTGAATAAAGCATTTTTCATCAGCAAATATTTTAGTGACACCATCAGAAAGAGGTTTAATATTTTTAGGAGACCGATCGAAGAGAGAGAAAGAGAGAATAAAAATATTTTGTAAAACTTTTTATGAGATCCATCAAAGAGAGAAAGAGAATAAAATATTTTAGAACTTATGAGACCATCAAGAGAGAAAGAGAATAAGTAAATATTTGTAACCTTTTTAGATGAGACCACTAAAGAGAGAAAAGAGAATTGAAAATATTTTTGTAAACTTTTTTTATGAGACCATCAAAGAGAGAAAGAGAACAAAAATATTTTTGTAAAACTTTATGAGACCATCAAAGAGAGAAAAGAGAATAAAATATTTTTGTAAAACTTTTTTTATGAGACCATCAAAGAGAGAAAGAGAATAAAAATATTTTTGTAAACTATGAGAGACCATCAGAAAGAGGTTTATTATTTTGTGATAGAAAGGAAATAGAATAGAGGGAGTGTCATAATCGTATCACACTATTGACAGAAAAGAAGAAGTAACGAGAGGTAACTTGTGAAGTTACAACATTTTTGTTTGCAAACCGAATATAGTATGTCCGGTAATACACCATCGAAGAGAGAGAAAGAGAATAACAAAAATATCGACGAAGAGAGAAAGAGAAAATGAGAATGAGAATAAAAATATTTTAGTTCTTTTCGCAACGGGTTTCGGCGCGAAACACGCTGAGGGGGCTGCAGAGATTCTCTCTTCACGGCGCCCGCCATCACCTGGAGGCCGGTCGTACTCCTCCCGCCCAGGGTGTTCTGTTTTTCTGAATCAGGCCAGTCCGCCCAGCCGCCCGGGCGGTACCGGATGCCATCATCATGAGTCTGCTGCGTGCGCTTCACATGATCGTGACCCAGGCCACGAGATCGAGGAGAGGGCAGAATTCCTGCGCACCCGACCGCCGTTGAAGGTGACCACCAAGGTGCCTCTGCCTGGGGACATCCTGTCCTCGCCTGGCGTACGGCCCACGGTCACGTGAAGCAATCTCCCGCCGACATCCTCCTTCGAAGGACTGCTCACGTTTCGTCCTTCGAGGGCGAGCGCGCGCGTATGAACTTTCAGGCGTGAAACCTCCCGCGAGTTCACTACAGCCGCCAGCGGCACCAACTCCGCAGAAGGGAGTGCGATACCGTGCGAAGGACCATATGCTGGCGCCTACAACCGTCAATAACATCAAGTTGGACATCGCCTCCCACAACGAGGACTACACCATCGTGGAACAGTAATGAACGCCATTGAGGGCCGCCACTCCACCGGCGGCATGGACGAGCTGTACAAGTCTAGATCTGCCACCATGTTTGTGCATTTTGTTGTCTAAGGTTCACTGTGGCAGATGTGAATAGCAGAGTATACGAGGAAGGATTATGCTCTATATTTATCTTTTTGACTATCCTCACTTTGAAGCTATCTTGGCAGAGAGGCAGATGTATATGGATAGATATGCTGAATATAGGGATAAACTGGTAGGAAAACTGTAAAAGCAGTGATTGAGTTGATTTATAAAAGGATTGGTATAGATAAAGTCACTCATAAAAGGGATGTAGATCAATAATAGGGTGGTCAGCCTAAACTTGTTTATTTGTAGCTTATATTTGTTACAAATAAAGCTAAATAGTGACTGCAATTTCACAAATAAAGTATTTTTTCACTGCAAAAAATATTTTAGTGACAATTCCATCAGAAAGAGGTTTAATGAGAGAAAGAGAATAAAATATTTCTGTAAAACTTTTTATGAGACCATCAAAGAAAGAGAAAGAGAATAAAAATTATTTTGTAAACTTTTTATGAGACCCATCAAAGAGAGAAAAGAGAATAAAAATATTTTTGTAAAACTTTTATGAGACCATCAAAGAGAGAAAGAGAATAAAAATATTTTTGTAAACTTTTATGAGACCATGAGAATAAAAATATTTTGTAAACTTTTATGAGACCATCAGAAAGAGGTTTAATATTTTGTGATACCCTGAAAGGAAATAGGAATAGAAGAGGAATAGTGTCATAATCGTATCACACTATTGAGACAGAAAAAGAAGAAGTAACGAGAGGTAACTTCTTTTTGTGAATGTAGTTAAGAACATTTTATTTTGCAAACCGGAATATAGTGTCCGGCAATAATTTTTAATTGTGGTGTCGAATCGTTCGATTAACCCTACCCATCCAATTTCAGATGAATAGAAGCTATCGATTCAGCATCATCGCTTTGAGTTTTGTTGAATCGATGAGTGAAGTATCATCGGTTGCACAGATGCCGATCATACTTGAATCCATCCTTGACCTCAAGTTCAGATTTTCGATTCCTTGCACATGTTCCGATACGAACGCTAAACTCTAGATTCTTTTATCATCATTATTTTGTATCGACGATCGTTGGAACCGACGATATCTTCGTAACTCACTTCTCTTATATTAGACCGGAGTACTGATGGGTCTTGATGTCGCTGTCTTCTCTTCGCTATGATCTGATGTCGATAGACACCCTCACAGTCTTGATCATAAAGCCAGAGCTTCTTCACGAGTGGATCGCGGAGAGTCCTTACCTGTCCTGTACGCTGGACAATCTAGCATTCACTGTGTTTCCATCGAAGCGGATTCTGAGGATGTGATTTAATCTGAGGACGGTAATCCAAAGTTCATTCAGGACCTCCAATTGATGATGGAGTACAAGTGGTAGGAGGATCACATCTCGACTGGATGTGGAATCACTTCTGATTCCAATCTCGATCTGGATCTGACTCGGACTCTGTAATTTGCTACGGATTAAATCTTATCATTGGTCGGTGTTTGGTCTTGCTTTGGTGGACTTTGATAATAACATCGATTCCTATAAGGATGTTTGTTTTCTTCTTCAAACACGAGGAGGAGGATGAGGATTGCTGAAGACTGGCAGCAATGATGCATGCCAGGACGATATATTGTTTCATAATTCGTTATTGATTGAGTACTGTTCTTTATGATTCTACTTTTACTCATGCACAATTAGAATATATTTTCTAATTTTACGAGAAATTAATTTATTGTATTTATTATTTATGGGTGAAACTTAGCCTATAAAAGCGGTGGTTTGGAATTAGTGATCAGTTTATGTATATCGCAACTACCGGGCATAGTATTATCGACATCGAGAACATTGTACCTACATGATAAAGAGATTGTATCAGTTTCGTAGTCTTGAGTATTGGTATTACTATATAGTATATAGATGTCGATAGATATATAGTCTCCGGAATGCGGCATGATACCGTCATCATTCTTTGCTTCGCTAACTGTTTGGAGGAAGAATCTTTGTTGCATTTAACTCGAAATTCAGAGTGCACACCTTTCTCTCCTGTAAAGAATCCTGAAGTCGCTACCTTATTAAGAACGAGAAGTATCCATCACGAAATACGGGATTATGCCTTATGGATTCATAGTAATAGTTAGTTCCGACGTTGAGATGGATTCAATTGAGACCGGTAGTGGTCGTCCGAGGCTAATGACGTCGTTGACGGGACATGATTAATTTCCACATCGATATAGTTAAAGGTATTTCTGGGTATGGGTTCGCATTTATCTGCGGAAGAGACGGTGTGAGAATATGTTCCGAGACCACACGGAGAACAGATGACGTCTCCGGATACTCCGCATCCTATTCCACATTTTGTTTGGGAAACACATGCCTTGCATCCGGATGAGATCCTTTTGAGAACATTGGGCCAATATCTGGGAGAGCAGACCAGATTCTATTGTGAGTGTTACACGATCGTCTTCCGTTACAACTTAGACAAGCGGGTAAATGATTATTGGTGAGATGTGAAGTACTCCGAACCACACGGCGTACATTGTGTGTTAGCGTCTTGCTAATGCGCATAATCTGAAGCGCGTATGTTCCCGGACACAAATTATGGCGTTTGTATTCGCTTGGTTTCTTTACACTTTCCATCGGATGGCATGCGGTGCTATATCTCTTCCGTTTATTATTAATATACATGAGAGAAACAATATATACGAGTATAATACGGACTTCATAATTTAATAATGTAGTAATCAAGTTGTCGTTCCTGTTTCCTACTTCTCCAATCATATAGATATTTTCTTTCTATCATGGATAATATTTGTAATGGTTCTTCCGGTACAACATACTGTTTAGATGATATTGCGCATAATTTCCGAGGCAAATACGATAGTTCAGATTGACCGATGGTAGACTCCTAATTTATTGAGTGCTTTGTCGACGAGTTTCATTTCTTACGCTCCATCGATAGATGGCACTGTTCTATGAGATCGTCGTACATGGGAAATGAAATGTGTTCGTCCGAATGTATGATTCCAAGATAGCTGTGATACCGTATACAGGTCGGTATGCTGAAAGATTCGAATCTCTTTGAGGCGACTTATGTCACGATGATGGAATCTATCTTACTGAATGATATATTTTCATATAAATACACTTTTATAGTCCGTTTAAACAGAATTTACTATGTAGTTCCGCGAATGACTCGTCCCTTAATAGGCAGTAGGCTAGTATCTTTTACGTAGTAATCGTCGTAGGGAGAGAATTCTGACATCTTGTAGAACAACGATTTAATCATAGGTAGAGATACTTTCAGTCTGTGGTGGATGATAAACATTCACAACATCCGCCTTGTATATGATGTTTCTGTTTTCAAACACCAAGTCGAATACCGTCTTAGTCGGAAGGTTGATGTCGTATCCGATGTATGAGGCAAACATTGAGTTGTTACACTTGAAAATATGGCATATAGTATTCGTCTTTCTGAATGTCGAACCTATCTAGTAGATACCGTAGTATATTGAGAGTGTATCCTTGATTATGTTTTTATGAATAGATAAAGTAGATGGCTGTCCTTCTTCCTTTTGTTCGCCAATTCGAGTAACATTGGCGAGAATATGACCTGTTGCACAATCGTTCCATGATGGGTGTACAATCAAGATTATTACGTATCCTCGAGATAAAAGAGCATACACCACACGAGACTATGTTTGGTATACTGTTGAAGGTAAGTGTGTAACCGCGTTAATGTTTGCTCCATAACGGCTATCGCGTAGATGAATTGCCGGCTCGCATCTTAGTGACTTAACTTGTAATAATTGCTTTGGTAGAACGTGGATATGTGTTTACAGTAGCAATGAAGAGAAGAAAGCGTTCATCCTCGTCGGCGCAATTAGGCTTCCATCCTTTGTACAGAACGCAGTAGTTTAAGCCCTACTGAATTTATATCTAAGATAACACAGCAATAGATCGGATGATTTTACTAAAGTCATCAATGGTGTCCGAAAGTTAGTATATCAAAGATCTTGTTATCGATTGATAGTGAATGAATCAGATAGTGGTGTCCTCCTTTTCATCCTTGCTATCAAAGCATGCAGTGCCGCAGGTAACAACAATATCTTAATACAGATGGATTTCAACCGTGTATTCATCGTATAGCAATGTAATGGAGAGTTACCCTGTTTATTCAGATCGCAAGTGTTTAATAACTAGCTTAACAGATGAGACGATGTATCCACATCAAAGAACGTAAAAATACATATGACAAACATTGCTGACAGAAACGTGACCTTCATTCTTACCTCGTCGTCCATAAATACGTTAGGTATGTACCATATACTGTCGCGAACGATGCTCACAATCTCGTCCATCTCGCCCATCTCTTATTATTTCCATCATTTACTTTTTTCATAATTAGAGATGTACGAAAGAAAAAAGAAAAAGAAAAGAAAACAGAACAATATATTTTTTAGTAATAATGTTTATGCGAGACATATAAAAATAAACTCCGTGTTTATGATAAAGCTGGTAAATGTTTTATCCATCTTTGGACGGAATCAGATTTTGTAATATGTCATGGAAACAAATGAAACAGGACATTATCAATTCCATGATAAACAATTATTTAATGGAGTAATAGCATTCCATGGGTAATTTCGAAATCAAGTTCGTCTGTATTAATCATTTGATGTCGGATTCTTTGCGTATTCTAGTCTGTGGCGTTTGCTTCTGTTTAAATAATATATCAAACATGGAGACGCCTGATATGTAGGCATTCTTCATTCTATTAATGTTCACTTATAGCGCTTTAGTTCCTTATGATTAATCGGTGATATCAATACTTACTTTAGAAGGAAAATCATCATCTAGATTAAGGCGCATCTGATACAGGCGAATAATGTCCAGGATATAGATATGCATATCTCATTAAATGCGTCAATCATAGTCTCAGAGGGATGGCAGCTAAGTAATAAATCAAACTTATCCCTCGTTTTGTTTTCTTCTTGGTAACTGCTTTTCTCTGGATGGCCGTATTGATTATCGAGCGTGATGTTGTAACACTCTGCCCATATTCCAATAACCGCTTTGCAAATTGTATATTATTGACATCGACCGTGTAATATAGTAGAGTTTTATTCTCATTATCGATCATATCTATATCATCCATGTACTTGCTTTAGTATATCAAATACATCTATTAGTATGGTTTCATAACAGTGATACCCGCAATTATTAAATCTCGATATCAGACCGTACATACATAGACGGGTCATTGTTAGATATGTGATTTAAGGCAGCCGCGTGTCCATATTTTCCACGATAAACCCATGACGTTTTAGCATCGGACGAGACATATTAACAAAGTAGCGTCGTGCAGAGGGATAGTTGTTGTCCGTTATCTAACATGCACGAAACTAATTCATATCGCCGTAATGTAAGTAGTTTAACATCAACATGGCTTGGTACGATGGATTCATCCTGTTCAACCCTTTAGAATGTTATCGATGATGTAGTGGTTATATTCTTGAATCGTACGGAAGTAATACTACGCATTACGTCGACAAGAGCATGACGTCTCTCAACAAGAAGATTAACGATTTGGCGTTCACATTATATGGGGTTACTGCAACGTTTAAGCTTAGATAATACGCCTCTTAATATAGGCTGACGTCGTATACCTTACACGTGTCCACATCCTTTATTAATAATAATTTAACAATCTCTATATCTATGGTTGAGAAAGACCAGTAGTATTGGATGGGTAAAGATCCTCCTTCGTCTCGCCATATGGATGGAAACATTGCGCTATGGATCAAACATTTAATTACATCCCTGGATAGAGATTGAGATTCTCTATGAGACGATATATAGTAATGAAGAGAGTTCTTACAATATTATCACTGTTGTATATACAGGTACGAAAGATGCAACCGGTGCTGTAAACATCTCTGATTTAATAGCCATAAGTTAGTTTCTGGTCTCGGATTAGGCGTCGTTACGTATATATCCACCCAATCCGATGAATTCATTGATTGTATAATTGTACTTGGACGGACGTATCCGTTTATCCACAATTAGGTATTTGAAGACGTAAGTCGATTATCCGAAGACAGATCGAAATCATTTATATTCGACTTGAGTTCGTTAGAGGAATTCGAATAGCTGGATATCGCAGGATGCACAAATCTGAGATTTTGTATTTACTTCCATGTTTACTGTATGCTCCTAGCGGAGTTAACCTTCGTTGTTTCTACAAAGTCTCGACTCCGCGAGAGAGTAACAGTCGAACAATCTTAATGTCTGTTATCGCATTTATTGGAGACGTAACAATAAAGTGCATTGTTTCTCGTTCATCTATATATGTTTTGATAAGTTGTGACACGTTTCAATCTCTAGTTTATTTTTTTTGTACGTCACATCTTCATCCAGTAGACGACATAGAATAGTGTACCTTCTACCACAATAATCCATAGCTATTCTTGGTGCTAATATTCCTATTTCACGAAAAATGATGAAGGCAATCTGGCCTCATAAGATGATGATAAAAAAGTGTAGTGAGAGAGCATGAAGGAGATTTAGTATTTAGCAGTGCGGATGTTGATCCAAGAGGGTGAGATAGTCGTTCTCGTCTGTGAATCTTTCTCGCAGCATAAGTAGTATGCTCGATATATTTATCGTTGGAAGACTCTTCCAGAGACGATAGCTGATTGAGTACAAAGTCCAATGATTGCATCGAAGTTCTCGGCGGTTTTCAGTGGAGTCATTTCTGATGAAACATTTAATGATCTCACTTGCCTGTCGATATTGTCCCACGGAAGTGAACTCTTCCAACTCACCACCAAAGAGCTCCGTTGCATCAGTTCTGAAAGAGATGAGAAGCCTGTAGAGAGACCCTGCGCTTTCTCTATGGGTCCATCTATGAGAAACCCACAGGATGTATTCAGTCAGACAATGTTCTCCGACGTCGGCCACGGTATTCAGGGAGTCCCTTAGTATGTGGCAATGACAGGGTCTGAATGTTACCAAGGAAAGGCCATTGTAAAGGTAGACCTAGCCGTTTATGCTAATAGAGGCTTTAATTTCATTTTTAGAATGGGTTGTGGATGAGGAATGAGAGTGATATCATATTGAGAGCGTAGTTATGTAGAGGTGTATTTCCTATATTATTTACTTTCGGTTTATTTTACCAACTCTTTTAATAAATTTCTTTTCACGATGCATCTTAGTTGAATGATCGCGTTTTTCATAAGTGGACATATAGATGCAGAAGTAATGAAGAAAAAGTATTACTCTATCATCTACATAATTAGGGTCTGCTCCTTTTTAACAACTTATAGCATGTAGTAGTAGTTTACTAGTTCAAATCAAGTCTAGAATATATAGTGGATTAATATATTTTTATATTAGCTAAAAAGCTTTATCTATACTCATCAGAAAGCATATCATTCTCAACTTCATCATGAGTTAAATATTTGTGTAATGGATGTCATGAACATTAAACGTATTCATGACATACCTTTAATAGGTTTTTAAACAGTGATGATTCAAATCCTTCCATTCATTAGATAACAGTGTAACGGAGTCAAATTCTCTTACTAGTTTGTTTATATCATAAGCATCTCTACAAACAGTCTAAAACAACAATAGAGAAGACGGACAGATTCTTTTAACGCATAAATGACACATGTTATCAAATGATATTCGCTGGATGAATTATTATTAAACGTAGTTATGATAAATGATCTCTAACGACATCTCTCGCTAGAGATAAAATCTAGTATCAAGAACATTAAACATCTTTGCATCATACTCGCATAGCATAGTTTTTCATAATTATCACAATATTTAAAAAGACTTATTCGGAAAAGTATTTTACATGTATCATCGATGGAGATCCATATGAGGAGTCACTAGTTCTTTCAGTAGTAATAACAGTGCTATCATTCGATAGTATAATTATATAAGGCGTAGAAGGTTCATATGTTGTTGTAATTGAGTAACCGTTGGTAGTTCTGTGGAATCAATAATTATACTAACAGCAAATAGTATATAATATATAAATATGTTCCGTTGATATCATTATTTTAATGAACCCATTTCTAACACCCTGTTATATCGTCCAATTAAATGTAGCCAAACAATCTACCATGTTCTCGATTGACTACTTGTACGGTAGCGACGCTGACTATCTTTTATTGTCTCTGCTCCAATTGAATGTCGCGATACAACGCAGTTTCTTCTTAAGTGCATGTTTCATATCACCACGAACATAAGTCGGCAAGATATATAGCCAGAGATACTTGTAATCATGATTGCTTGGTCATAAACAAGCCCGTCAATAATTGTATACATATTCAGTTTAGAGCAAAATAATTAAGCACAATAGCGCTTAATCTCAAAATATGTTATGTTTATTTTTTCATATTAAACATACTGGTTAAAATCCTCTAAAGGCTGATCTTCTATCTATAAATCAAGACTACCATTTAGACAGTGGTTTCATGTTTATAAAATGTTCTTTTGTGTGAATAAGGAATATGTTTAATCAATAATCAACCATCGACCCCATTACGATAGTATGCAGGTTAAACCCCATTAGAGAGGTACGTGTAATCAGTCTCTCCAGTTTTAGTATTTTTATAAGTCATTACTACATAAACGGCTTTTAAACAGTCTCCTCGATAATAAGCCATATCTGAAATTCTATTAAATACTCGAGTCATTTTACGCACGGTCAAAAAAGTAAGTAATGCCGACGACTTCTACATCTCTATAGAACACCTTAGAATACTCATTTTCTTTGGAAAATATCCTCAGACTCTGATTTAACAATGCACGACCTATAGTAAAACCGCAGACCAAGTTATATTAGTCAATGGTATATCCAACCATCAGGTGTGGATAGTCCAGTCTTTGGTATCGATAGTGTAGTTATTGAACTGAGAAGTTACCGTATAAATTTTTGGTCATCTCTAAACAAGGAAACTAATAATTCTACACTATTGAACGATTTATCTTCCGTAATGGGTGGACTAGCGATGATGAAGTCACGAATATAAGACACGCTATTAATCCGTATATCATCATTTTGATATTACTTATAATAACGATTTGTTTAATTTTAGTTTATACTATTAATTGTAAATGATATTGTGATAAAGTCTATTTGTTTATATTTATTGCGGATAGCAGTATTTCCATAAACGCAGCAAGGAGATGTCCTAGGGTTTGACTATACATGTGAATGATTGACGAGGTAGATCTGTAGCCGATAGGTTCATCGAAAAGACTGTAATACTCCTGTAACGGGCTGATGCTTCCTAGGCACTTTCATTAATGACGAGAACGTCGGGAATTTTTCATAAAGATAGTTTCTTATTACATCTCATAAATATCCGGAATTATTGCAGTCCCGAATTTCTTTAAAATCGAGAAATTCAGATTATACGGTTCTATTATTTCCAGACTTTTAAACTTTGTATTATATTAAGTCTATACTTTTGGTAGATAACATACCATTAATCATTTTAATTTTTCATCCATGCTTTTATGTTATACACCTCTATCGTCCAGTATAATCACCATTACTAGAGTTTCTCGTATGGAATATCGATAATAGAGACCCTCCGAATAATTCATTAGCGGCTATTGCAAGTTATTCTCGGTACCACCATCATATCAACACTGCACCATATTCTTAGAAATATAAAACTTGTCTGTATATGTAAGATGTGTTTAGAAAAGCCGGATATTTCACATTGCTTTGTAAAATGGACGGCGCTAACAACTGTCATATTTATATTAATGGATAGCGGATTAGTCAATAAGGAATTACTCTTACCATTTGTCACGTCTTAACCCCATTCGCTGATTAGTTCCTTTGTTTGGTTAGCATTATTAAAGCTTACAGCTTGAAAATCGTCTTTATTTTTGTAGGAAGGAGGCGTGGAATCCCGATACTATCGCTACCGTATATTTTATTTGCGGTAGCGCTAGTGTCTGCACAATACGAATATCTACGTCCATGTCATTATTGCTCATCGGGTGTATTCTCATTCATATTCTCTATATATTTTGATAGTTGTTCAGCTGTAGAACCAGCTGCCCATGATTTAGAACAGACAGTAGATAAAATAGAAAACCGAGAAACAAAACATTTTCATCAGGGTGTTTTACGATTAGTTCTTTAAAGATATCCATGGTATGTGGACCAAACAATAACGATAACGATATATATCATAAATAGCCAATGTTAAATTTCAGTTTATGTTTGGTACCCCGTATTCATACATTTCAATAAATTGGTATTGCGTACACAATCAATCATATTACATACCATTAATAATGCAAGCATAAAAATCGTTAGTAGATGTTTCAACATAGGTTCCGTAAGCAAAGAATATAAGAATGAAGCGGTAATGATAAAATCAATCGTTATCTAAAATGATCATACTCATTTATTTTATTCTATTATATTAACACATACATTTTTAAATAACAGCAACACATTCAATATTGTATTGTTATTTTATATATTATTTACACAATTAACAATATATTACTAGTTTATATTACTGAATTAATAATATAAAATTCCCATCTTGTTATAAACACACACTGGAGAACAAAGCATAAACACAGAATCCATCAAAAATGTCGATGAAATATTCGATGTTGTTGTTCGCCGTTATGATAATCAGATCATTCGCCGATAGTGGTAACGCTATCGAAACGACATTGTTGCAAAATTGAAAACGCTACAAACAGATATTCCAGCTATCAGATTATGCGGTCCAGAGGGAGATGGATATTGTTTACACGGTGACTGTATCCACTAGAGATATCGACGGTATGTATTGTAGATGCTCTCATGGTTATACAGGCATTAGATGTCAGCATGTAGTATTAGTAGACTATCAACGTTCAGAAACCTAAACACTACAACGTCATATATCCCATCTCCCATTATGCTTCATTAGGCATTATTATTGAAGCGGCGCACGAAAACGCGAAAGCGTTTCACGATAAATGCGAAACTAGGGAAACACGATAGAACTGGAAAGGCTTCACGTAACTGAACGAAGTACAACAG

### Isolate 5

ACGGTACTTCGTTCAGTTACGGCGTATTGCTCTTGTCCAGGGTTTGTGGCCTTGTTTCCGCATTTATCGTTTGTGTCTAAACAATGAATACGCTTTATACGATTTCATCTTGAATAGATCCATAAACGGGGGCTCCATGAATATCGTGTTTGAAGTAACGTCGTCGGATACATAAGAATACTTTGTTCAATTACGGCTTTTGGATTTGCTCCAATATCATCCATTCTTCATCACTCCGTCGAAATCAGCATTTTTGAGAGTTGGCAATTCGGGAGATATTTTGATAGTATCTCCTTCGGTAGCTCCTGATAGATGAAACGATGACGTTTAACCATCTATGTAGAGACGGCTGTCTTCCAAAATAATGCCTTATATTCTTGAACGGCTTCTTCACTGCCCAATCACCAGGCAATAAATGTATTTTATTTTAATAAACTTTCCTTGGCATATTCTTGTTAATTGGTTTAATCGTTTATTAAGTAGAATTTAGCAGTTTGAGACGCTAATAGTTGTTTAACTTATCCTGTAAAGGCATTAACAAATATCTTTTCTGTAGTGTATTTCCTGTATATGCGGGCATTCCTACCTCGTAACGGTGATAGATGTACTGGGACCAATTACAGATCTAGCGGTCTGATCTTTTCGTCGAGGGCGACAATATAACTTCTAATCTTATTATTTTTGCGGATGTAATATATGATAAATTGATACTGGAAGTGTTGTAGAAATAATTTTAATATCATCGTATTCTGTACTGCCTTCTGGATAACCTGTTCATCCAGCATTCAAGTTACAATTCTTAACGATCATACCTAATAAAGTAAGTTAATTCATTGGTCTCTTTGGGTACATACTATCTATCAAAAACTAATAGCCGGTCTAATAATCAGCGGAGGGATGGAAAGTAGTCGCAACATAAAATAAGTTGGCTGGATATTGATGGGTATAATAATGGCTAAAACTTTTTCATGAATAGAAATTAACTTTTTGATAGATGAAGGAAAGTGGAACGTTAATATCATCCAACTTGTTGACGAAACAAACCTTTCTTGAAAAGTAATTTTTGATACGGTTGCATACATTCGCTGTTCCAACATGAGCTTTTCTTGGATAATATTTATCCTTTAATGCTCCCAGAGCGTGTCCCATTCAACTCTTTAGGTTAATATCGTCCAGACCGGTTCTCGTGAACACCAATAATCCGCAGTGAATACATATATGATTCGGTAAACGAATAATTTCTGAAATAAATTCAGGCTTAACTATGAGGTTTTATAAATACTTACTTACCCCAGTGACCAAGAAACATTCAATTCCGTTTCCACAAGTCTTGTAATGCCCCATCCATAGCACCTAGTCTACGCTCTTTAACGGTACCGATATCGTCGTCATTTTTTAACACTGACTAATGATAATATCTGTATTAATCTCTTTTGATCATATAGACTATATACGTAACTAGAGATTACAACCATTTTTATCAAGTCAGTTTCTTTTAAAGAACCAAAGTATACAATCAAATTTCCCTTTTTATTACAACTATAAAATAATAGTTATATTTACACACTTTAAATTTTTTATCATGACAAGGACGAACAAATTTATGCATTCTGTGATGCTAACAAAGACGATATACGATGTAAATGTATTTATCCTGACCAAAAGCATAGTACGGATAGGAATAGATACAAGATACCCTATTATTATTGTTGGTACGAATAACCATGTAAACGAAGCGATGCGTTGTTACCAGCCTCTTTAAAAAATATAACAAAATGCAATGTGTATCGGATTGTACCATTTCATTGGGAAACGTTTCCATTACAGATAGTAAAATTAGATATGTAAATAATGTTTGTGATTCCCAAACGGTAGCTACCGAGAATATAGCTGTCCGCTATTTAAATCAGGAAATTAGATACCCTAATTATGATTGATGGCTTCCGATTGGATTACTAGTTAGCTATTTTAATATTACTTTTCTAAGCAAGATATAGAATTATAAGATACCAGATATATATTGTGATTGTATAATGTTCTTATCTCATCTCTACTAATTGATTAATCAGCGACTGAAATAGCAGATCTATCGGCTATCTCTACTCCAGTTACCATGTTGTACTTGGAAAAATCTAACAATTTTTAATGGTATATTAGGGCGATAGAATCTTGACAACTATTTCCGTCTCCCAACATTTTGGGAAACGAAAGTCTTTTTTACCATCGCCTGTTTGTAGTACACTATTAACTATATTAGTTTCTGTAAGTATCTAGTCTAACAAGCTGAAATAGAAGTCGAGATATTACCACATCTATGTATAGAAAATTTACAGAAACTTGTTATGGGATGATTCGTTAAATACCTGAAACACAGTTTCCATAGAATCAATTTTATGTCTAACAGTTGTCAAAAAGTAACTAAATCATTATACTCTATATCGGTAGTATATCTCAGTAGTACGTTTCTATTTATTACGCGTACGGATCTCTTGCTATTTCTGTTTTAGACATAGAATTTTGCTAGATATTTTTACGTTGTATTGGTTCTTGACTAACTTTTCAGTTGATGTTGTTGAATATTTAAGAAACGAAATATAGGTGTTGTAGAAATAGTACCTTTGCTTTAGTAGTAGGAAATGTTTTATTGCAGTACACGGTCCTCAGCATAAAGTACATGTAGAAAATAGTCATATTCTGATTAGGATAATCAAAGTTAACAACTACTTTGTTACGGACGATCTTATTAAGGTAGTACATCTTTTCATAATTTACAGCGTCTGATTTGGTAACTCGAGGTCCAGTCTCATGTTCTCTTTCGGTATAAATACTTAATAATCTCATTCAGCTGAATATGAAGGAGCAAGGTTGTAACATTTTATTACAGTGTGGGATATAAAGTCCTTGATCCAGTGATCTGGAAACGGGCATCTCCATTTGGAGCTAGATGCCACGGGTTTAAAATACTAATCATGACATTTTGTAGAGCGTAATTGCTTAGTAAATCCGCTTAACACTAGGTTCATTTCCTCCTCGTTTGGATCTTATCAGAAATTAAATAATCTTAGAAGGATGCAGTTGTTTTGATGGATCGTAGATATTCCTCATCAACGAACCGAGTCACTAGAGTCACATCACGCAATCCATTTAGGAATAGGATCATGATGGCGTCCGTCAATTAGCATCCATTTGATAATCATTTCTAAATTGCCAAAATGATCTCTCAAATAACGTATATGTGTACCAGGAGCCGATCTATGCATACACTACGGTGGCACCATCTAATATACCGTGTCGCTGTAACTTACTGAAAAAAATAATTCTCCCTGAGCCAATAGTTTTAACGCTGTCCTTGATACGGCAGTTTTGCGACCTCATTTGCACTTTCTGGTTCGTAATCTAACTCATTATCGAATTTCACTCAAAATACATAAACGGTTTATCTAACGACACAACATCCATTTTTAAGTATTATATTCTAAAAATTTAATCAATGTTTATTTTTAGTTTTTGAATAAAAATATAATATTATGAGTCGATGTAACACTTTCTACACACCGATTGATACATATCATTACCTCTATTATTTCTATCTCGGTTTCCTCACCCAATGTTTAGAAAAAGGCCTCCTTAAAGCATTTCATACACACAGCAGTTAGTTTTACCACCATTTCAGATAATGGAATAAGATTCAAAATATTATTAAACGGTTTACGTTGAAATGTCCCCATCGAGTGCGGCTACTAACTATTTTCCTTCGTTTGCTGCCATACGCTCACAGAATTCAACAATGTCTGGAAGAACTGTCCTTCATCGATACCTATCACGGAGAAATCTGTAATTGATTCCAGGGCATCCACATAGTTTAGTTGCTTCCAATGCTTCAAAAATTATTCTTATCATGCGTCCATAGTCCGTTCCATTATCTATTATCGTTAGAATATTTATAGTCACGCATTTATATTGAGCTATTTGATAACGTCTAACTCGTCTATTAATTCTGTACTTTTACCTGAAAACATGGGGCCGATTATCAACTTCAGATAATGTCCGCCGTTCATGATGACAATAAAGAATTAATTATTGTTCACTTTATTCGGCATTTAATATATCCATCACGTTAGAAAATGCGATATCATGACTAATCACAAGTCGAACAAATATCCTTTATTAAGTTGACCCTTCCATCTGTAACAATAGGGACCAGCGTTAACTGGTTTTAAATCTTGAGAGTCTGTGAATTTTGTGGTGTCTGTATTCCTCTGAAGAGATTCATAACAATGACCCACGGCTCTAATTTGTTTTGATTGGATCAATAATAGTGTAGAAAGTCTAGATATTGAGTGATTGCAATATATCAGATAATGAGAGATTCATCATCTTGACTAGCCAAATAGCAAATGAATCATCCTCTGCGAGAACATCGTTAAGGATACTGGTTGTGATCCATTTATTGATCATAAAAGCTTTGCACAATCTTTATACTATCGGTTTACTATTTATTGATAACGCAGATGTTTGAGTTGTCATCCATGGTAATCCATAGATCATTAATTTATCGTCTTCACGCTAGATTAGCACGTCGTAATCTGTCAATAGAGGATCGAGGTATTTTTCGTTTTGACGCGAAGAACATATTTAATTCAAGATCTAAAAATACATATATTAGGAATGAATACAACAGGAATACTTCAAATAAAACTATTAATCTGTGTTTATAAACGCAATAAAGGAATGTTTAAACGTGGGCTCTATAAACTTTTAAGAGTTAAAGTATACATTAGGAACATTCTCCATTTATAGTAATCAATCCTTTGTCCGGAATATCTGTTAGAGAATATTCTTTTAACACATTCCAATAGTCCCAGAAAATCATTTAACCTAATAGGTTCTTGAAGAGGCTAGAACTATACGATTCTAGTTCCTTATTAGTACTAAGTTTCTCTAGTTTATTTTTATTGGTGAATCCGTAAATGGCATTCAATCTCATGGATGTGGAAGGAGGAATACACATTCCAAGGACATTATGAATTCCGCATATTTGGAATATCAAAGCCTTGTTGTCTCCGATAGGTGTATACTCAGTCGATGCGGATTCCATATTTTTCTTTATAAATATTAATCTTTTACGAGTTTCGAAAATGCACAAGATGAGTGATGATCCTTGACGAAATAAGGGGCATAGCTTCATTGTTAGGATCCAAAATGGTTTTCAATGGCTTCAGCAGATTTGGTTTATCATCTCCCTAGACCCAAATTGTTTTCTTATATTATACTGTTGTAGTAAACATTGCTCCTGTGATATTTGCCTGGAGTATGAATCAACTCCAACTCATATTTTGACATCGTTCTAGGTACTATATTAAAATAGAAAGCATAGATCTTGGAAATTTGGATTTTGCGCCGGCAATAACCATTGTAAATCATCATACTCAGATATGCTTCCTGCAAAAATATGGTATCTTCTTCGATGAGGTTTTCTAGCAGGCGCTCATTTAGAAGTTTTTGTGATAAATGAATACCCATACCGATGTTTACAAACGATAATATAGACAAAAATCCAACTAAACGGTGATAGAAATATACCAGGAAGAAATGAAAAGATTGTGGCCAAAATAGATTCAACAACGATAGACCAAGGCCTAAACCAAAGACTACAGCCTAATCAGCCACCGAAACAAGATAATAAATGCAGAAGTGGAGATTTTATCAATATTAGTTGTGTGCCTACGAAAAGGAATATTGCAATGACGGATATCTATCTCCTGCCTATTATATGTTAAAACAGGTGGATGATGAAGAAATGAGTTGCTGGTCAGAACTATCGTCGTTGTGAGATCCAGAAGGCGGTAGGATTTCCTCTATTAAAAGGCGGCTAAACGTATTTCTGTGGATCGATGCTATATTTTAAACAGTTTAAAAAACAGTAAAGTTGTGAAATTAACCCACTGGGATTAAATGTTTAAATGATACTGTTATTTTCAAACTGTAGTTATTTTATATTCAATGTATAAACGTGGCATATATTCTAACAATTTGTTTTGATCTGGTTTCTATTCCCAGAACAGACATTATTTTTTCTGTTAATCAATTAATGTTTAACATTTGTGCAGACATATTGGTAGTTCTATCTATTTACGGCAACCGGCTCTATAGAACAATCTACCACAGTCGTGTTACTTAAATTTCATGCAGCCATGAGACAATAGCCGTGAGGATATGTTACTCAATTACTTTTTCGGTGGTTGATAAAAAATCACATATCGCTATTGACCAAGCAGAACGATGGATATTCTCAGAGTAAAGAAAAGAAGTATGCTACAGGAGCACCAGTAAATGAGTTGTTGAACCTGGTACACTGGTATATGTGCCCAAGAAGATTATTACTTTATGAGCATGTCACTCACCGATGTGTCAATTAGCAATAATGTCAGGTATTATTTTCCACAGATGGAATAGTGTTAGAAATAGAAGACTTTAATATCAAGCATTTGTTATGGCAGGTGAATGTTTGTTAGAAGTCAGTCTAGTACTATTATGATATAAAGTAATAAAAAATAGTTAATGTGATGACTAGCGCCACCAACGCCAACAACATTTGATAATTTCTACCTAGACGTACCGTAAAAATATATAAATTACTATAACAAATAATGATATATCAATAAACAACCTAATTAATGGTCGAGTATAGCGAGGACATTGATGCTCTAGACCGTGTATAACAAAATCTACAAATTTTTCGTCCGCTATATTTTGTTTCACTATATCGTCGATGATCAGCGACCAACTTCCATGTTAATCTATTAAAATAACCATCAGCGTATATTTTTTCGTTTGCATATCCGTGGTAGCAATAACCATCGGAGAAGTTCTAAAGAATGTGTCATGTAAGTAGTCAATGGACGTTTCCTTATTGGCTAAAATAAGTTTGATTTTATCATTGGTGGATGTGAACAGCATACGCTTGTAGATACATAAGCACTGCCAATATTATAACACCGATAACAATCATATAAAAACTGAACTCCTGTACCAGCAACTTGTCTAGGTGCTATTTAGGTAGTGGCCTTGGTAGTCCAATTGCATCAACGCCTTAATGGCACAATTTCCTTTGCTAGATCCTGTTAATAAGTTCCAAATTTGTTGGAGATCCTGGGGCTCCTTAACATTCATCTATGATTACGTTTGTATCTTTAATTTGTTATCGACGACCATACTTCAAATTTACAGGTTGTTTTACATAATTTTCAAAATCTCTGGCAACAGTGTTTACACTCGTCTGAATGTTTAACTTGCAGCAGTAAACATAGCTGGCGTATGCTTTGTTCCGGTGTTAATCCACTACTATGTTTCTGTAGCGGCTGATAACCACAGCATCAACTGAGCATCCGCGTCCTGAAACACATATTTTTAACTGTGAGTTACATCCATGGTTTTGGCCGGATATAAAATTTCCGATTTCTATATCACATTTGTTTTGAACATAGCATTCACTTCTTGTTCTAATTTAGACGAGATACGTTCAGCTAGGTGTATTCACCGTCGTCTGGCAATAGCGCTGCGGCTTTTCCATTTTAAATAGCTACAATTAGTATCCATGTGCAAGAGATAATAAACTGATCAAATGCAATTTTAGGTCGAACGCTTAATCCACATGACAATCTTAATGAGAGTTTATCCATCTGTAAGTTGTTCACGCTAGTATTACATCGTACAATATTGCAAAGTCTAAATTATTATAATTACGTGTTAGCAAGAAATTAACATTGGCATTCGAACCTCTGGATCCCAACATTCTCGAGGTTCCGCATTTTAATGACTCTTCTAACTTATCTCTAGTGGGATAACTACATCTCCTTATATTTCTGTTTAAAGTCCGCAATTTATTTGTCTTAGAATATAATCGATCATCTCTTTGCTATCTTCTGTATTGTGTGCGTAAATGATGCAGAAATGATTCACATATTGGTACACTAGCATCTTTACTACATAATGTTTGCATCTTATTATACAGATTTATTAACGATTGTTGACCCTCTACAGTTCTATTACTCTATTAAAATGAACCGATCCACTGATGGCATATGTTTCTATCGAACGTATCCCCTGACACCAGTCGAATAGAGAAATCAGCATCGCATTTCCAGTGTCGTGAACGTCTGGCCAACGTTCTAATACTGCACCCTCTTCATACTTGTCTGATATCTTTCCATCCGCTTTTTCCAATAATGAGTACGATTAAAGTGGAGCAGCAGTGGGTGCGGTTCCACGACAACCGCGATTGTCTTAACAATGCCGAAAGACCACCGGGTCCTGTTGACTAATCTAAACTCACGGATATCTTTTTCTTTACTAGCTTTTCACCAGCAGGTTTGCCGCAGAACGCAACTGGAAGCTTCGAGGGGCTCGCATCTCTCCTTCACGCGCCCGCCGCCCTACCTGAGGCCGCCATCCGCGCCGGTTGAGTCGCGTTCTGCCGCCTCCGCCTGTGGTGCCTCTGAACTACGTCCGCCGTCGCTCCTGGGCAGCAGTGGATCCCTGCGGAAGCTTCTAAAAATTGAAATTTTATTTTTTTTTGGAATATAAATAGTGAGCAAGAACGAGGAGGACAACATAGCCATCATCAAAGGAGTTCAGCCTGCGCTTCAAAAGGTCACATGGAGGGCTCCGTGAACAGCCACGAGTTCGGATCGAGGGCGGGGCGAGGCCCGCCCCCTACGAAGGCACCAGACCGCCGGCTGAAAGGTGACCAAGGGTGGCCTGCCCTTCGCCTGGAACATCCCCTCAGTTCATGTATACGGCTCAAAGCTACGTGAAGCACCCGCCGACATCCCCGACCGCGGGGCGCAATGTCCTTCCCAGGGAACTTCAGAAGTGGGAGACGCGTGATGAACTTCGAGGACGGCGGCGTGGTGACCGTGACAGGACTCCTCCCGGGACGGCAGTTCATCTGAGTGAAGCTGCGCGGCACCCAACTTCCCCTCCGACGGCCCCGTAATGCGAAAGACCATGGGCTGGAGGCCTCCTCCGAGCGGATGTACGGAGACGGCGCCCTGAGGGCGAGATCAAAGCAGAGGCTGAAGCTGAAGGACGGCGGCCGCACTACGGCCGCTGAGGTCAAGACCACCTACAAGGCCAGAAGCCCGTGCAGCTGCCACGCGCCTACAACGTCAACATCAAGTTGGACATCACCTCCCACAACGAGGACCTGCCACCATCGTGGAACAGTACGAAGACGCGCCGAGGGGCCGCCACTCCACCGGCAGCATGGACGAGCTGGCCTGGAAGTCTAGATCTGCCACCATGCTTGTTTTGTTATTCGTTGCCCAATGCGGGTGATGTAATAAAGAGGGCAGTATACGAAGGATTATGCTCTATATATTTATCTTTGACTATCCTCACTTTATGAAACTATCTTGGCAGAGAGTGTTGAATGCATATGGATAGATATGTTGAATATGAGGGATAAACTGGTAGGGAAAACTGTAAAAAGTTAAAAGTGATTAGAGTTGATTATACAAAAGGATATATAGATGATCAATTACAAGGATGTGGAGCATCAATAATAGGCGACCGCCTAAACTTGTTTTATTGCAGCTTATAATGGTTACAAATAAAGCAATAGCATCTAAATTTCACAAATAAAGCATTTTTCACTTTTGCAAAAATAAATTCCAAATTTGTTGGAGATCTGGGGCTCCGTAAACATTCATCTATGATTACGTTTTGTATCTTTTAATTTGTTATCGACTGACCACTTGCTAAATTACAAGTTGTTTTACATAATTTTCAAAATCTCTAACAACAGTGTTACTCGTCTGAATGTTTAACGCAGCCAGTAAACATAGCTGGCACGTGTCTTTTGTTCAGTTAATCACTATATGTTTCTGTAGCGGCCTGATAACGGCATCCAACTGAGCATCCGCGTCCGCAGAGCACATATTTTTTAACAGTGAGGTTACATCCATGGTTTGTCGGATATAAAAATTTCCGATTTCTATATCACATTTTGTTTGACACTAGCATTCTTCTTGTTCTAATTTAGACGAGATACGTTCGCTGAGTGTATTCACCGTCGTCTGTATGCTTGCTGCGGCACCATTTAAATAGCTACAATTATCCATATTACCAAGAGGATAATAAACTGATCAAATGCAATTTAGGTCAGTGAGTGTTTAATATTATGTTGAACTACTTTTGCTCTATTGACGGGAACAGTAGAAGAATCTATCACTATTGCTTAATCCGCGACAATCTTAATGAGGAAGTTTTATCCATCTGTAAGTTGTTCGCGCTAGTATTACATCGTACAATATTACCCAAGTCTAAATTATTATAATTACGTGTTAGCAAAATTAACATTGGCATTCGAACACTCTGGATCCCAGCATTCTCGAGGTTCCATATTTAATAGCTCTTCTAACTTATCTCTATTAAGGATAACCCATCTCATATATTTCTGTTTAAAGTCCGCAGACTGTTGTCTTAGAATATAATCGATCATCTCTTTGCTATCTTCTTATTGTGTGCGCGTAAATGATGCGAAATGATTCACATATTGGTACACTAGCATCTTTACTTACATAATGTTTGCATCTTATTATACAGATTTATTAACGATTGTTGACCCTCTACAGTTCTATTACTCTATTAAAGGCTGAACCGATCCGCTGATGGCATGTTTCTATCGAACGTATCCCCTGACGCAGTCGAATAAATCAACATCGCATTTTCCAGTGTCGTGAACGTCTGGCCAACACGATTCTAATGCTGCACCTCTTCTATTACTTATCACTGATATCTTTCCATCCTTTTCCAATAATGAGTACGACAAAGTGCATTGTAGTGGGTCGCAGTTCCATTATGCGATTGTGTAACAATGCCGAAAGAACCACCGGGTCCTGTGTTGACTAATCTAAACTCTGGATATCTTTTTCTTTACCTAGCTTTTTTCGCTTGAGTTTGCCGCCAGAACACAGCTGAAGCTTCGAGGAGACTCGCATCTCTCCTTCACGCGCCCGCCGCCCTACCTGAGACCGCCATCCACGCCGGTTGAGTCGCGTTCTGCCGCCTCCCGCCTGTGGTGCCTCCTGAACTGCGTCCGCCGTCTAGCTCCTGGGCAACGGTGCGGATCCCTGCAGAAGGCCATCATCCAAGGAGTTCATGCGCTTCAAAGGTCCACATGGAGGAGCTCCGTGAACGGCCACGAGTTCAGTCGAGGGCGAGGGCGAGGGGCCGCCCCTACGGGGAGCGAACCGCCAAGCTAGAAGGTGATAAGAGGTGGCCTGCCCTTCATACAGGACATCCTGTCCCTCAGTTCATGTACGGCTCCAAGGCCTACGTAAACGCACCCGCCGACATCTGACTGCAAGCTGTCCTTCCCCGAGGGAGCTTCAGAAGTGGGAGCGCGTGATGAACTTCGAGGACGGCGGCGTGGTGACCGTGACCCAGGACTCCTCCCTGCGGGACGGCGAGTTCATCTGAGTGAAGCTGCGCGGCGCAACTTCCCCTCCGACGGCCCCAGTAATGCAGAAGAAGACCACATGGGGCTGGGAGGCCTCCTCGAGCGGATGTACCCCGAGGACGGCGCCCCTGAAGGGCGAGATCAAGCAGAGGCTGAAGCTGAAGGACGGCGGCCACTTTCCGACGCTGAGGTCAAGACCACCTGAGGGACAAGAAGCCCGTGCAGCTGCCCACGGCGCCTACAATCAACATCAGGTTGGACATCACCTCCCACAACGAGGACTACACCAATCGTGGAACAGTACGAACGCGCGGGGCCGCCACTCCACCGGCAACCAGACCGAGCTGTACAAGTCTAGATCTACCACCATGCTTACATTTTGTTATTCGTTACCCAATGCGGTGATGTAATAAAGGAGCAGGAAATTACGAAGGATTATGCTCTATATATTTATCTTTTGACTATCCTCACTGAAGCTAATATTGGCAGAGTGTTCAAAGATGCATATGGATAGATATGTTGAATATAAGGGATAAACTAGTGAGGAAAACTGTAAAAGTTAAAGTGATTAGAGTTGATTATACAAGGATATATAGATGTCAATTACAAAGGATGTGTGAACATCAATAATGACGGCCACCCTAAACTTAATTTATTACTTGGCTTATAATGGTTACAAATAAACAATAACATCTAAAATTCTAAATCGTATTTTCACAATAAATTCCAAGTTGTTGGAGATCCTGGGGCTCCGTAACGTTCATCTATGATTACGTTTTGTATCTTTAATTTGTTGTATCGACGACCGCGCTAGAGATTACAAGTTTGTTTTACAGGTACTAAAATCTCTAACAACAGTGTTTACTCGTCTGAATGTTTAACACAGCAGTAAACATAGCTGGCACGTATGCTTTTTGTTCCGGTGTTAATCCGCTATATGTTTTCTGTAGCGGCTGATAACACAGCATCAGCGCTGAGCATCCGCGTCCGCAGGCACATATTTTAACAGTGAGGTTACATCCATGGTTTTGTCGATATAAAGAATTTCCGATTTCTATATCACATTTGTTTGACACTAACATTCGGCTTCTTGTTCTAATTTAGACGAGATACGCGTTCGCTGGAGTGTATTCACCGTCGTCTGTGTCTTGCTGCGGCCTCCATTTAAATAGCTACAATTGGTATCCATATTGCAAGAGAGATAATAAACTGAGTAAAATGCAATTTTAGGTCGAACGAAGAGTGTTTAATATTATGTTGAACTACTTTTGCTCTATTGACAGGAACAGTGAAAATCTATCACTATTGCTTAATCCACATGACAATCTTAATGAGGAAGTTTATCCATCTGTAAGTTGTTCACGCTAGTATTACATCGTACAACAGCCCGCAAGTCCTAAAATTATTATAATTACGTGTTAGCAAGAAGTGCTGTGGCATTCCAGACACTCTGGATCCCAACATTCTCGAGGTTCCGCATATTTATTGACTCTTCTAATAACTTATCTCTAGTGGGATAACTACATCTATATATTTCTGTTTAAAGTCCGCAGACTGTTGTCTTAGAATATAATCAATCATCTCTTTTGCTATCTTCTGTATTGTGTGCGCGTAAATGATGCAAAAAATGATTCTATATTGGTACACTAGCATCTTTACTACATAATGTTTGCATCTTATTATACAGATTTATTAACGATTGTTGACCTCTACAGTTCTATTACTCCCTATTAAAGGCTGAACCGATCACTGATGGCATATGTTTCTATCAGACGTATCCCCTGACACCAGTCGAATAAATCAACATCCTCATTTTCAGTGTCTTGAACGTCTATTAACACGATTCTAATACTGCTTTCCTCTTCATACTTATCTGATATCTTTCATCCTTTTTTCAATAATGAGTACGATTAAAGTGCGACAGCAATTGGGGTGCAGTTCCATTATACGATTGTCTTAACAATGCCGAAAGACCACCGGGTCTGTGTTGACTAATCTAAACTCTGGATATCTTTTCTTTACTTTATCCTCTTTATCTTTTGCTAGTATTGGTCCTATATGCACATATTCTAAAAGCTTAGCGGGGAACTATCGTCATGCATTTTATCCACGTTTAATAACATCTCATCAGTGGGGTACTCCCGAGGCGGATCCCGTTTAGGAGCTCAGCACTACTCCGCCACCTTATTTATCTCATTGAAATTATTAATCTAAAAACGCCATAAAGATGTTGATCTTAAAGGATTGAACTCTATCCGAAAACAACATTCTAGAATGTTATCGTCATTATCATTACGATTCTAGTTTCAAAAACGATGACTCTCTTTTGAATCCTCGTAGTTTGTTGAGACAGAATAGCTATTTTGAAGAAAACTTTTGTAGTTCTTGAGAACATTCAGTCATAGAATATTCCTGGAAAATGCATCAGTATTACCAGAGGTCTTCATAATAATATTGTCATCTTTAAACATAATAGCCAGATGCTGATGCTGACTAATACATGATAAAGCCCAACAACCATTCTAAATGAATGACAGTTTCTACCACAACATTTTTCTTCATGCGAAAATTAAACACATAAAGATTTCTTTTGATCTATACTTAGACAAATAGTAGTATCTGTCCTAATGACATCAATTCATATTACAATCGACACTTACACATGATAACTAGAAAGTTTTACTAGGTATTTTTCAGATCAGGTTCTATATCTATGATGGCATCATTGTAGAAAACTACAAACACGATTTTACCCCTTGGACACAGGAAGATTAAACACAATGTTTCGGCTCCTCGGTAGAACACTGATGCACTTTGAGAGAGGGTATAATAGACTTTAAATGTTTACAAGATCCGCCAAGGCGGCAAAATACATTAACTCATCCGTCGAGGTCATTTATAGACACTTCTTCACTGAACTCTGAAAAATATGCACCTGGTAGCGCAGTTTATCGATTTTTATACGGATGCTCATTTTAAATTTTTTGTAAATTATTTAAGTTAAATGGCTGCAGAACAGCGTCGTTCTACAATTTTGACATAGTTTCAAATGCCATAATTGCAATCTGTATTGAATATATCTATTAATTCTGAATACATAAGTCCAAAGCTAAACAATTGTGCTGTGTCGGCATCGAAAAGTCGGTGATTAATGGTATCTACAATTGTTGCGAGTCAAATATAGAAATAATGGACAAGAGCAGCTATTAAAAATATTGGACAATCTTCGATGTCATTCGGCTCTATGTATGTAACACCACAGATTTCTGGGGCTATATAATTCGTTAAAACAGTTTACACTCATACTACCACATTCTTTATACATACAAGCCCACTATTCTAACCACGCTAAACACTTTATTAACCTGATTTTTATCTAACAAGTTGTGTATGCGGCAGAAATGGTAGAGTATCTGAAGACCAACTAGATTCATCAAATAAATCAATGTCTCAAGAGCTAGCAAGAATTGTGTGGAAATGAAATATGCTCTCATTAATCTGGTACAATATAGGATTTTGCCAATGATCATCAGTGAGCCTATTATAGTAGCTGTGGATTTCTGGTTAAAACCAACTCTGATTATTCTGCAGAAGTGGAAAGGCTAATGGAACTACCAGTTAAAACTGATATAGTGAATACCATGACTGTAGCCAGAAAGGTATTGATACTAGCAACAATATGAAACAGAATATATAAACCGGCAGAAAATAGAAGGAAATTGAAAAGGTAGAAAAATATTTACCAGAAGTTATATCTACAATTGCCAATAGTAATATAATAAAAAATAAAAATCTATCTTTCAGGCCAATATCAACGATAAACAGATCATGGAATGCTCTGAATGCTAGACACGAGTGAGAAATACTCTGGGATACTCGAAACTGATGGAACTGTGACTAGTCCATTGACGGGAAATAATACAATTACAACATTTATACCAATTTCTGCGTCCGATATGCAAAAGTTTACCATTTTAGAATATCTTTACATTATGAG

